# A site selection decision framework for effective kelp restoration

**DOI:** 10.1101/2024.08.08.607095

**Authors:** Anita Giraldo-Ospina, Tom Bell, Mark H. Carr, Jennifer E. Caselle

## Abstract

**Highlights:**

- Site selection is one of the most important factors for ecosystem restoration success
- A spatial prioritization framework for application to kelp restoration in California
- The framework merges kelp metrics derived from *in-situ* surveys and satellite imagery
- Site prioritization classification for every kelp forest site in California
- This framework can be applied to other species and regions with similar datasets

We present a decision support framework in the form of a spatially explicit site classification scheme to prioritize locations for conducting kelp restoration. The framework was created for the entire coast of California, where kelp has been lost and restoration projects are increasingly proposed, but the framework is broadly applicable to other coastal habitats or species that are being considered for restoration. We first created spatial distribution models using almost two decades of *in situ* kelp forest monitoring data and a comprehensive suite of environmental and biological variables, and used the outputs to evaluate the historical stability of kelp forests prior to a marine heatwave (MHW). We then used kelp canopy abundance data derived from satellite imagery to measure the impact of the MHW (i.e. extent of forest loss) and the recent state of kelp forests, including the trend of increase or decrease following the MHW. Finally, we integrated these site-specific kelp metrics to construct a classification tree for prioritizing restoration sites. Outputs of site prioritization are mapped across the study region, readily usable for managers and restoration practitioners with site-specific recommendations for restoration approaches. The framework can be updated due to knowledge of the important predictors of kelp and with new satellite imagery. Further, the framework can be adapted to other species and regions with similar data sets. This regional site selection framework is intended to be used in addition to socio-ecological, socio-economic, and administrative considerations.

## Introduction

Kelp forests are responsible for billions of dollars in ecosystem service provisions worldwide, underpinned by very high primary production, nutrient cycling and the creation of three-dimensional structure that supports a rich biodiversity (Eger et al., 2023; Reed et al., 2008). They provide critical habitat for species that comprise important fisheries including finfish, abalone and urchins, and are iconic marine habitats, culturally important and a major draw for tourism (Bennett et al., 2016; Eger et al., 2023). All these add to the innate value of kelp forests and their cultural significance for indigenous peoples and contemporary society (Eger et al., 2023; Thurstan et al., 2018). However, across the globe, many kelp forests have become increasingly threatened by multiple stressors that are exacerbated by climate change (Arafeh-Dalmau et al., 2021; Krumhansl et al., 2016; Wernberg et al., 2016). Globally, macroalgal cover has been in decline for the past 50 years (Krumhansl et al., 2016; Wernberg and Filbee-Dexter, 2019) due to factors such as marine heatwaves (Beas-Luna et al., 2020; McPherson et al., 2021; Wernberg et al., 2016), the decline of grazer predators with a subsequent increase in herbivory (Bosch et al., 2022; Rogers-Bennett and Catton, 2019), and the flourishing of new or invasive species of macroalgae (Félix-Loaiza et al., 2022; South et al., 2017). The loss of kelp forests can have significant impacts on biodiversity and associated ecosystem services they provide, whose economic value has been estimated to be between $500,000 and 1,000,000 USD per kilometer of coastline (Filbee-Dexter and Wernberg, 2018). Such widespread, and sometimes, dramatic loss of this iconic marine habitat represents a challenge for resource managers and conservation practitioners, since natural recovery may take years and it is hindered by increasing anthropogenic pressures (Bell et al., 2023).

While losses of marine habitats and ecosystem services can sometimes be counteracted by mitigating stressors, active restoration is increasing as an intervention strategy to recover terrestrial and marine ecosystems worldwide, including coastal marine systems (Perring et al., 2015; Saunders et al., 2020), with projects led across diverse groups such as universities, NGOs, businesses and local communities (Eger et al., 2024). Indeed, the United Nations has declared 2021-2030 as the Decade on Ecosystem Restoration, aligning with other global environmental protection challenges to be met by 2030 (e.g. 30×30; Target 3 of the Kunming-Montreal Global Biodiversity Framework). A key challenge in kelp restoration is its cost, which, depending on the intervention technique has been estimated at 1,000 to 1,000,000 USD per hectare (Eger et al., 2022b). The expense, combined with the ever-increasing spatial scale of kelp loss, compel the need for a framework that allows for scientifically informed decisions that increase the likelihood of successful restoration, while taking into account the effects of a changing climate (La Peyre et al., 2014; Zedler, 2007).

A major question driving ecosystem or species restoration success is that of where to restore (Bayraktarov et al., 2016; Eger et al., 2022b). The ultimate goal of site selection in kelp restoration is to identify sites where restoration actions are most likely to succeed and restored forests will persist (Eger et al., 2022b; Elsäßer et al., 2013; Gann et al., 2019). Selection of areas for restoration should be based on thorough analysis using the best possible information to attain the maximum benefit with limited investment instead of the often *ad hoc* allocation of funds for restoration projects. Prioritization of sites for restoration requires knowledge of historical distribution and abundance dynamics of species targeted for restoration because, in most cases, regions where species existed before their loss should be prioritized (Gann et al., 2019). Here, we define restoration success as the long-term persistence of a restored kelp forest.

The coast of California has experienced some of the most extreme declines of kelp forests documented around the world in the past decade. A marine heatwave in the Northeastern Pacific ocean that extended from 2014 to 2016 (Di Lorenzo and Mantua, 2016), combined with the widespread mortality of the sea star species *Pycnopodia helianthoides* (Hamilton et al., 2021), a key sea urchin predator, resulted in a decrease of over 90% of *Nereocystis luetkeana*, the dominant canopy-forming kelp in northern California (McPherson et al., 2021). This also resulted in the closure of the recreational red abalone fishery in 2018 and disaster declaration for the commercial red sea urchin fishery (Rogers-Bennett and Catton, 2019). Portions of central and southern California, as well as Baja California, Mexico, whose kelp forests are dominated by the giant kelp, *Macrocystis pyrifera,* also saw sharp declines, although the effect was less widespread (Beas-Luna et al., 2020; Smith et al., 2024). Importantly, these kelp forests have not recovered to pre-MHW conditions, and there is now increasing interest in assisting recovery of these ecosystems through active restoration.

In this study, we integrated the outputs from models of kelp distribution and abundance in California with remote sensing data and constructed a decision-making framework to identify locations with the highest potential for kelp restoration success. We modeled the two primary canopy-forming kelps in California, *Macrocystis pyrifera* and *Nereocystis luetkeana*.

Specifically, our objectives were to: 1) use the outputs of spatial models of kelp abundance and distribution to estimate historical stability of kelp at sites along the California coast, 2) use estimates of kelp abundance derived from remote sensing to calculate the amount of kelp lost following a large MHW (2014-16 NE Pacific MHW) and the current state and trends of kelp across California and 3) integrate the estimates of stability, loss and current state into a classification and prioritization framework. The ultimate goal is to enable resource managers and restoration practitioners to identify locations that are likely to benefit from active restoration interventions and those that are more likely to show natural regeneration (Gann et al., 2019). This framework can also be supported by the inclusion of socio-economic criteria and logistical considerations to further inform the optimal use of resources for ecological restoration.

## Methods

### Study area

This study encompassed the entire 1,350 km of coastal California, between the borders of Mexico to Oregon, including offshore islands (Figure 1). In California, there are two dominant kelp species that form a surface canopy (Carr and Reed, 2016). Bull kelp (*Nereocystis luetkeana*) is an annual species with high interannual variation in forest density and area (McPherson et al., 2021). Individuals are characterized by a single long stipe, up to 25 m in length that extends through the water column from the subtidal rocky reef, buoyed by a large pneumatocyst (Springer et al., 2010). In California, bull kelp is distributed from the Oregon border in the north to Point Conception in the south. North of Monterey Bay, central California, it is the dominant habitat-forming kelp, whereas in central California bull kelp usually grows in mixed forests with giant kelp (*Macrocystis pyrifera*). Giant kelp is a perennial species dominant in the temperate eastern Pacific and Southern Oceans (Schiel and Foster, 2015). In California, giant kelp ranges predominantly from Pigeon Point in the north to the border with Mexico in the south (Carr and Reed, 2016). Giant kelp abundance in California is very dynamic since individuals as well as entire forests are highly susceptible to dislodgement by ocean waves (Edwards and Estes, 2006; Graham, 1997).

**FIGURE 1.**
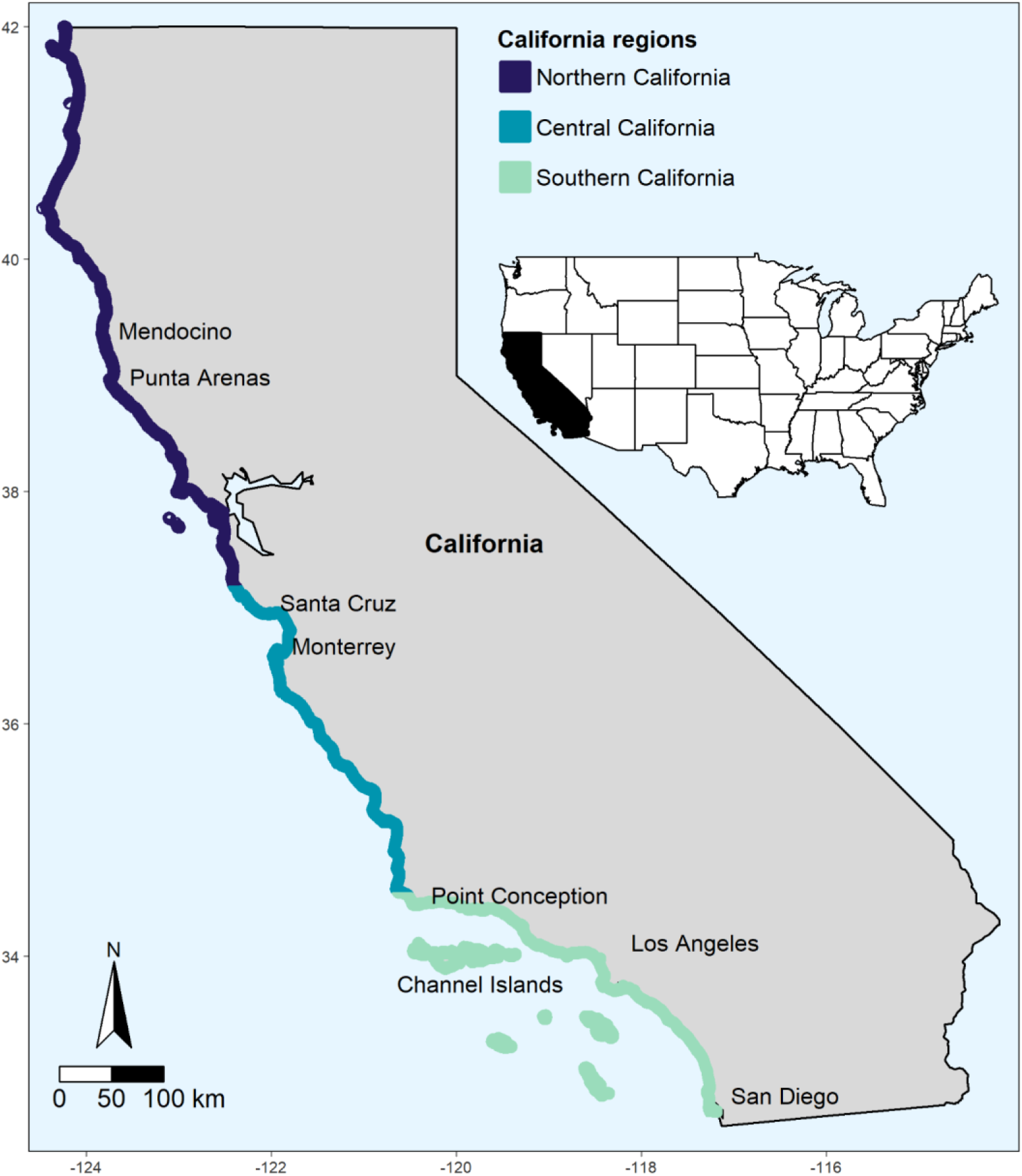
Map of California showing the three biogeographic regions: Northern, Central, and Southern, which includes the Channel Islands. Bull kelp is the dominant species in forests of the Northern region, while giant kelp dominates forests in the central and southern regions.

### Kelp metrics to incorporate in site classification framework

Three metrics of kelp dynamics formed the basis for the site classification framework. The first was temporal stability of kelp abundance prior to the NE Pacific MHW estimated from the historical maps obtained from spatial models of bull and giant kelp abundance (Giraldo-Ospina et al., 2024). The other two metrics were calculated from satellite-derived kelp surface canopy abundance and included an estimate of kelp lost after the NE Pacific MHW and the current proportion (percent) of historical kelp abundance. All three metrics were calculated for each site (cells of 300 x 300 m, the same resolution at which spatio-temporal maps of kelp were constructed with distribution models, Giraldo-Ospina et al., 2024) along the coast of California and integrated to classify each site into one of four restoration priority classes.

### Reconstruction of historical kelp density - in situ data

To construct maps of historical kelp density along the entire coast of California, we used spatio-temporal models of bull and giant kelp density. The dependent variables in the models (density of bull and giant kelp) were obtained from long-term *in situ* SCUBA monitoring surveys from the Partnership for Interdisciplinary Studies of Coastal Oceans (PISCO https://www.piscoweb.org/) and Reef Check (https://www.reefcheck.org/country/usa-california/). Sea urchin abundance, used as a predictor variable in the models, was also obtained from these *in situ* surveys. A suite of spatio-temporal data was obtained for variables thought to be associated with processes affecting bull and giant kelp densities. These variables included sea surface temperature, nitrate concentration, wave height, orbital velocity, net primary production, zoospore availability, and several descriptors of seafloor terrain (Giraldo-Ospina et al., 2024).

We modeled the density of each species (bull and giant kelp) separately using generalized additive mixed models (GAMs) (Wood, 2006) to investigate the relative contribution of variables in explaining spatial and temporal variation in density of bull and giant kelp. Annual maps of kelp density for each species were created by projecting the density predictions over the study region using the historical spatial data of predictors selected in the best models. All predictor variables were converted to 300 x 300 m resolution to produce a total of 18 annual maps for each species (from 2004 to 2021). See Giraldo-Ospina et al (2024) for additional details on model selection and evaluation.

### Calculation of kelp stability

Kelp stability was estimated using the time series of kelp density for the years prior to the MHW (2004 to 2013). Kelp stability was calculated for each cell (pixel) as the inverse of the coefficient of variation for each cell and scaled by the mean kelp density prior to the MWH (2004-2013), so that kelp beds with similar coefficients of variation would be ranked even higher if they had higher kelp densities.

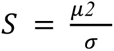

Where *S* is stability for each cell, μ is the mean kelp density estimated previous to the MHW 2004-2013, and σ is the standard deviation estimated previous to the MHW (2004-2013).

### Current Proportion and Kelp Loss-Satellite-derived data

We generated a time series of kelp canopy cover (bull kelp) and kelp canopy biomass (giant kelp) from remotely sensed imagery in order estimate the amount of kelp lost after the NE Pacific MHW and to estimate the current proportion of kelp compared to a baseline period. Kelp canopy area (m^2^) and biomass (wet weight in kg) were derived from Landsat 5, 7, 8, and 9 imagery and given for individual 30 x 30 m pixels (Bell et al., 2023; Bell et al., 2020; Bell et al., 2023). We extracted the maximum area for bull kelp in the northern region, and biomass for giant kelp in the central and southern regions, observed in each year to obtain the maximum area or biomass for each pixel per year. We then aggregated the data from 30 x 30 m pixels (Landsat resolution) into 300 x 300 m pixels (our ‘site’ resolution) by summing the total maximum canopy area or biomass.

### Current proportion of kelp compared to baseline

To create a historical ‘baseline’ of kelp abundance prior to the NE Pacific MHW, we averaged kelp abundance between 1985 and 2013 for every site pixel. We then calculated the current mean abundance of kelp for the most recent three years for which we had data (2020-2022) and used it to estimate the proportion of the historical baseline. Sometimes the current proportion of historical kelp was more than 100% indicating that in the last three years the mean kelp abundance was greater than the historical mean.

### Kelp loss post MHW disturbance (2014-2019)

We first estimated the lowest kelp abundance recorded between 2014-2019. Although the MHW was strongest during the years 2014 to 2016, kelp did not show a significant recovery during the years immediately following the MHW and 2019 was a hotter year than normal (McPherson et al., 2021; Smith et al., 2024). We then found the difference between the minimum kelp post-MHW and the historical mean of kelp abundance. In cases where there was a gain of kelp compared to historic baselines loss was described as zero.

### Classification of sites into restoration priorities

We integrated the metrics of pre-MHW stability (‘stability’), current proportion of historical kelp (‘current proportion of baseline’), and loss during and after the MHW (‘loss due to MHW’) into a three-dimensional space, where each metric constituted an axis (Figure 2). For each California region, sites (300 x 300 m pixels) were placed into a 3D space based on the logged values of the three metrics. We divided the sites into eight groupings, by finding the median of each logged metric to divide each axis in two parts resulting in eight sub-cubes (Figure 2).

**FIGURE 2.**
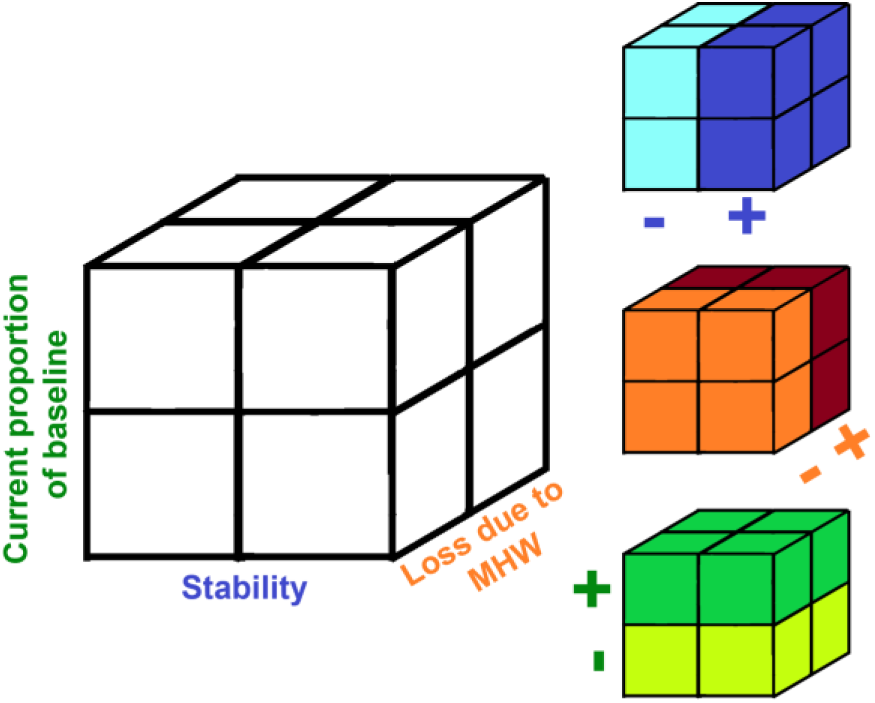
Three dimensional space formed by ‘stability’ in the x axis, ‘current proportion of baseline’ in the y axis, and ‘loss due to MHW’ in the z axis. The colored cubes show the characteristics of a site according to where it is located in the three dimensional space. For graphical purposes, we divided each axis by the median of the logged values of each metric to depict axes split in half. Sites to the left (light blue) showed lower stability than sites to the right (dark blue). Sites to the front of the cube showed low kelp loss (orange) compared to the ones at the back (dark red). Sites on the lower part of the cube currently have a lower proportion than their historical average abundance (light green) and the ones on top of the cube have a higher proportion than their historical average (dark green).

A hierarchical classification tree was then designed to classify sites according to each of the three metrics compared to other sites in the same region, so that each site is assigned one of four prioritization classes for restoration (Very low, Low, Mid, or High) (Figure 3). The first step in the classification tree is to separate the sites with higher historical stability from those with lower stability (Figure 3). The next step is to identify the magnitude of loss from the MHW at those sites. The final step in the decision tree is to evaluate the current state of kelp in each site with the metric of current proportion of kelp compared to a baseline. With this last question we can divide sites into four classes (Figure 3 and Table 1). Very low priority sites are historically unstable sites that, regardless of the effect of the MHW, currently have a lower proportion of kelp compared to their historical mean. We consider very low priority sites to be the most risky for an investment on restoration, as they historically have not sustained stable kelp densities and are currently in an unfavorable state for kelp, potentially requiring a large investment in restoration with uncertain outcomes (Figure 3). Low priority sites are those that, despite their lower historical stability, have a high proportion of kelp compared to their baseline, thus may also be a lower investment priority (Figure 3). Medium ‘Mid’ priority are historically stable sites that may or may have not experienced high losses of kelp after the MHW, but currently have a high proportion of kelp compared to their baseline (Figure 3). These sites are historically stable sites that are currently doing well in terms of their kelp abundance, so they are not in urgent need of an intervention but are considered mid priority for restoration and potentially high priority for other actions, such as monitoring, to assess a continued recovery trajectory (Figure 3). Finally, high priority sites are sites that were historically stable prior to the marine heatwave, and that may or may have not experienced high kelp losses after the MHW, but currently have lower proportion of kelp compared to their historical mean, and therefore are the ones that could benefit the most from restoration activities (Figure 3). Table 1 describes each of the resulting categories of prioritization and expands on a set of potential actions that could be employed.

**FIGURE 3.**
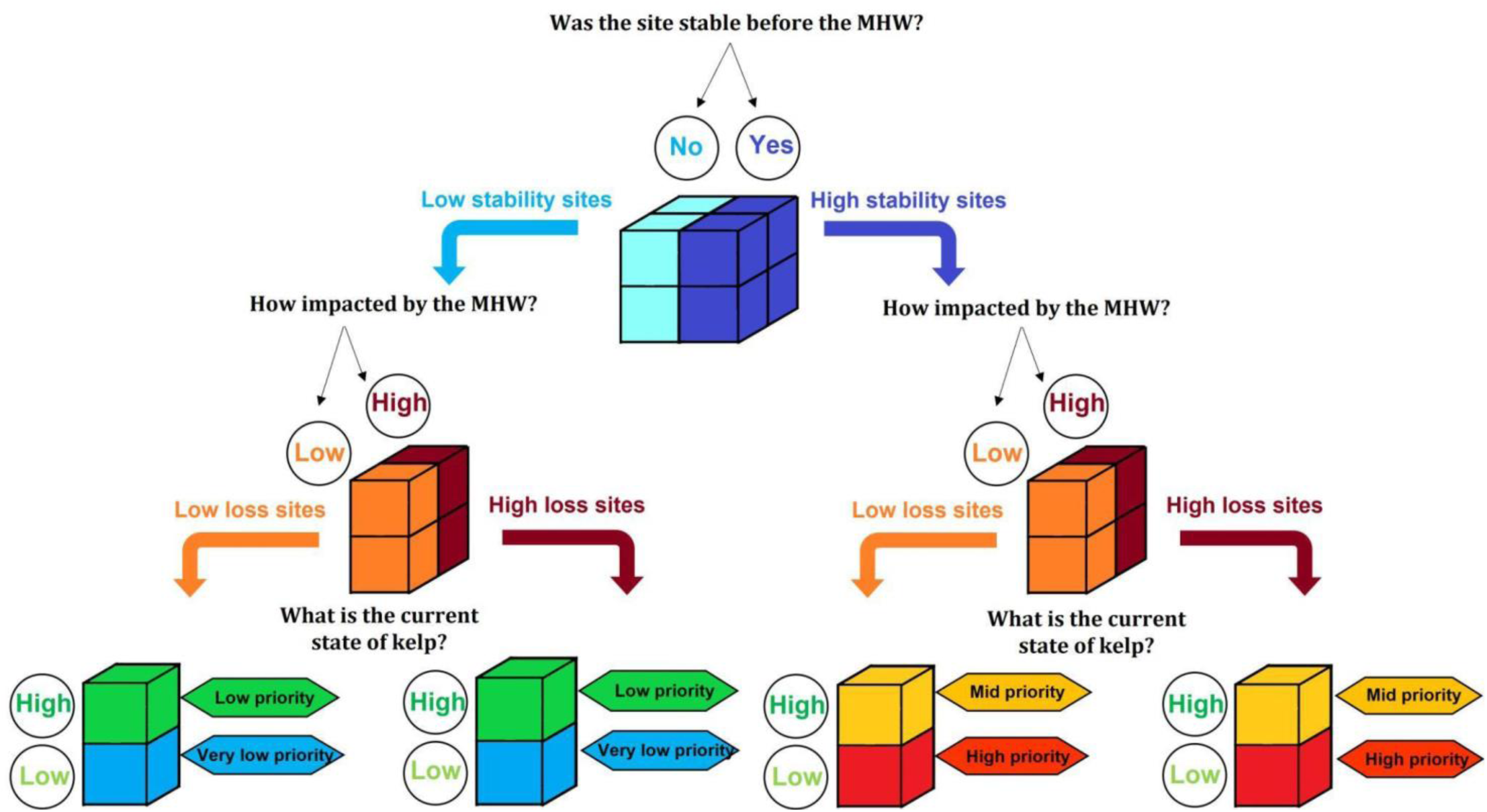
Classification tree to prioritize sites for kelp restoration activities in the state of California. The classification uses the values of the three metrics to assign sites into one of four prioritization classes: Very low (blue cubes), Low (green cubes), Mid (yellow cubes), High (red cubes). The prioritization takes into consideration the historical stability of kelp density prior to the NE Pacific MHW derived from modeled predictions of kelp density using the environmental predictors in combination with kelp loss after the NE Pacific MHW and current proportion of kelp compared to a historical baseline derived from Landsat imagery.

**Table 1.**
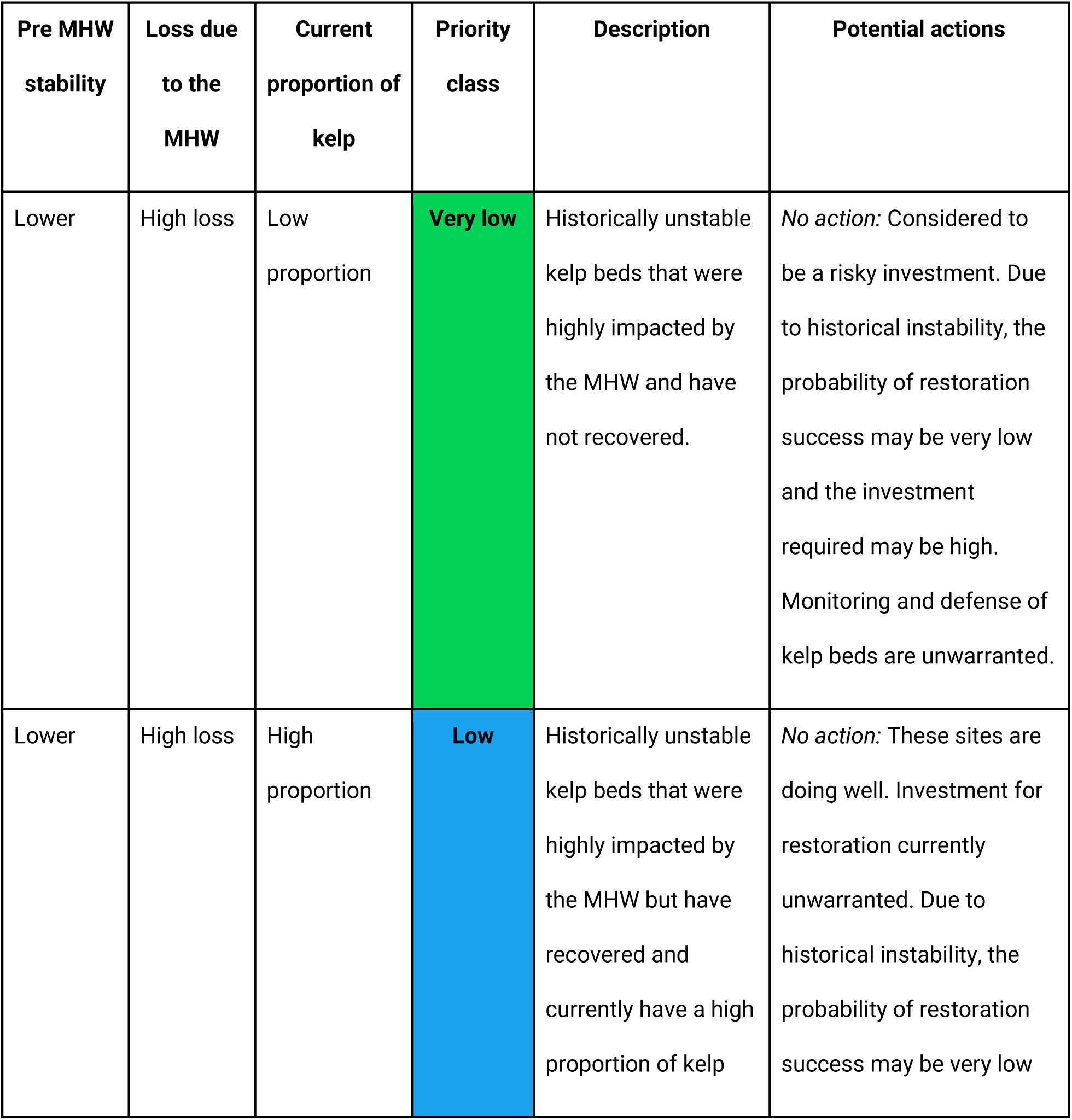
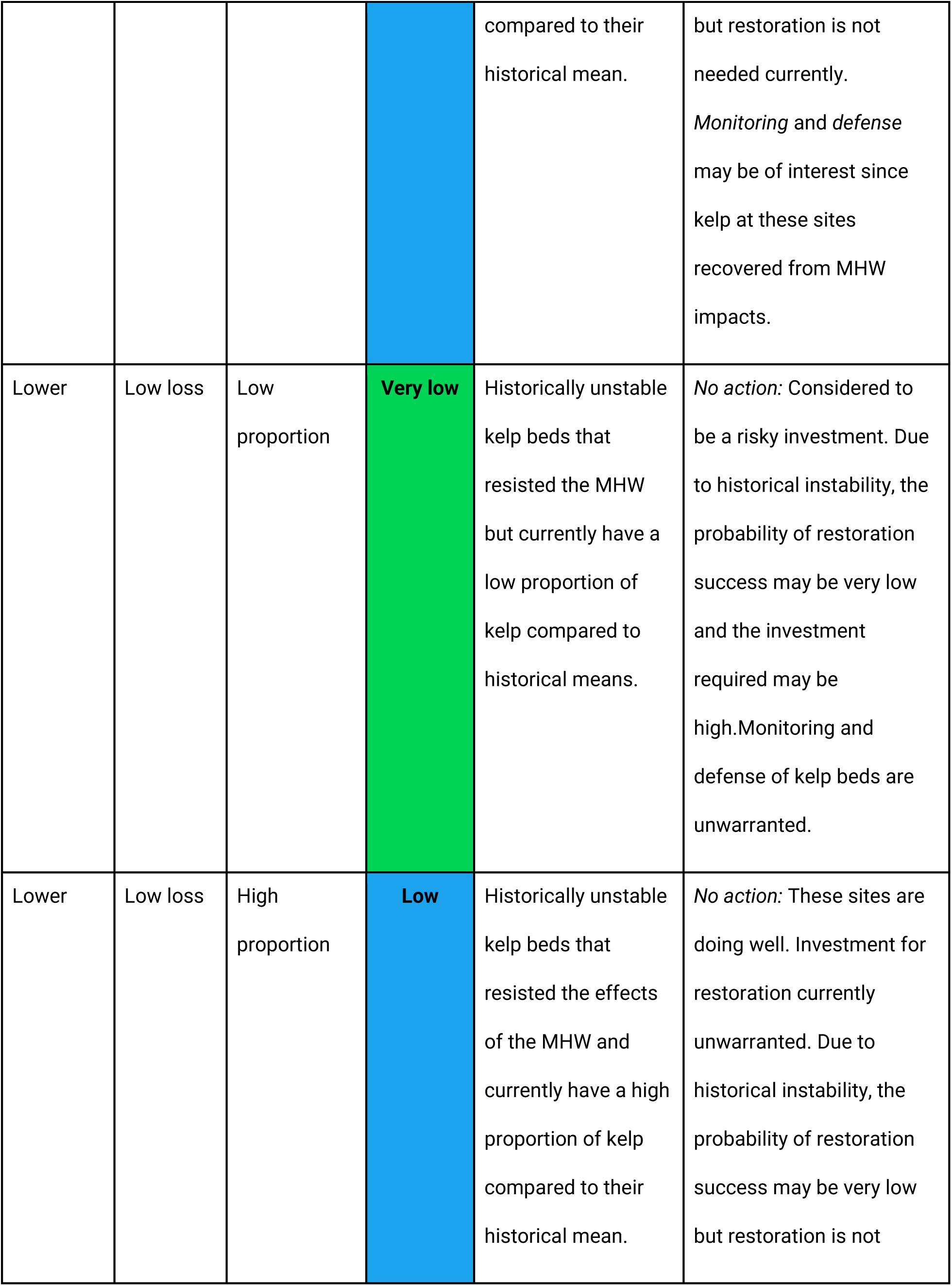
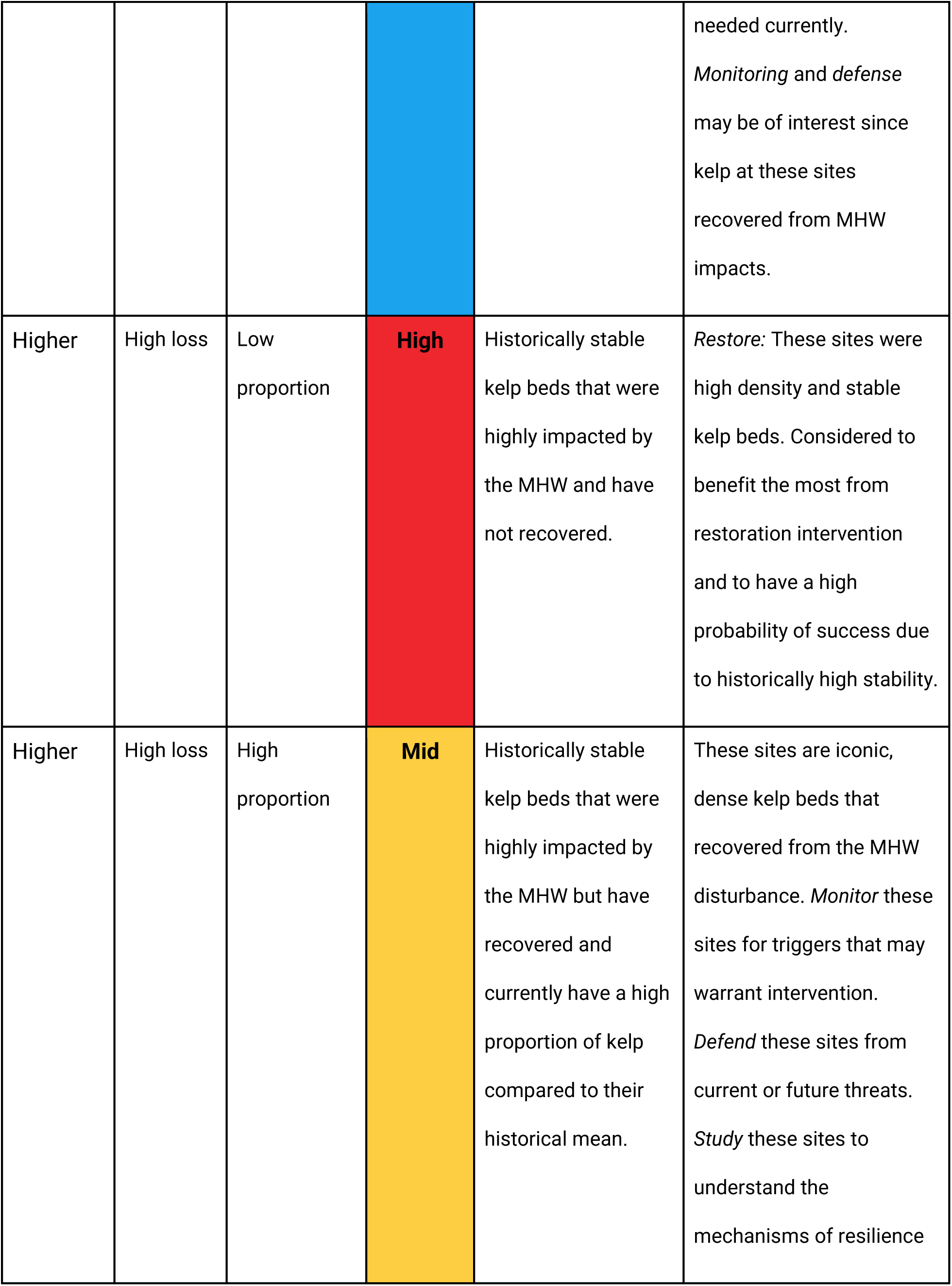
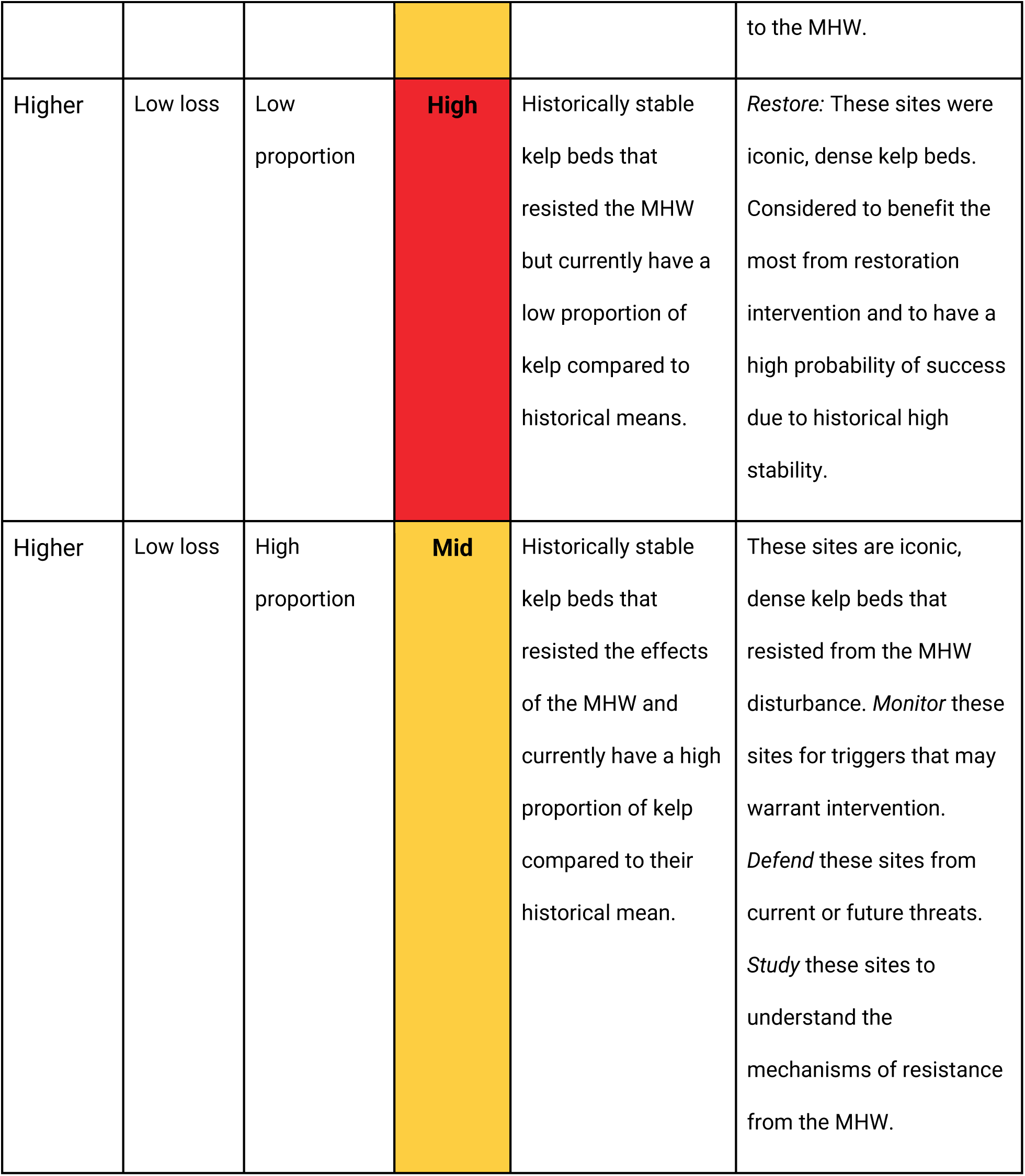
Description of the classifications resulting from the tree in Figure 3 (Very Low, Low, Mid and High). Color coding of priority classes corresponds to classes in Figure 3. Potential suggested actions for each class are described.

Finally, we estimated the recent trend of kelp abundance (increasing or decreasing; most recent five years). For this, we extracted mean kelp abundance data (area for the north coast and biomass for central and south coasts) from Landsat from 2018 to 2022. A simple linear model was fitted to the five values of kelp abundance for each pixel and classified as ‘increasing’ (positive slope) or ‘decreasing’ (negative slope). Sites with no slope (a slope of 0) were considered decreasing as they generally depicted sites with no kelp due to previous loss. These two categories of post-MHW abundance trend were used to further divide the four restoration priority classes into 8 categories to provide additional information on the recent conditions of kelp at each location (i.e. Very low-increasing, Very low-decreasing, Low-increasing, Low-decreasing, Mid-increasing, Mid-decreasing, High-increasing, and High-decreasing).

## Results

### Kelp metrics for classification scheme

Regions identified as high stability for bull kelp prior to the 2014-16 MHW were mostly located in the shallower parts of the north coast, and extending north and south from the coastline of Fort Bragg, Mendocino, and Point Arena (Figure 4a), indicating these regions have sustained dense kelp forests that experienced lower variation abundance before the MHW compared to other kelp forests in this region. The majority of sites (pixels) located in the deeper areas of the stability map for bull kelp showed very low stability. Most sites in the north region showed some loss of kelp after the MHW, however, the area between Fort Bragg and Fort Ross experienced the highest losses (Figure 4b). This area also has the lowest proportion of kelp compared to the historical mean kelp abundance in northern California, indicating it has not recovered from this disturbance (Figure 4c).

**FIGURE 4.**
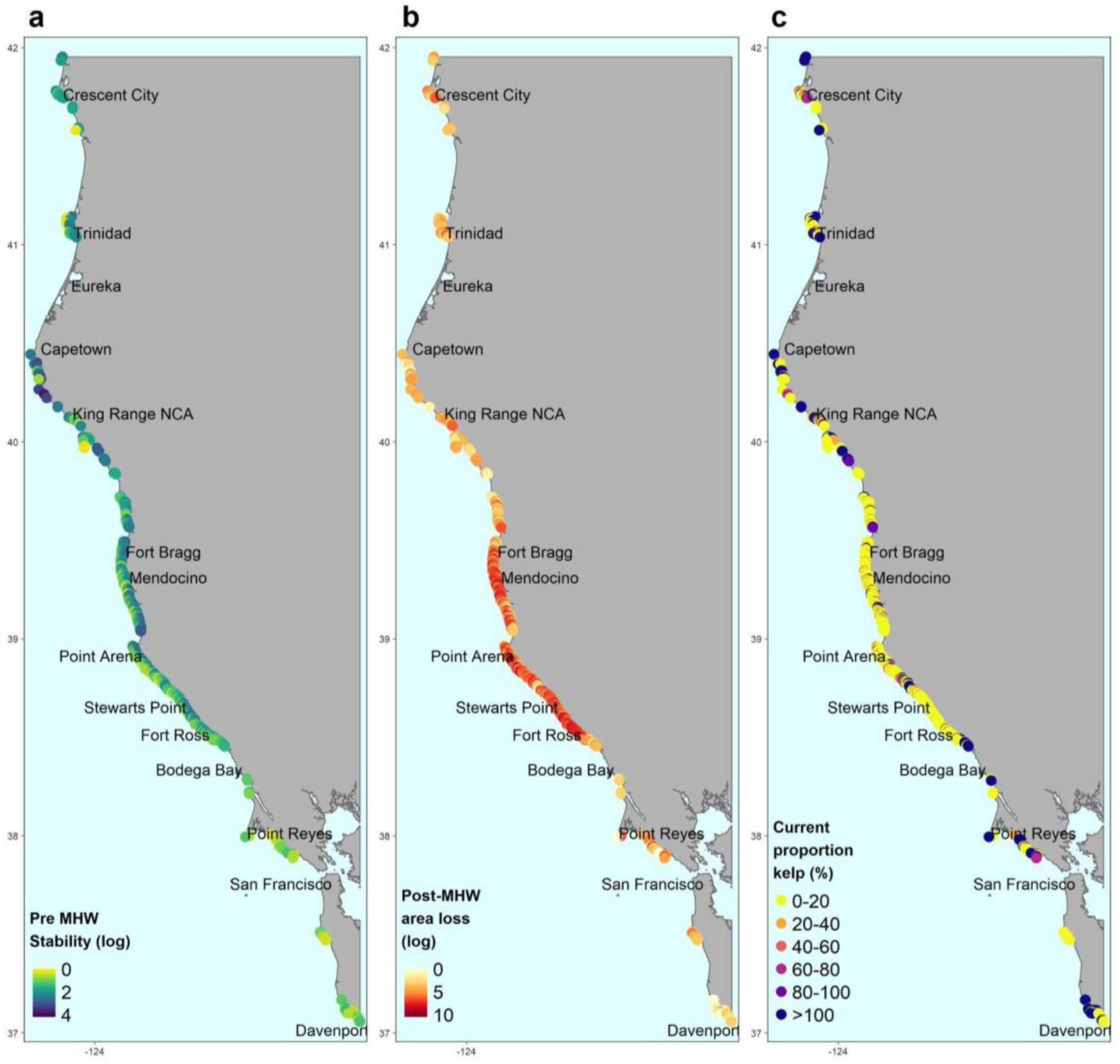
Maps of a) stability (log scale; years), b) kelp loss after 2014-16 marine heatwave (log scale; area), and c) current proportion (%) of kelp compared to baseline of bull kelp in the north coast.

Giant kelp stability in the central coast was generally high with the highest stability sites along the north of Monterey Bay and from the Monterey peninsula to the Big Sur coastline (Figure 5a). Kelp loss after the MHW (compared to historical mean) was high across the region with lower losses north of Monterey Bay and San Luis Obispo (Figure 5b). The current proportion of kelp compared to the historical mean varied along the central coast. Relative to other locations, recent kelp cover remained particularly low at several locations along Santa Cruz, around and south of the Monterey peninsula, Big Sur, and San Luis Obispo (Figure 5c). Other locations between San Luis Obispo and Big Sur, and north of Point Conception had more kelp than their historical mean (Figure 5c).

**FIGURE 5.**
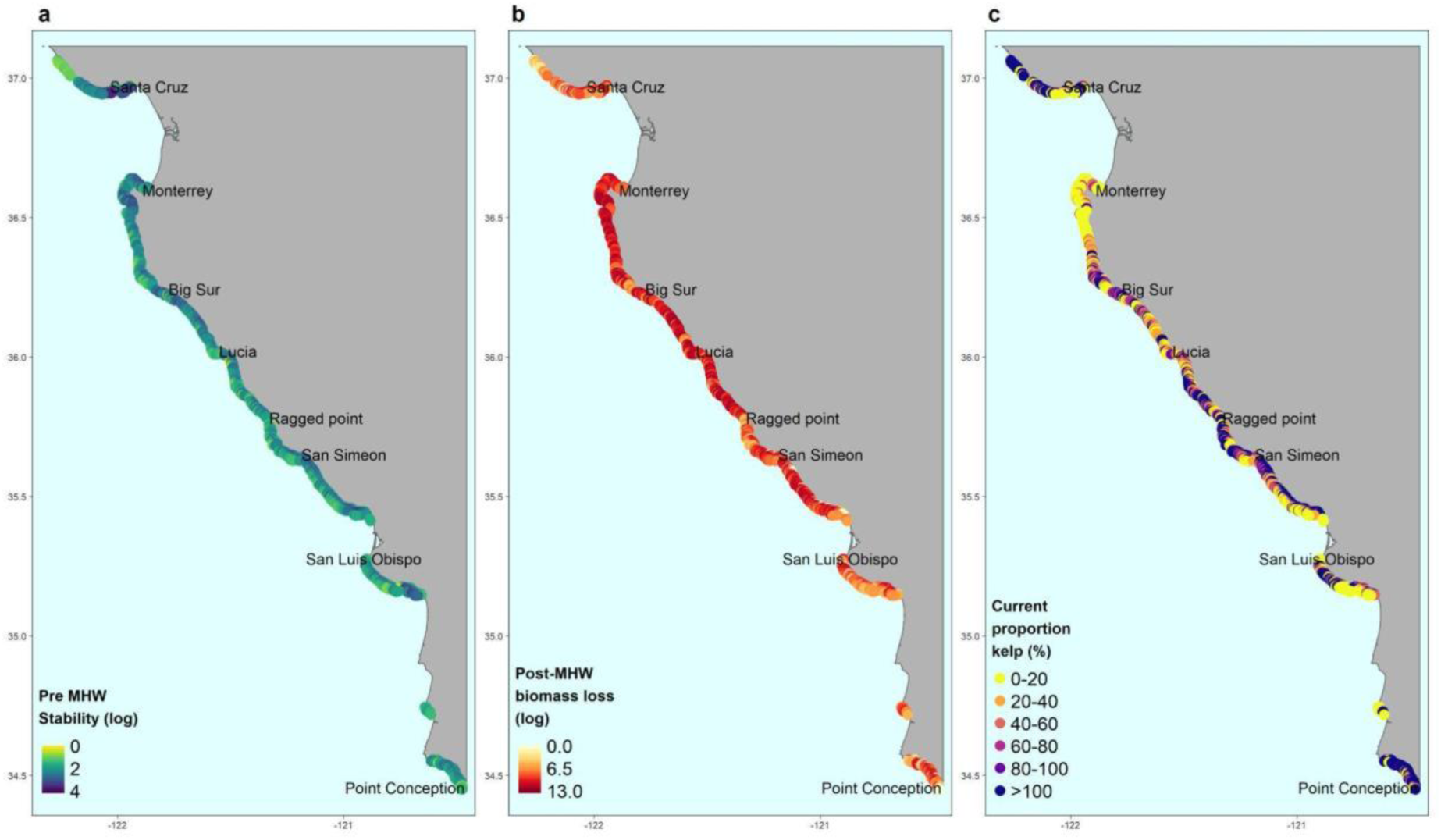
Maps of a) stability (log scale; years), b) kelp biomass loss after 2014-16 marine heatwave (log scale; biomass), and c) current proportion (%) of kelp compared to baseline of giant kelp in the central coast.

The south coast and islands showed a more patchy distribution of high stability sites for giant kelp, compared to the north and central regions. High stability of kelp was observed at all the island sites, and some mainland sites like Palos Verdes and San Diego (Figure 6a). All other areas showed low historical stability with the very low stability sites located along the mainland coast (Figure 6a). Loss of kelp biomass after the MHW was widespread across the region, with the highest losses observed around Santa Barbara, San Diego and the Channel Islands (Figure 6b). The current proportion of kelp compared to the historical mean in the region was less than 20% for several locations that showed high stability previous to the marine heatwave, such as San Miguel and Santa Rosa Islands, and San Diego indicating that these previously stable sites, have not recovered from the NE Pacific MHW (Figure 6c).

**FIGURE 6.**
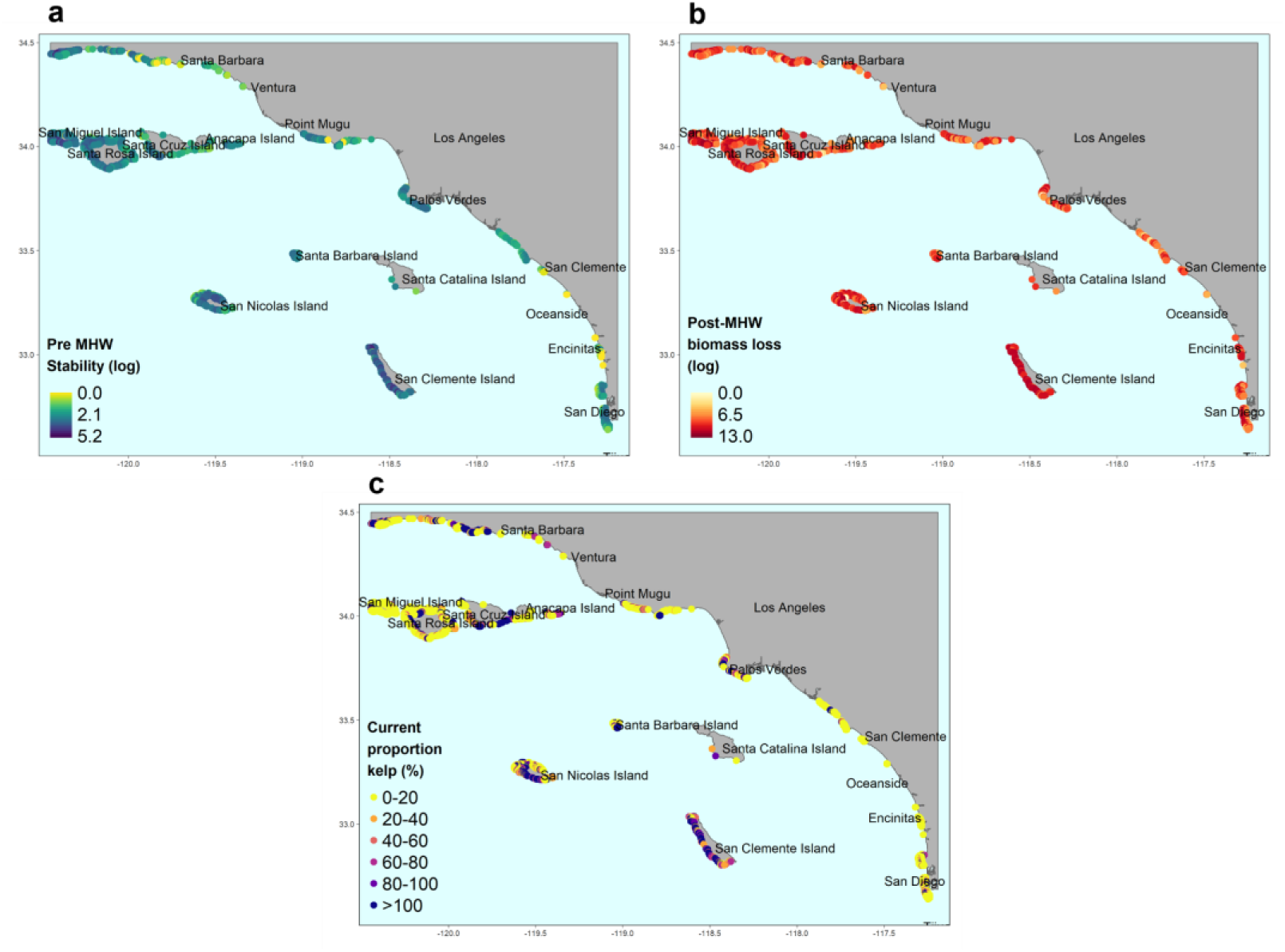
Maps of a) stability (log scale; years), b) kelp biomass loss after 2014-16 marine heatwave (log scale; biomass), and c) current proportion (%) of kelp compared to baseline of giant kelp in the south coast.

### Results of site classification scheme

In the north coast, the shallower portions of the coastline from Fort Ross to Fort Bragg presented the most sites which were classified as high priority for bull kelp restoration, while deeper sites were classified as low or very low priority (Figure 7a). Sites north of Fort Bragg were generally classified as a mix of high and mid priority sites. South of Fort Ross all sites were classified as low or very low priority, reflecting their lower stability compared to others in the region (Figure 7a). In the central coast, several regions like the Monterey peninsula had the most sites classified as high priority for giant kelp restoration, indicating these were sites with higher stability compared to others in the region, and which currently exhibit lower proportions of historical kelp densities (Figure 7b). Sites in the south coast classified as high priority for giant kelp restoration are visibly clustered around San Miguel and Santa Rosa Islands, while other high priority sites were located west of Santa Barbara, and in the San Diego region near La Jolla and Point Loma (Figure 7c). See close-up maps of site classification in Appendix A for visual identification of site-specific restoration categories.

**FIGURE 7.**
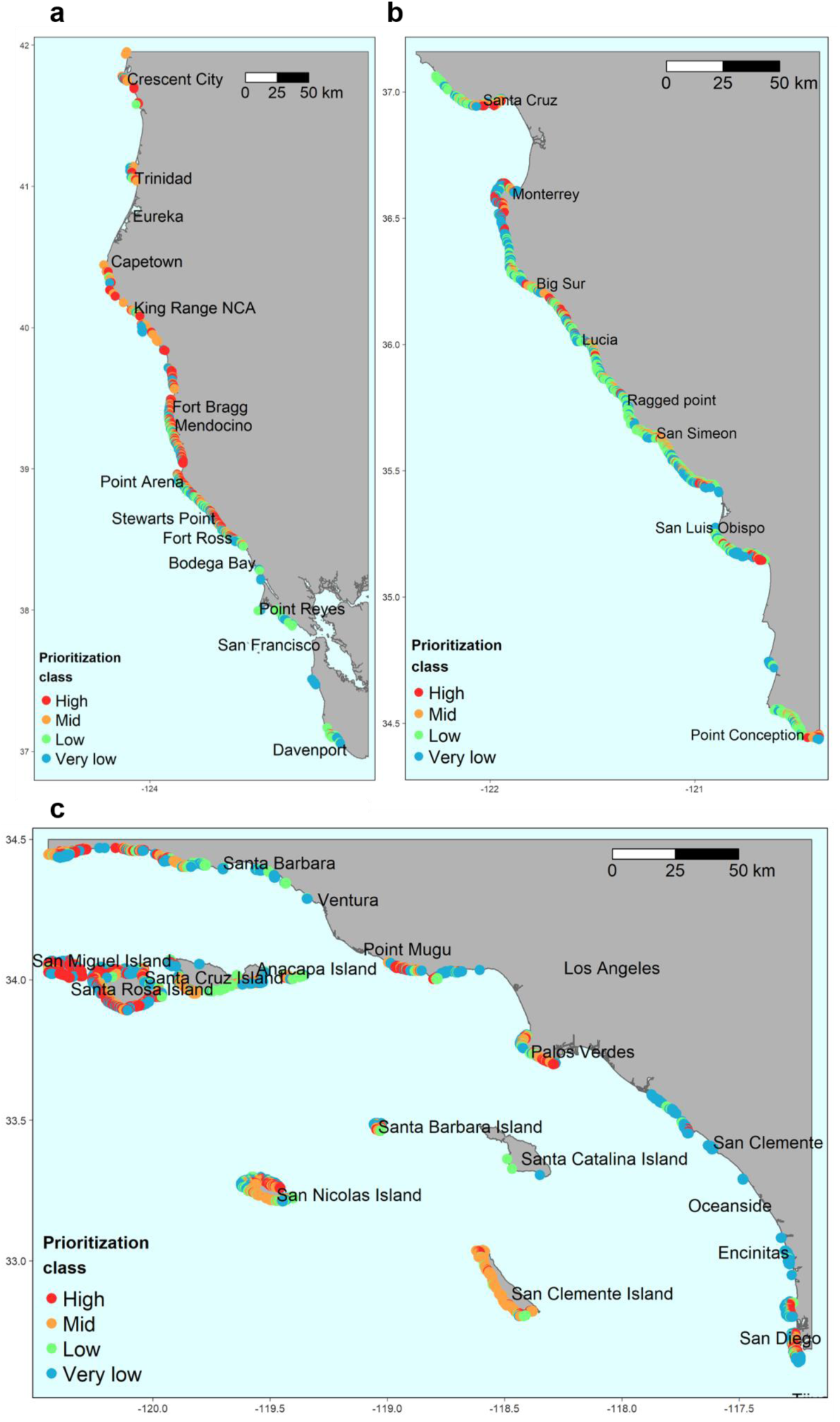
Maps of restoration priority classes for a) bull kelp in the north coast, b) giant kelp in the central coast, and c) giant kelp in the south coast.

### Kelp restoration classes and protection status

Approximately a quarter of all kelp sites in California fell into each of the four main restoration priority classes, a consequence of the choice to split categories at the median values of the metrics (Figure 8a). However, regionally, we see differences in number sites falling into different prioritization levels. The south region had the highest proportion of sites with high and mid priority for kelp restoration, followed by the north region. As a simple example of how one could layer other factors onto the classification scheme, we calculated the proportion of sites currently located in Marine Protected Areas in California for each classification. For sites located inside MPAs, 19% were categorized as high priority and 25% as mid priority for restoration across the state (Figure 8b). The north region had the highest proportion of high priority sites located in MPAs, followed by the south region (Figure 8b). The result of recent trend of kelp abundance observed in the north coast over the past five years was of ‘no change’ for most sites (Appendix B, Figure B1). Sites with an increasing trend were mostly located south of Mendocino, and sites with a decreasing trend were mostly located north of Mendocino. Most sites in the central coast showed a decreasing trend (Appendix B, Figure B1). In the south coast, areas west of Santa Barbara, the northern Channel Islands, and San Diego contained multiple sites with decreasing kelp abundance in the past five years (Appendix B, Figure B1).

**FIGURE 8.**
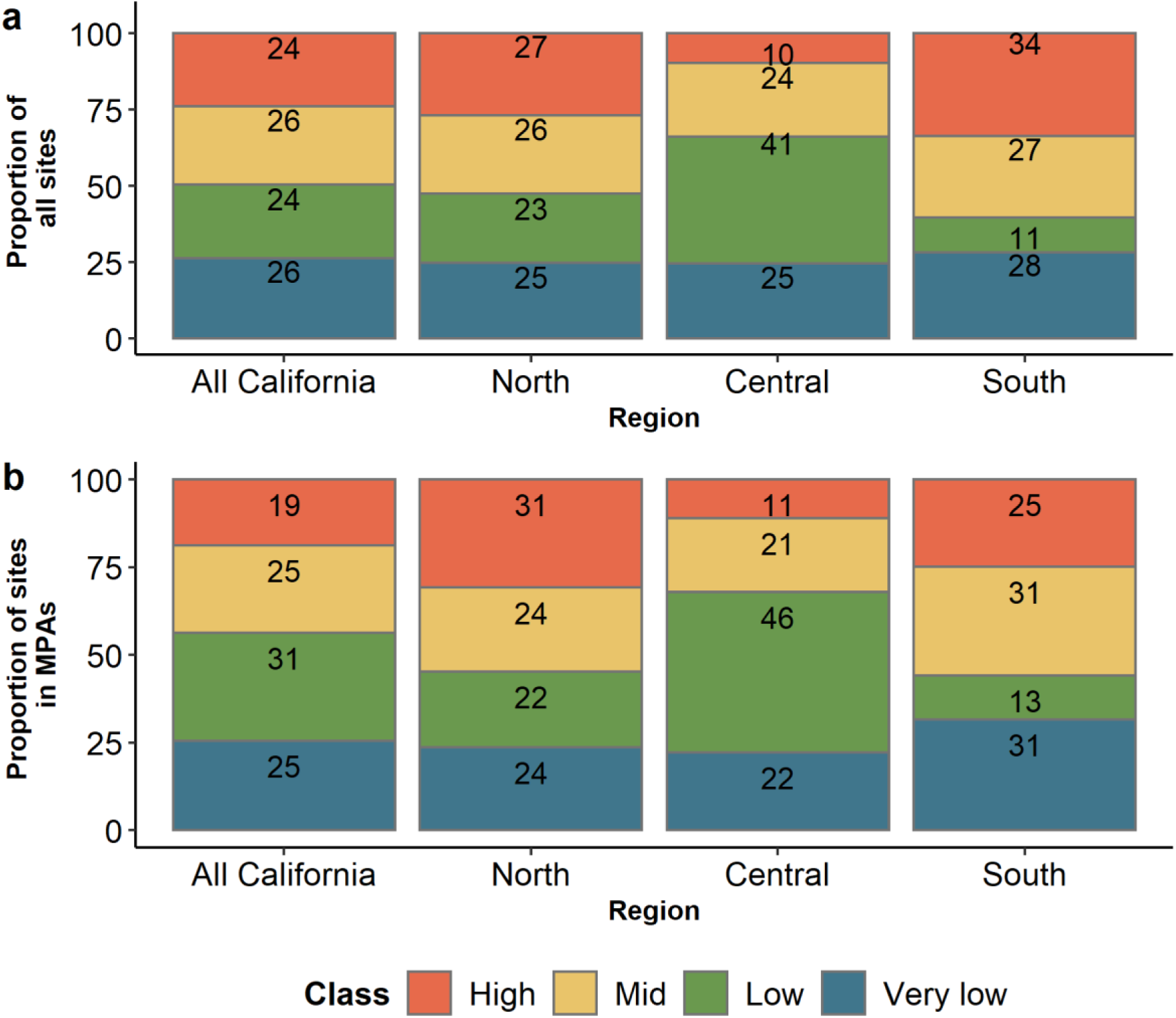
a) Proportion of sites within each restoration priority class in all California and for each region. b) Proportion of sites within MPAs for each restoration priority class in for all California MPAs and in each region of the state.

## Discussion

Globally, kelp forests are increasingly threatened by a wide variety of stressors, including climate change, directly diminishing the biodiversity they sustain and the ecosystem services they provide. Restoration of kelp forests has been increasingly used as an intervention to mitigate ecosystem degradation (Eger et al., 2022a). The consideration of site selection has been found to be more important for marine ecosystem restoration success than the magnitude of financial investment (Bayraktarov et al., 2016). Here, we created an ecologically-focused, spatially-explicit site classification framework to help managers and restoration practitioners prioritize among potential sites for restoration. The framework uses the best available ecological datasets to enable managers and others to consider, identify, and weigh the predicted abundance, stability and persistence of restored forests among alternative restoration sites. The decision framework was designed for the two canopy-forming species in California, giant kelp (*Macrocystis pyrifera*) and bull kelp (*Nereocystis luetkeana*). By including both species, the outputs of this framework are spatially scalable from local to regional, to statewide decision processes. Together with the many other considerations required to inform kelp forest restoration decisions (e.g. community support and input, fisheries consequences, logistical constraints, funding availability; Gleason et al., 2021), this knowledge can inform the relative values of where, when and how restoration might be pursued at potential or proposed restoration sites. The framework can also help practitioners better understand and contextualize the results - successes and failures - of ongoing restoration projects that were placed without consideration of ecological and environmental conditions. Most importantly, by emphasizing the role of forest stability, restoration can be prioritized at those sites where restored forests are more likely to persist longer into the future.

By considering post-MHW trends and current forest state relative to pre-MHW forest states, differences in potential enhancement (i.e. increased forest area and abundance) can be weighed among sites. This study combined *in situ* diver surveys of kelp forest communities allowing for co-located and simultaneously captured data of urchin and kelp densities. When combined with remotely sensed kelp canopy abundance the site classification scheme can be updated in almost real time, by estimating the metric of current proportion of kelp with the most up-to-date kelp imagery available (Cavanaugh et al., 2023). This means of revising the prioritization classes annually is key for the two species with high natural variability, like bull and giant kelp (McPherson et al., 2021; Rodriguez et al., 2013) and aligns with the decision-making timelines faced by restoration practitioners.

### The site prioritization scheme

Methods for ranking sites are common in conservation planning (Klein et al., 2010; Leslie, 2005), but are now being applied to ecological restoration (Eger, 2020). The priority scheme enabled us to suggest alternative restoration actions, which include no action, watch/monitor, defend extant patches, or restore (Table 1). This result enables those interested in forest restoration to consider a broader range of actions, tailor actions to the history and state of a forest, and further prioritize intervention where it could be most cost-effective. Restoration of historically unstable forests to their pre-MHW levels is less likely to persist into the future, suggesting that restoration might best be pursued elsewhere. High or mid priority forests that are exhibiting a trajectory of recovery may warrant less investment than high priority forests that exhibit no trend of recovery. Instead of active restoration interventions, high and mid priority forests that are exhibiting a trajectory of recovery may benefit from monitoring to ensure they remain on a positive trajectory and consider intervention if that changes. In our framework, forests that were historically stable but experienced high losses and have yet to recover are more likely to exhibit greater and more durable benefits from restoration. Nonetheless, the ultimate decision on where to restore will depend on the specific objectives of each restoration project and many other considerations that may include community support, fisheries consequences, logistical constraints, funding availability.

Site prioritization schemes require knowledge about organism’s distribution and spatial variability in abundance (Johnston et al., 2015). Occurrence or persistence data is frequently used for site prioritization in marine ecosystem restoration (Elsäßer et al., 2013; Johnston et al., 2015). However, presence and abundance may display different patterns of spatial and temporal variation (Gaston and He, 2011; Oliver et al., 2012). Our classification scheme benefits from access to spatially explicit abundance estimates and historical stability of kelp. The high and mid priority classes always include stable forests, while the very low and low priority classes always include unstable forests. The emphasis on stability ensures that kelp restoration is prioritized in sites where restored kelp forests are more likely to be persistent and abundant into the future, hence applying resources where they can have the greatest benefits. We recognize that many locations will not have access to the wealth of data that exists in California but suggest that the concepts of the framework will translate well to other forms of information on stability including community, traditional and indigenous knowledge.

Notably, our framework identifies the relative, not absolute, importance for restoration across the array of conditions that are observed at the time of the site classification. If this framework is applied in a period when forests across all sites have high abundance of kelp, the classification would still result in some sites being classified as ‘high’ priority sites relative to others. For that reason, further evaluation of ‘high’ priority sites is needed to confirm they warrant restoration, or if other actions are more appropriate, such as conservation. The framework assumes that functional relationships between abundance and the key drivers will remain similar into the future. If true, then the models used in this framework should accurately reproduce kelp dynamics across the state into the future. If these functional relationships change, for example with changing climate, new models and projections will be needed.

### Incorporating other considerations into decision making

There are many other considerations to the design and implementation of kelp restoration projects. The dynamic nature of kelp ecosystems, complex and regionally specific drivers of kelp loss, and predicted climate-related changes for California waters make for a complicated decision context for knowing when, where and how to intervene to maintain or actively restore kelp forest ecosystems. This framework used ecological and environmental models to inform multiple aspects of those decisions. The spatial prioritization scheme created here can inform multiple steps of a structured decision making (SDM) process when combined with additional information such as logistics (Puckett et al., 2018, Gleason et al., 2021), socio-economic factors (Gouezo et al., 2021) and legal constraints such as permitting.

For example, choosing kelp restoration sites within marine protected areas (MPAs) may improve survival and kelp recruitment due to the increased protection from other stressors (Cebrian et al., 2021) and result in additional benefits (e.g., enhanced fish stocks) (Hopf et al., 2022) yet in many locations, including California, restoration in protected areas is not currently allowed ( (Filbee-Dexter et al., 2024).). In our study, up to a quarter of the total kelp sites classified as high priority were within an MPA, highlighting the need to review MPA management plans regularly to ensure they adapt to climate change.

Although we use the term ‘prioritization’ for simplicity, there may be other (non-ecological) ways for stakeholders to prioritize locations to conduct restoration, and these will depend on the goals of a project. For example, community involvement may be a major goal of a kelp restoration project, and might be weighted equally with likelihood of long-term kelp recovery. Incorporating the outputs from this framework into broader decisions regarding kelp restoration may increase the probability of restoration success. Notably, this framework could be replicated to other geographies, other coastal habitats or species and can be adapted for other forms of data and knowledge.

## Acknowledgements

We thank the numerous individuals who contributed to data collection as part of the long-term kelp forest monitoring programs of Partnership for Interdisciplinary Studies of Coastal Oceans (PISCO) and Reef Check California. We thank K. Elsmore from the California Department of Fish and Wildlife and M. Esgro from California Ocean Protection Council for their continuous feedback during this project. We also thank D. Malone, A. Parsons-Field for assistance with data curation and J. Freiwald for assistance with Reef Check data acquisition. This research was funded by California Sea Grant (R/HCEOPC-18) as part of the California state-wide Kelp Recovery Research Program in collaboration with the California Ocean Protection Council. We also wish to thank M. Yeager, the Caselle lab members and Passionate Women in Science (PWIS) team for the fruitful discussions and support. This is Publication number XXX from PISCO.

## CRediT

***Anita Giraldo-Ospina:*** Conceptualization, Formal analysis, Methodology, Visualization, Writing - original draft, Writing - review and editing. **Tom Bell:** Conceptualization, Funding acquisition, Methodology, Data curation, Writing - review and editing. **Mark Carr:** Conceptualization, Funding acquisition, Methodology, Writing - review and editing. **Jennifer Caselle:** Project administration, Conceptualization, Funding acquisition, Methodology, Writing - review and editing.

## Data availability

Data are available in DataOne at doi:10.25494/P6/When_where_and_how_kelp_restoration_guidebook_2.

