## Appendix B for "A site selection decision framework for effective kelp restoration"

**Appendix B**. Maps of site classification including both their restoration priority and the trend of kelp abundance.

### *Results of recent trends*

The trend of kelp abundance observed in the north coast over the past five years was of ‘no change’ for most sites (Figure B1-a). Sites with an increasing trend were mostly located south of Mendocino, and sites with a decreasing trend were mostly located north of Mendocino. Most sites with ‘no change’ in the past five years were sites with zero or very low kelp abundances, which is why for the prioritization classes with trend, they were grouped with the ‘decreasing’ trend sites. Most sites in the central coast showed a decreasing trend, although notable clusters of sites showing increases in kelp abundances were located around Santa Cruz, Monterey Bay, Carmel Bay, north of Big Sur, north of Ragged Point and at San Luis Obispo (Figure B1-b). In the south coast, areas west of Santa Barbara, the northern Channel Islands (San Miguel, Santa Rosa, Santa Cruz and Anacapa), Point Mugu, Palos Verdes, San Nicolas Island and San Diego contained multiple sites with a decreasing kelp abundance in the past five years (Figure B1-c). Increasing kelp abundance was only observed for sites around Santa Barbara, south of Santa Rosa Island, Palos Verdes and the southern Channel Islands (San Nicolas, Santa Barbara, Santa Catalina and San Clemente).

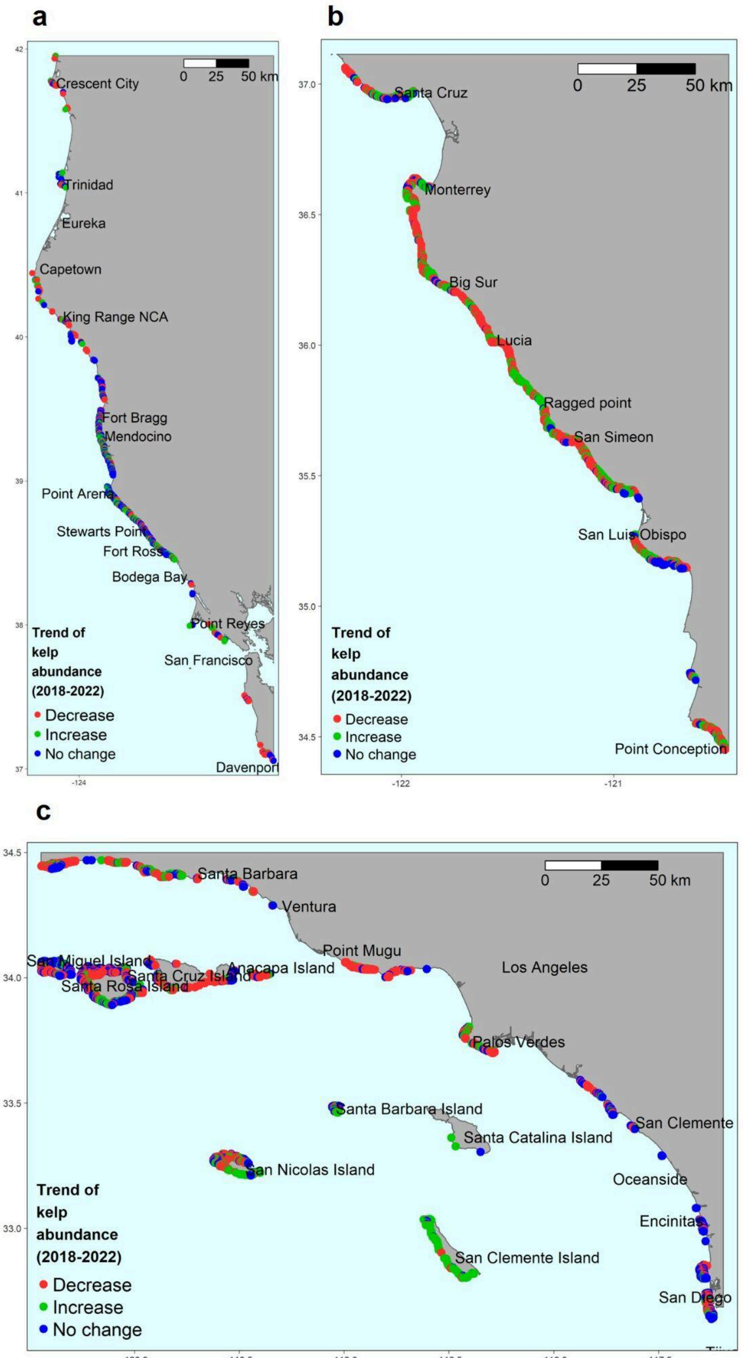


**FIGURE B1.** Trend of kelp abundance in the recent five year period (2018-2022). No change often indicated sites with zero kelp over the past five years. a) bull kelp in the north coast, b) giant kelp in the central coast, and c) giant kelp in the south coast.

***Site classification with the recent trend of kelp abundance***

The combination of the prioritization classes and the recent five-year trend of kelp abundance for each site resulted in eight categories of prioritization for kelp restoration, where the prioritization class is given the most recent trend of kelp abundance (either increasing or decreasing). The initial colors for prioritization categories were preserved (from Figure 7 in the main manuscript), and the intensity of each color varied to identify if the site had shown an increase or decrease in kelp abundance (decreasing or no change) over the past five years, with more intense colors illustrating a decreasing trend and lighter colors illustrating an increasing trend (Figure B2).

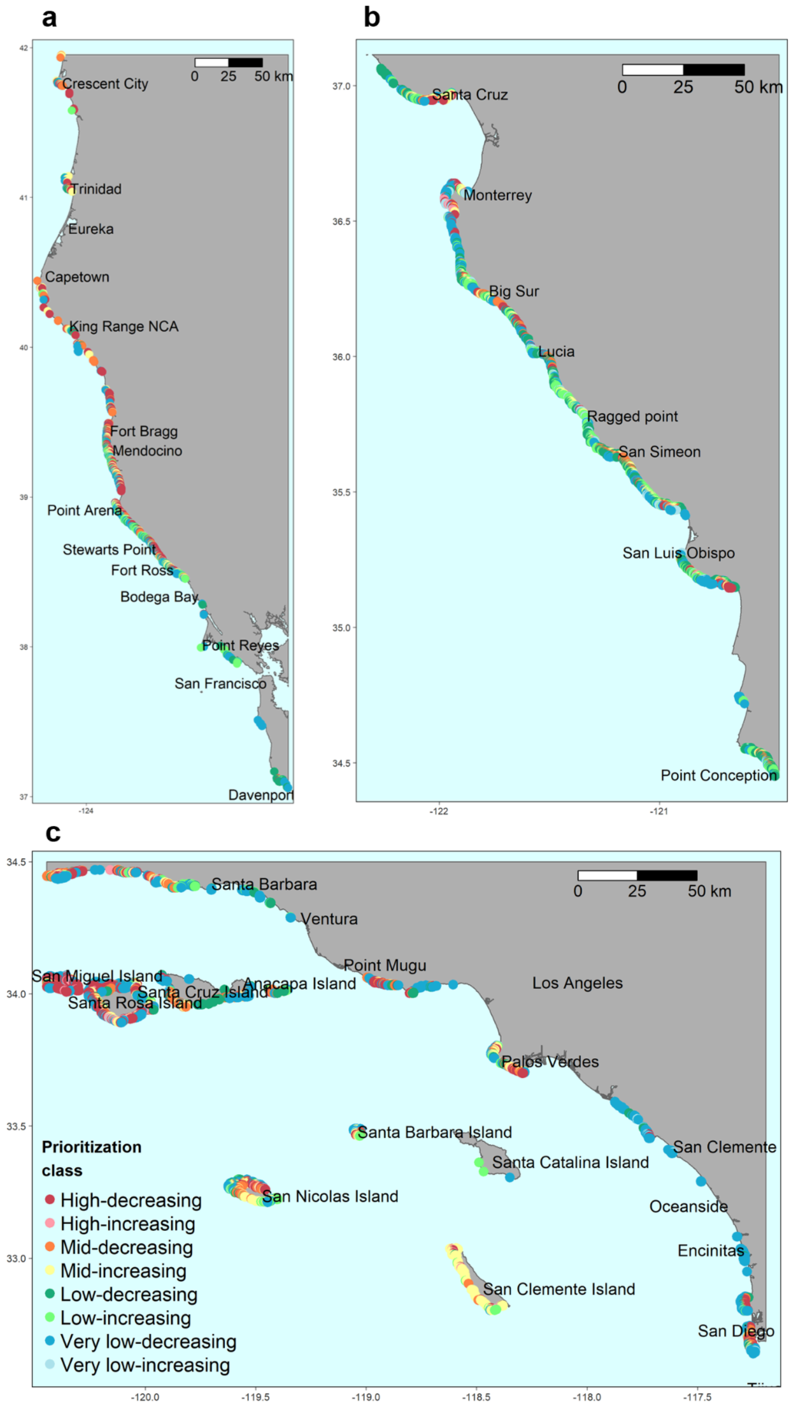


**FIGURE B2.** Maps of site classification including both their restoration priority and the trend of kelp abundance over the recent five years for a) bull kelp in the north coast, b) giant kelp in the central coast, and c) giant kelp in the south coast.
