## Appendix A for "A site selection decision framework for effective kelp restoration"

**Appendix A**. Close-up maps of kelp restoration classification for California.

### NC classification_map 1

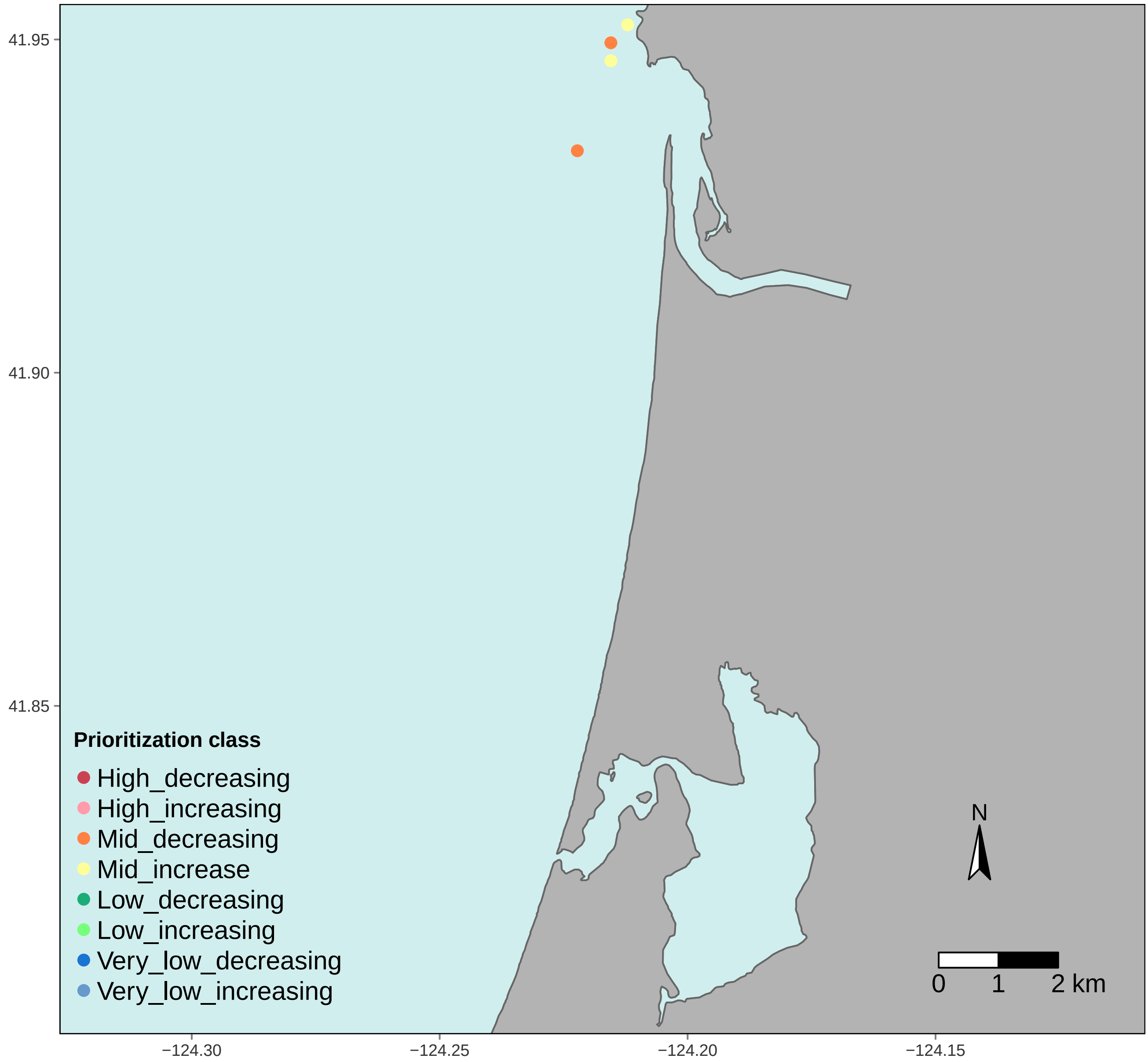

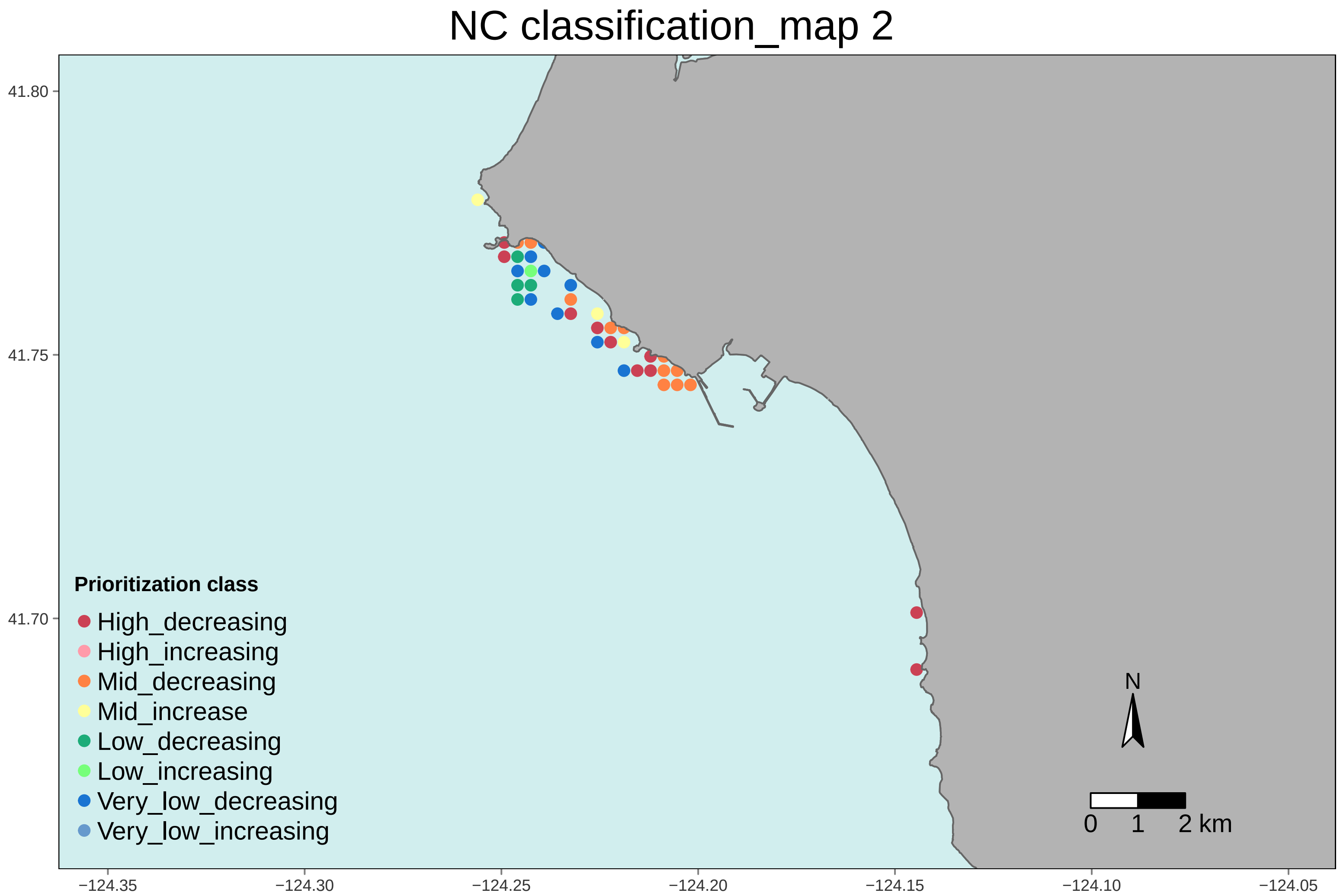
NC classification_map 2
41.80
41.75
41.70
−124.35
−124.30
−124.25
−124.20
−124.15
−124.10
−124.05
Prioritization class
High_decreasing
High_increasing
Mid_decreasing
Mid_increase
Low_decreasing
Low_increasing
Very_low_decreasing
Very_low_increasing
N
0
1
2 km

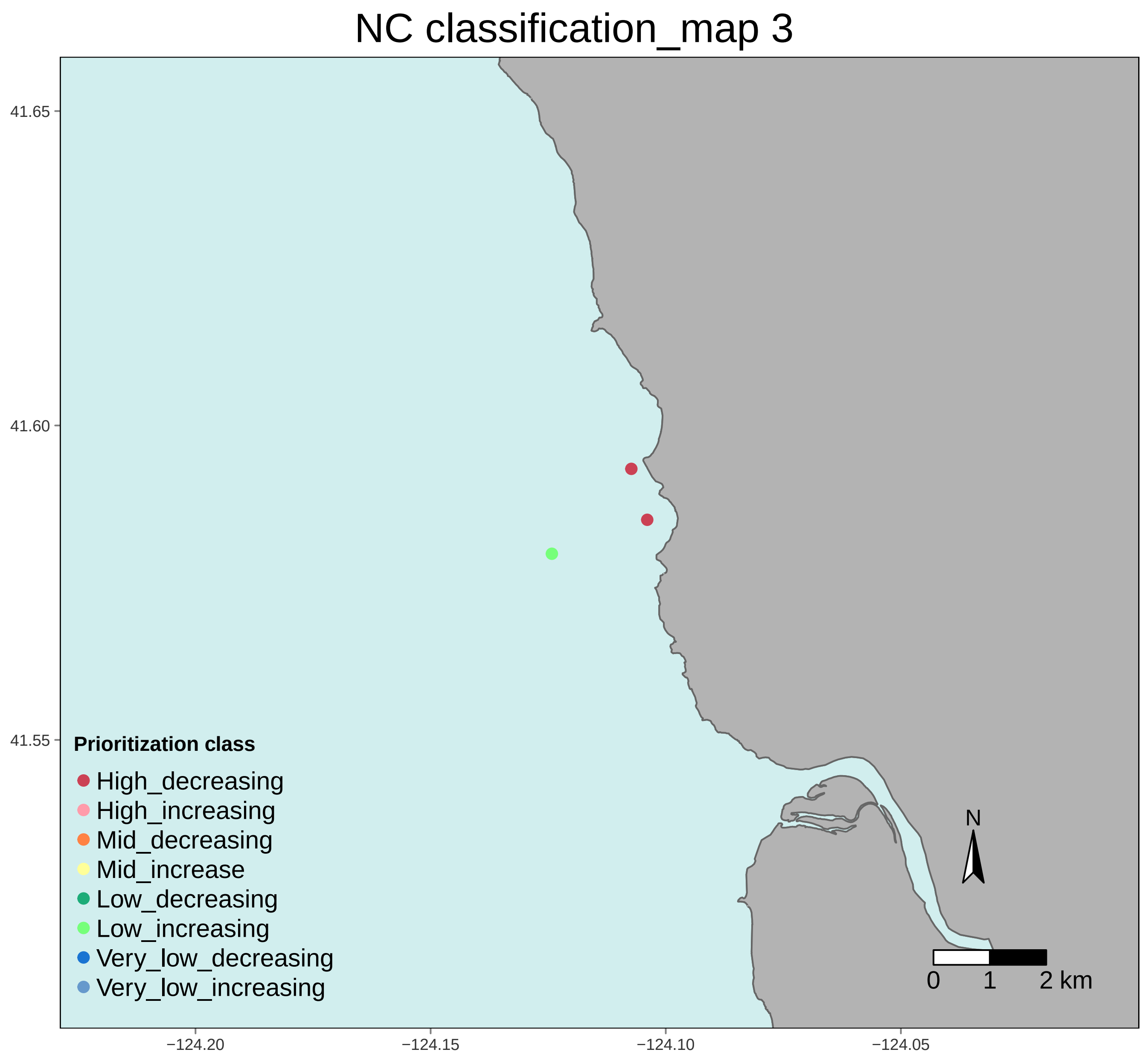
NC classification_map 3
41.65
41.60
41.55
−124.20
−124.15
−124.10
−124.05
Prioritization class
High_decreasing
High_increasing
Mid_decreasing
Mid_increase
Low_decreasing
Low_increasing
Very_low_decreasing
Very_low_increasing
N
0
1
2 km

### NC classification_map 4

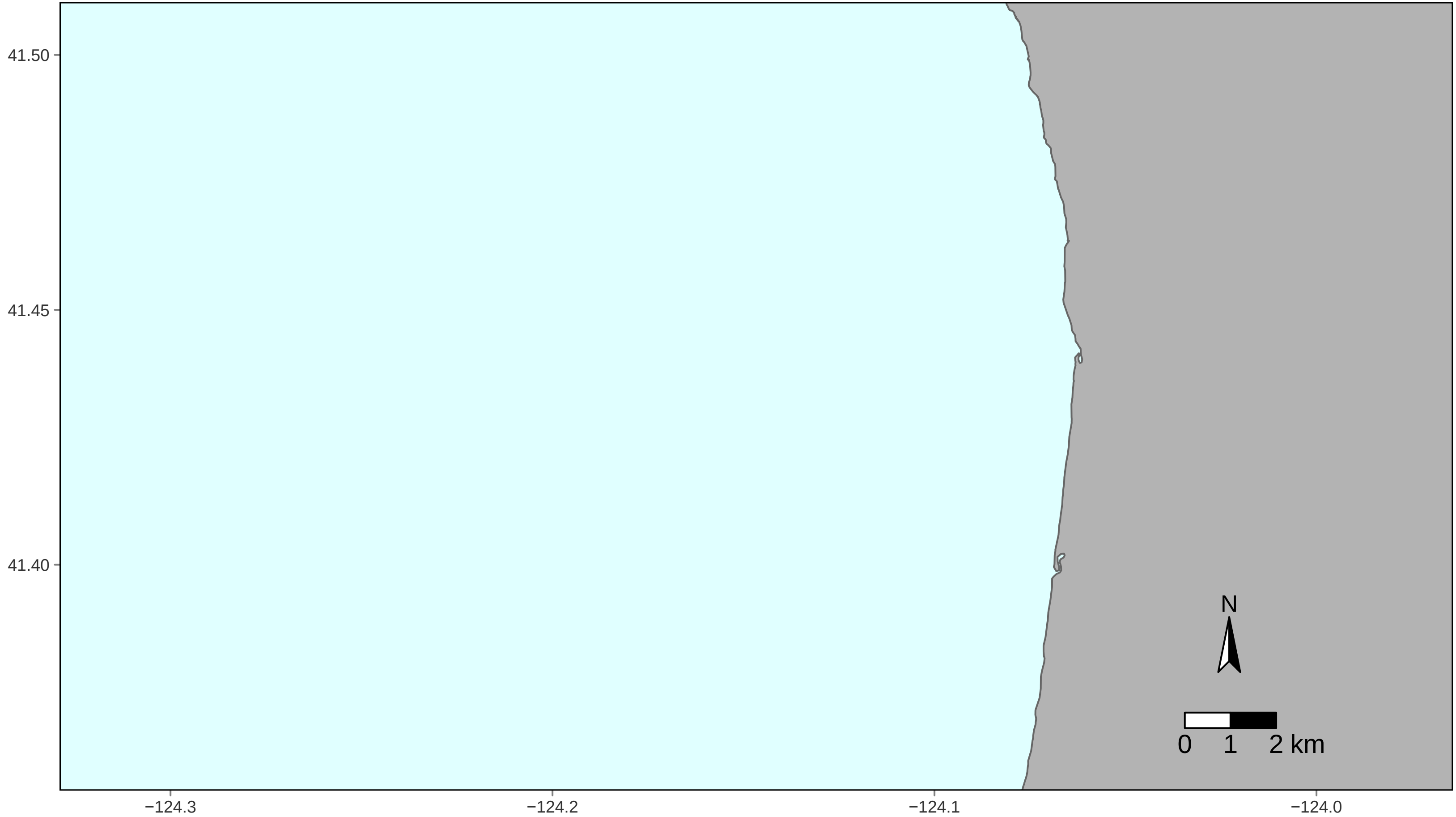

### NC classification_map 5

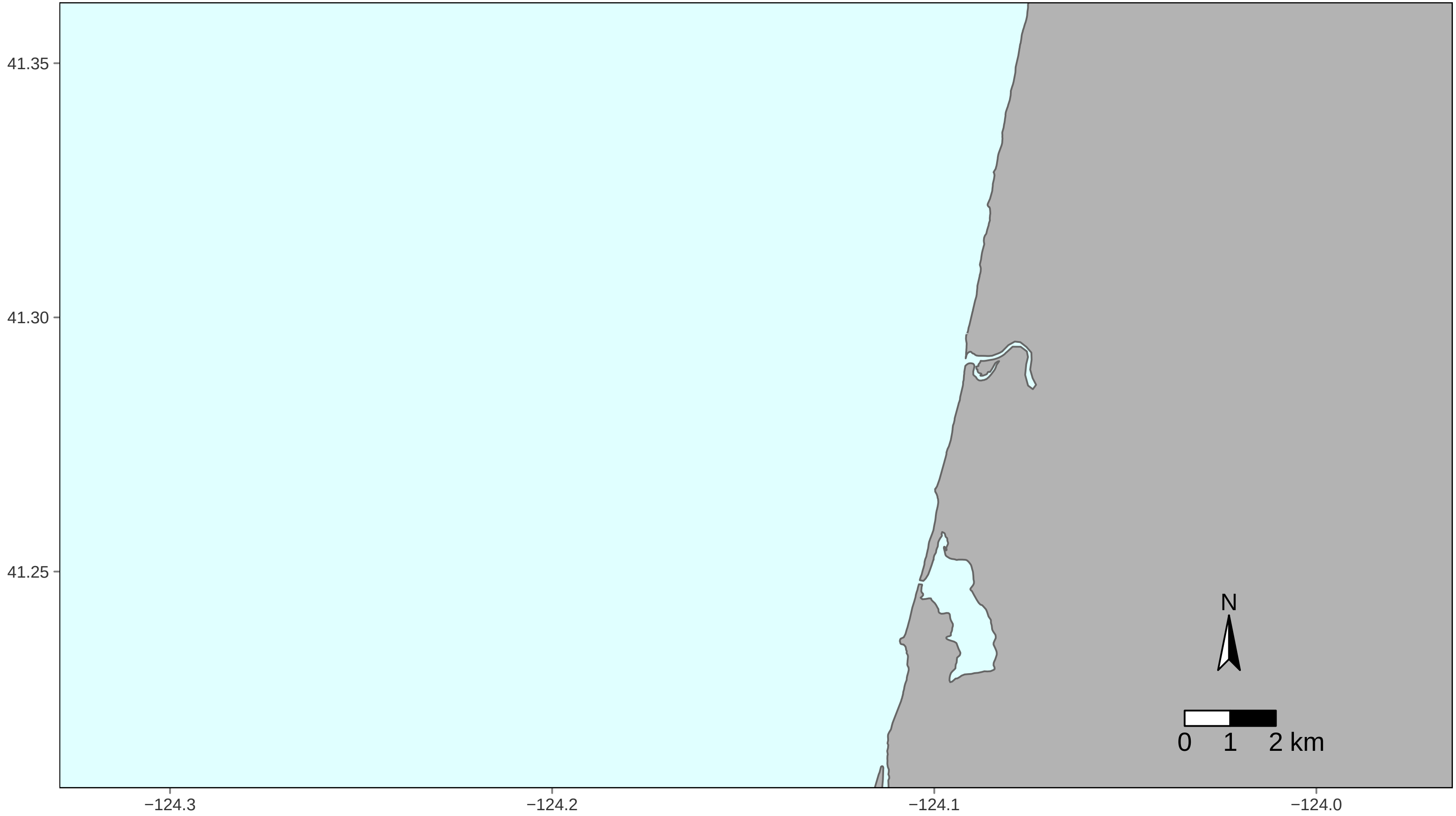

#### NC classification_map 6

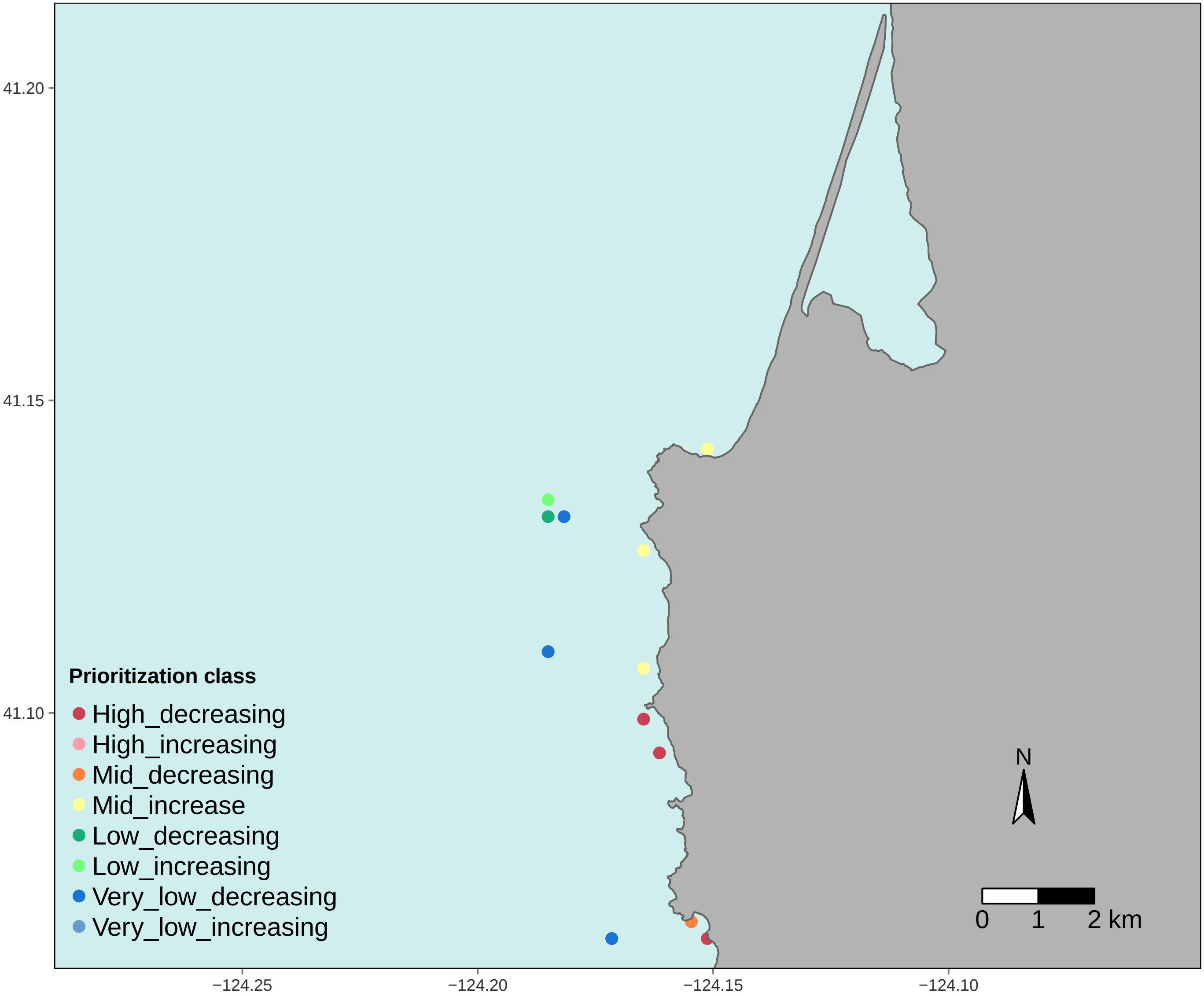

### NC classification_map 7

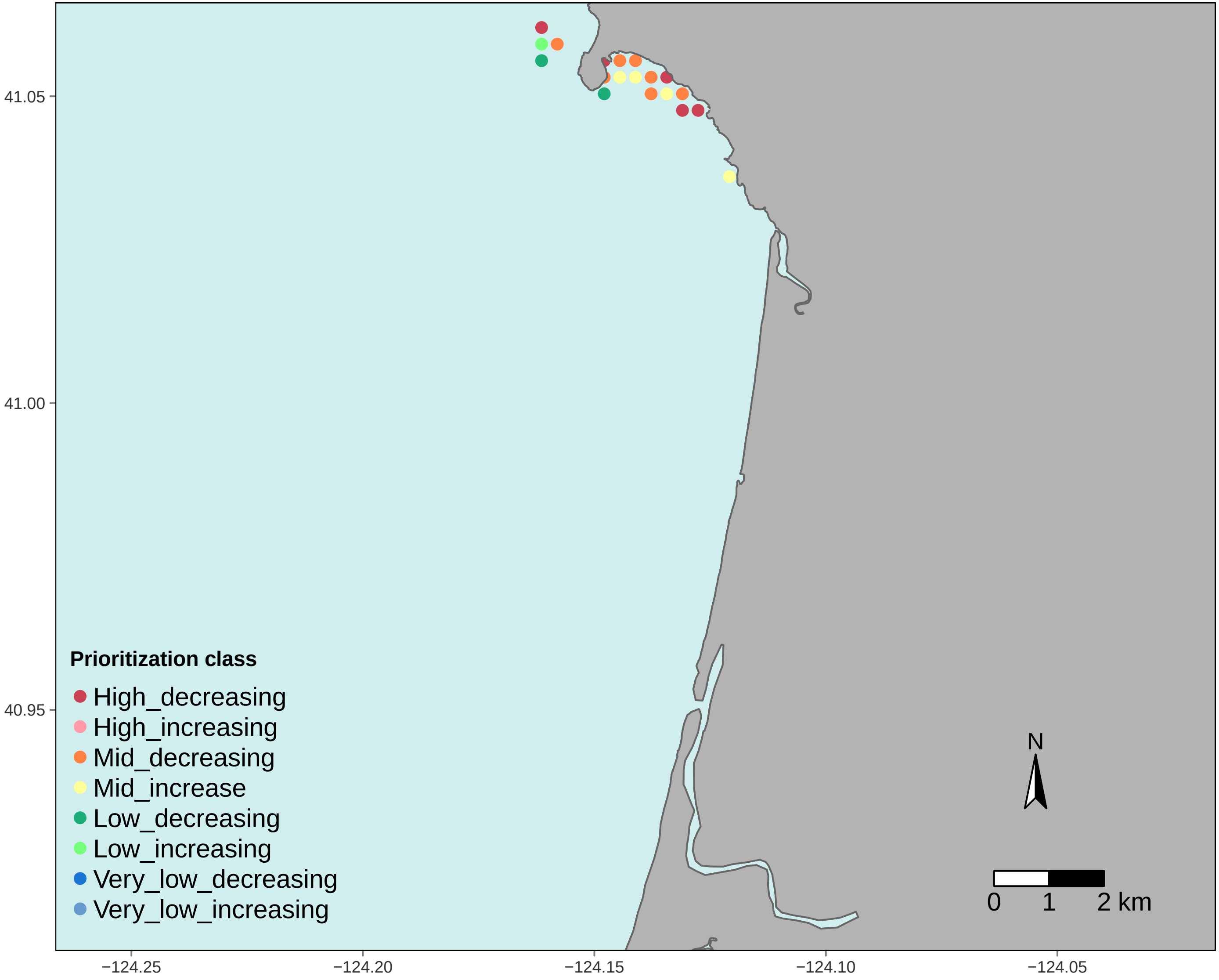

### NC classification_map 8

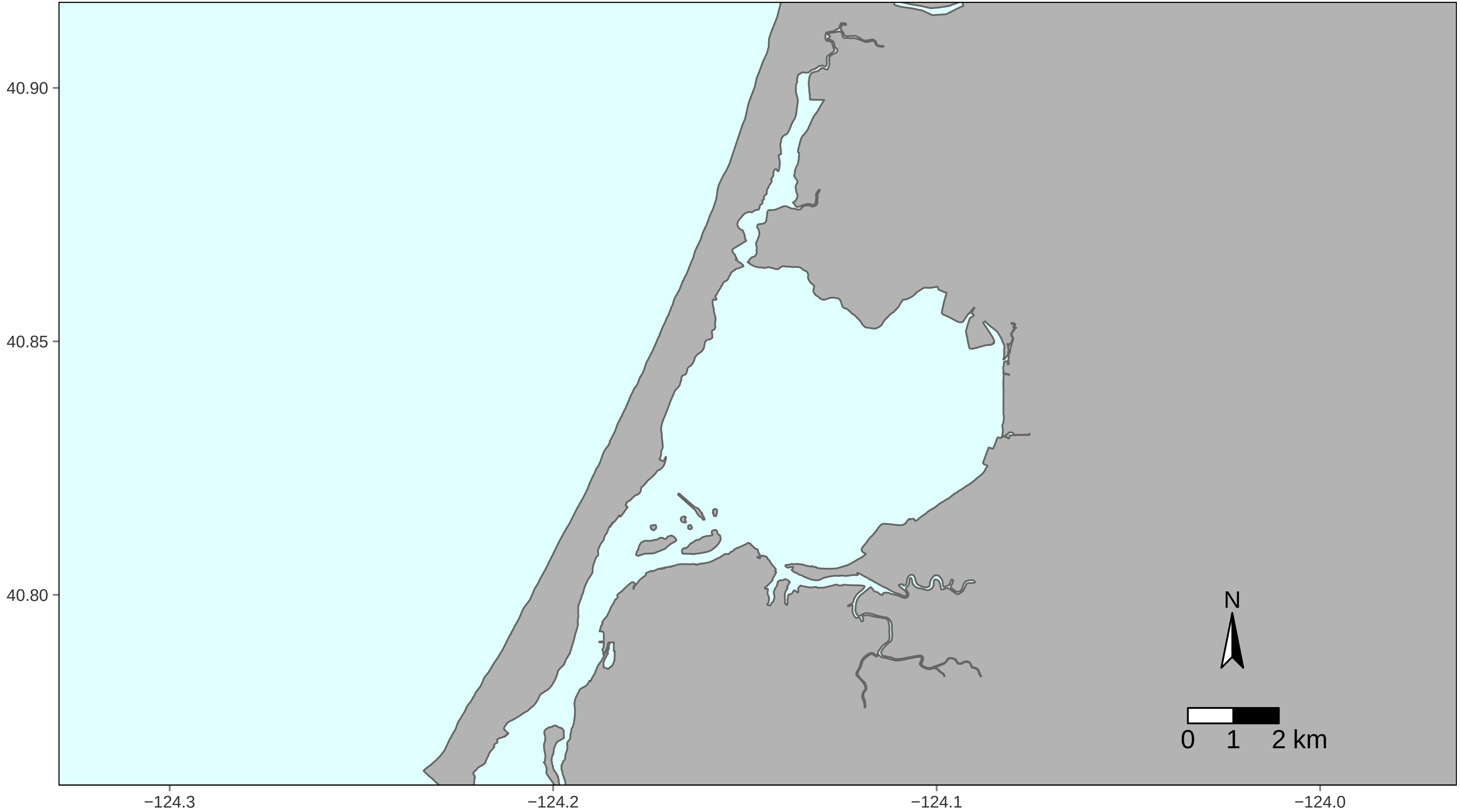

### NC classification_map 9

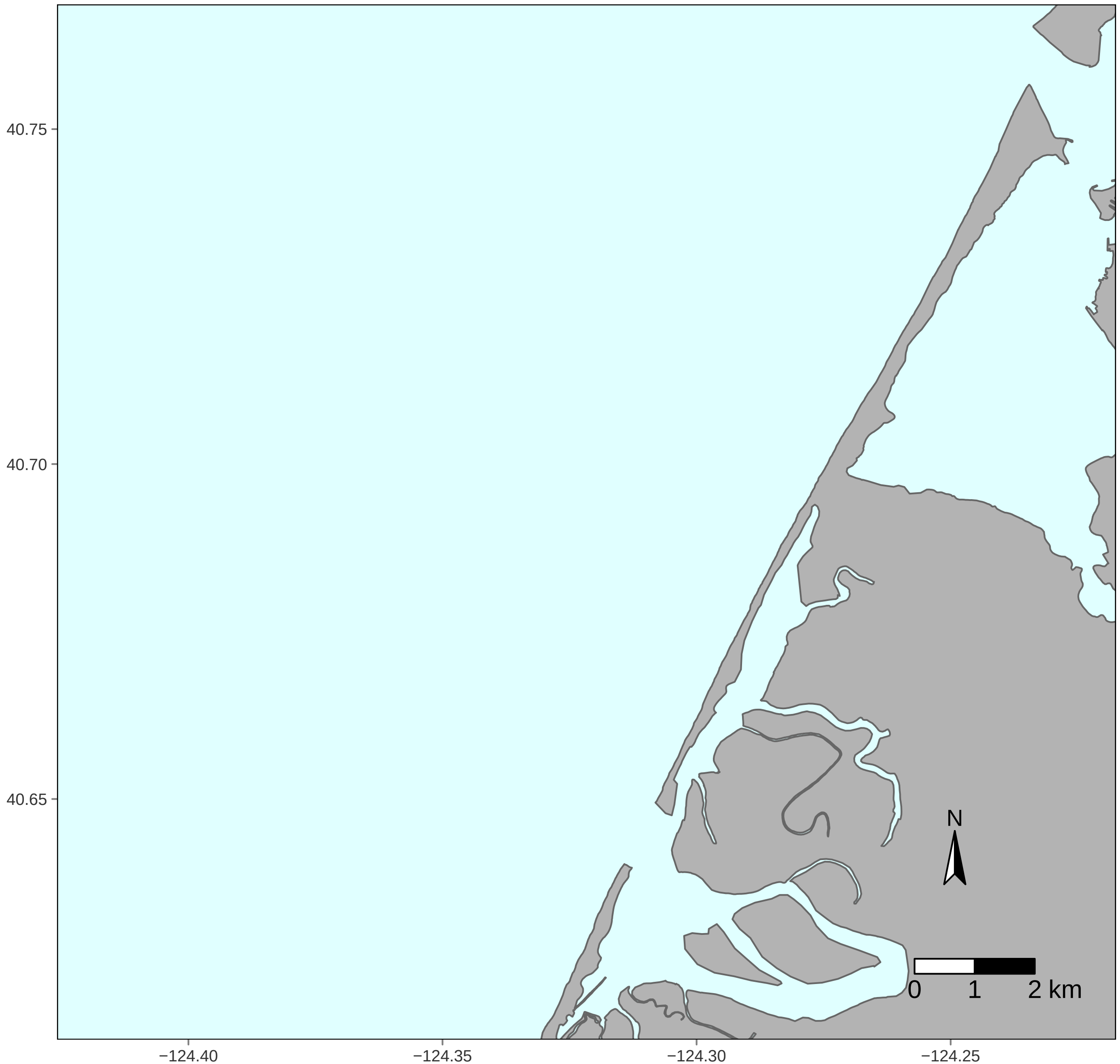

### NC classification_map 10

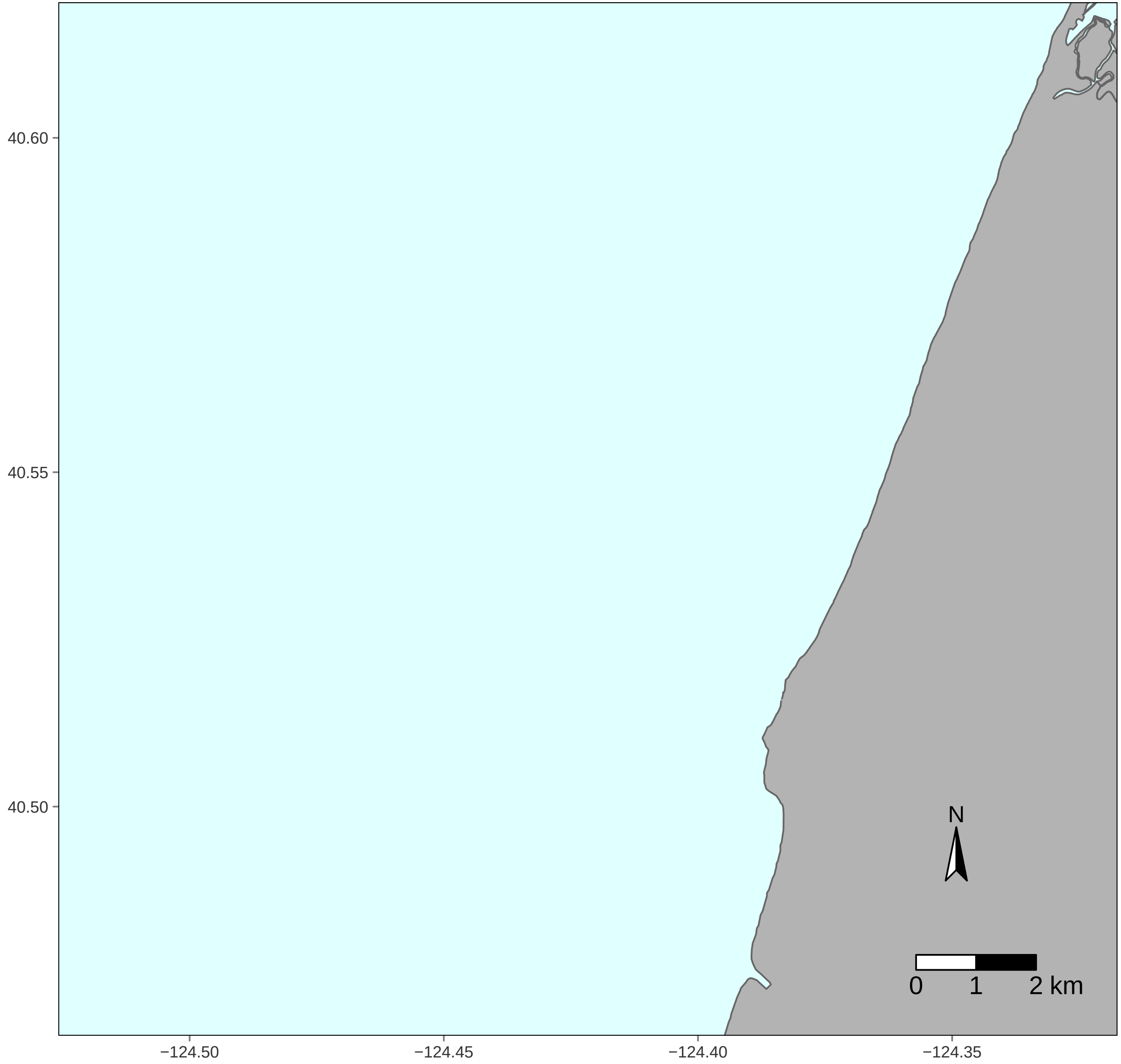

### NC classification_map 11

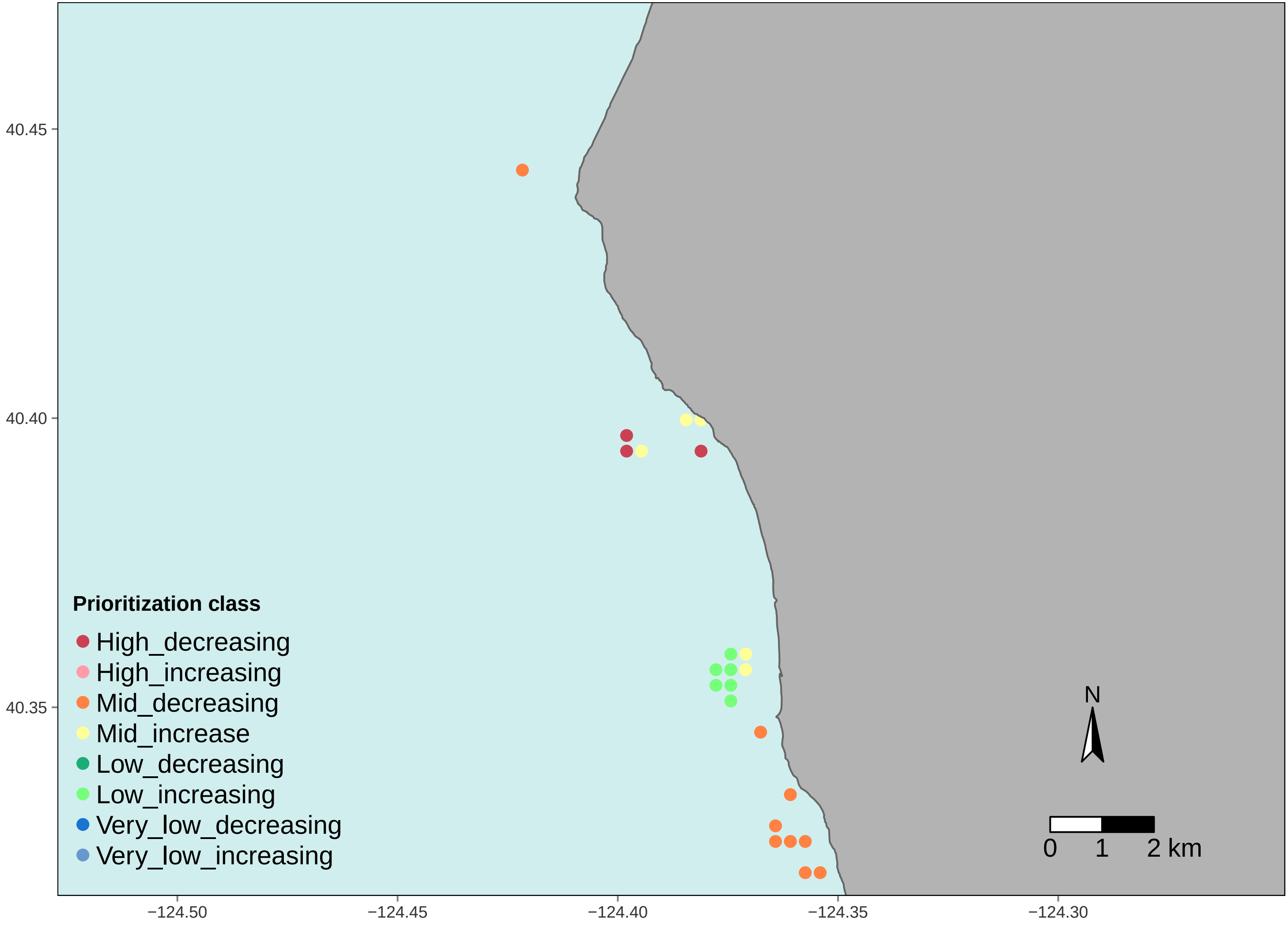

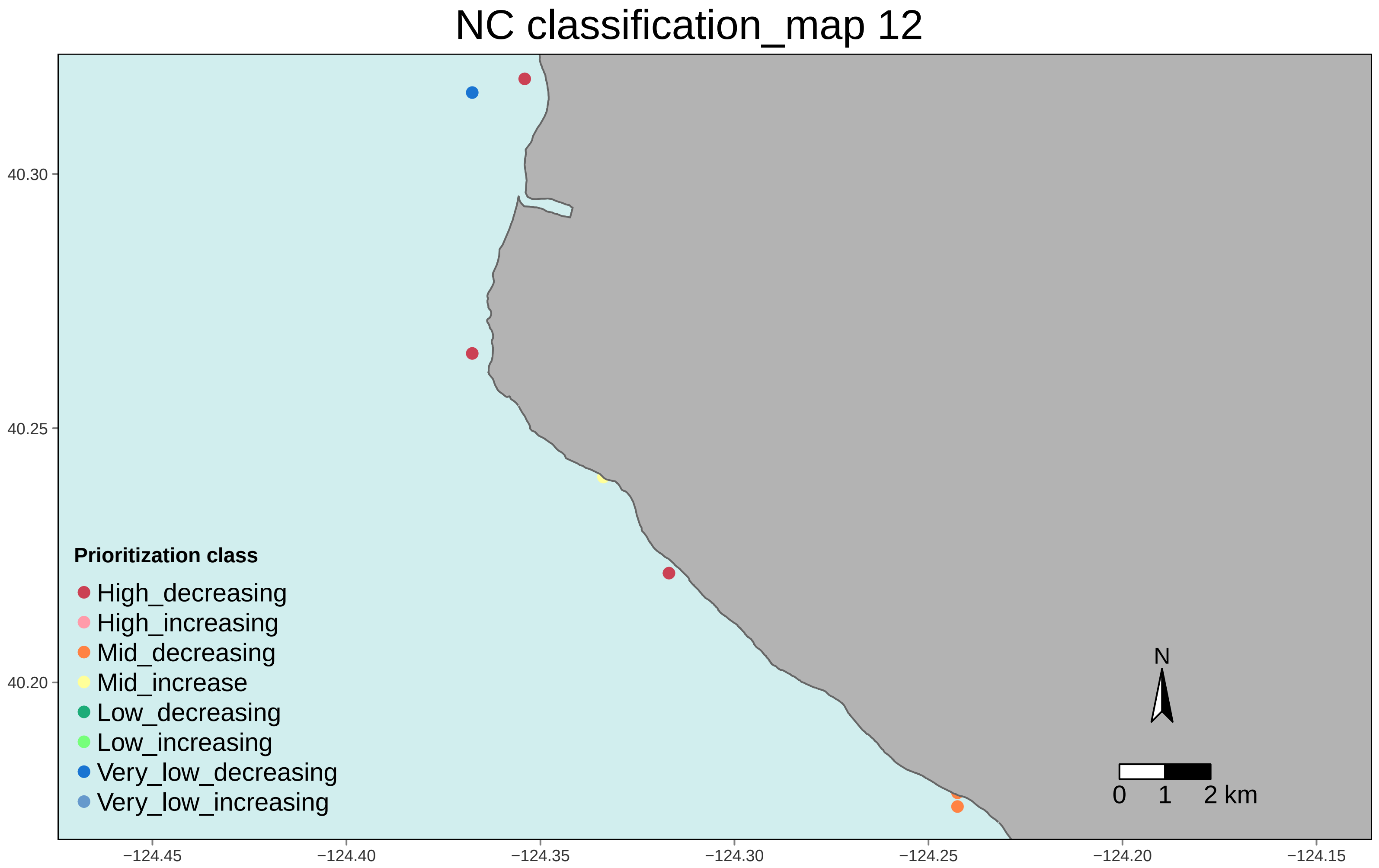

NC classification_map 12
40.30
40.25
40.20
−124.45
−124.40
−124.35
−124.30
−124.25
−124.20
−124.15
Prioritization class
High_decreasing
High_increasing
Mid_decreasing
Mid_increase
Low_decreasing
Low_increasing
Very_low_decreasing
Very_low_increasing
N
0
1
2 km

### NC classification_map 13

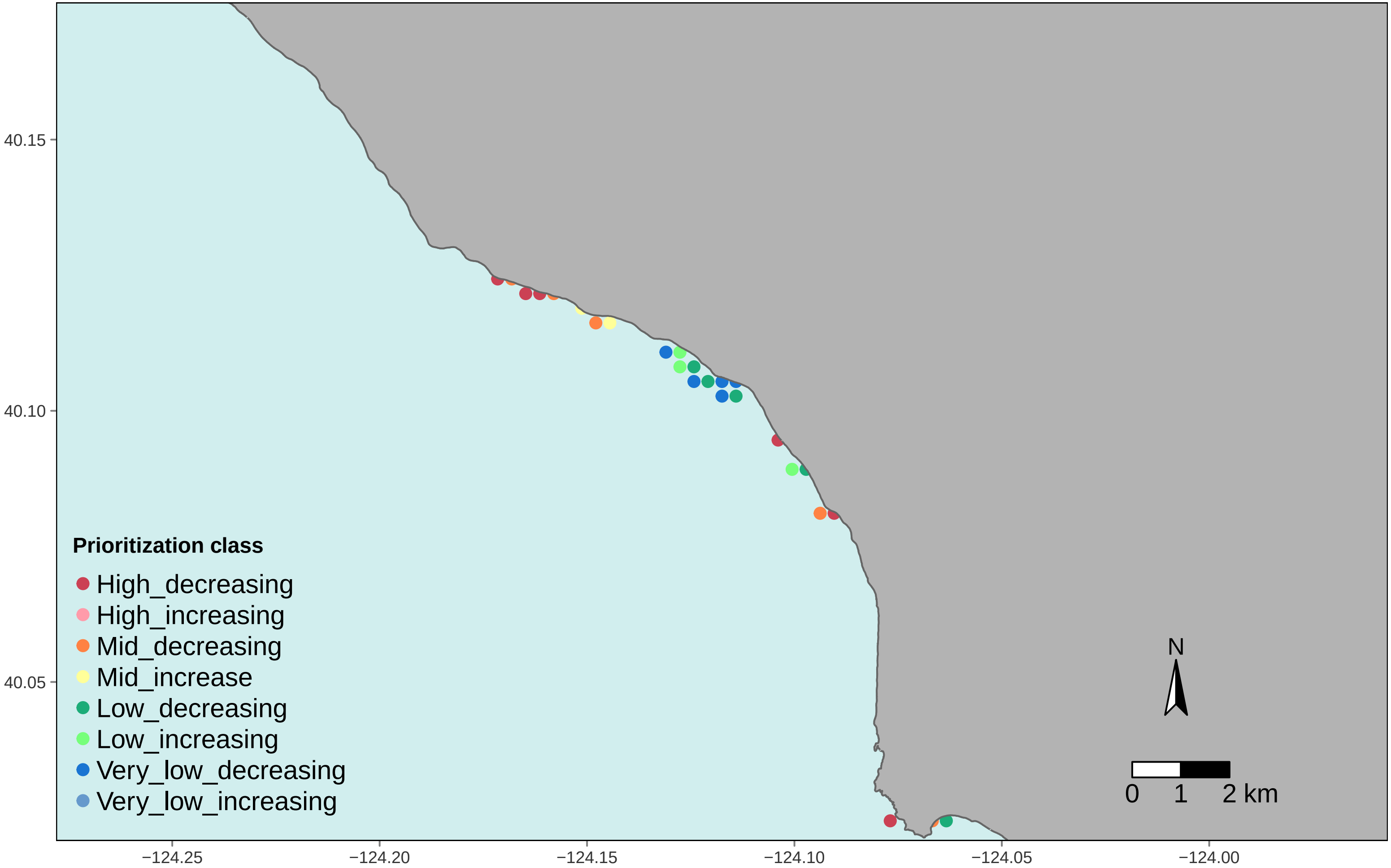

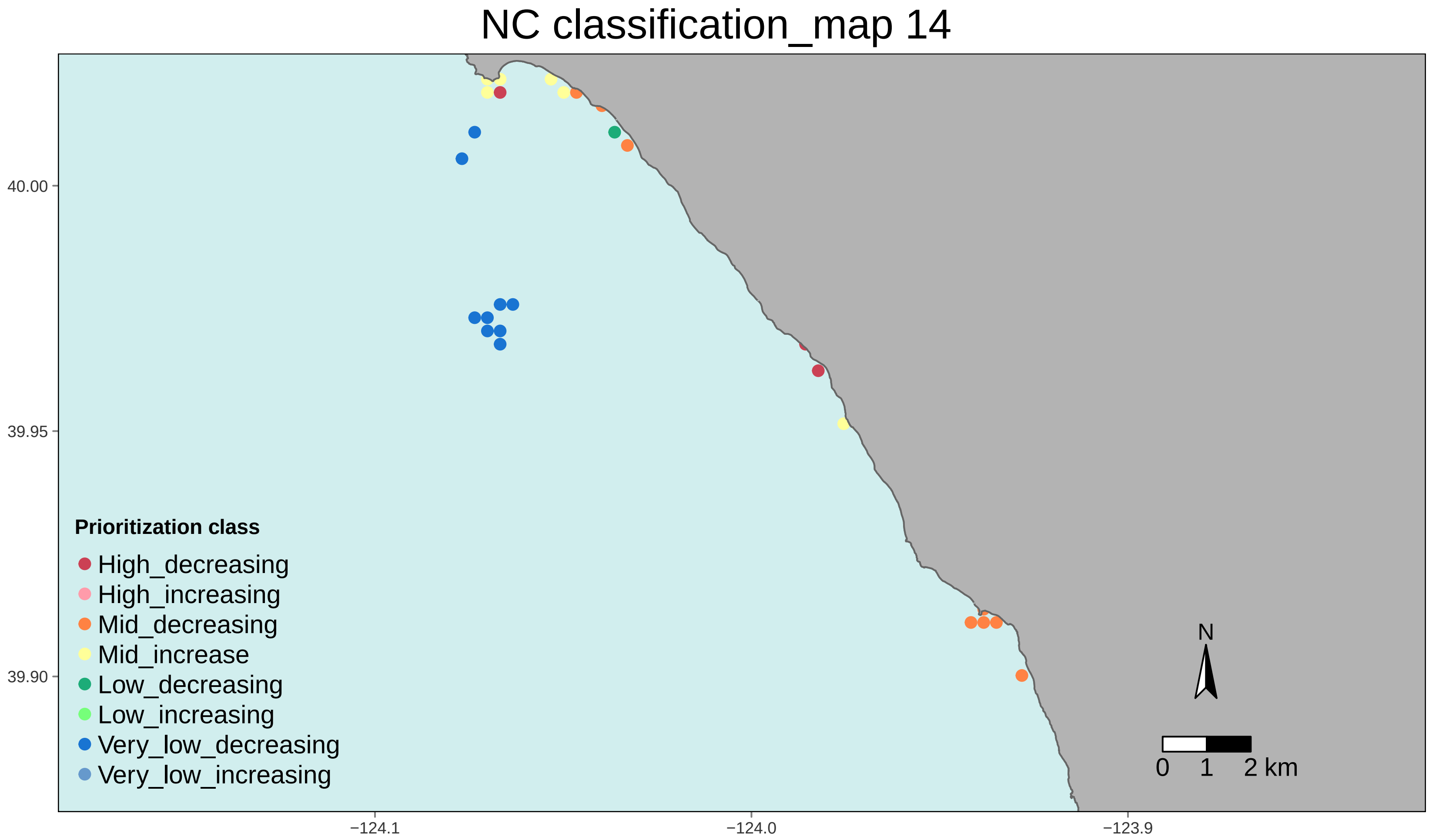

NC classification_map 14
40.00
39.95
39.90
−124.1
−124.0
−123.9
Prioritization class
High_decreasing
High_increasing
Mid_decreasing
Mid_increase
Low_decreasing
Low_increasing
Very_low_decreasing
Very_low_increasing
N
0 1 2 km

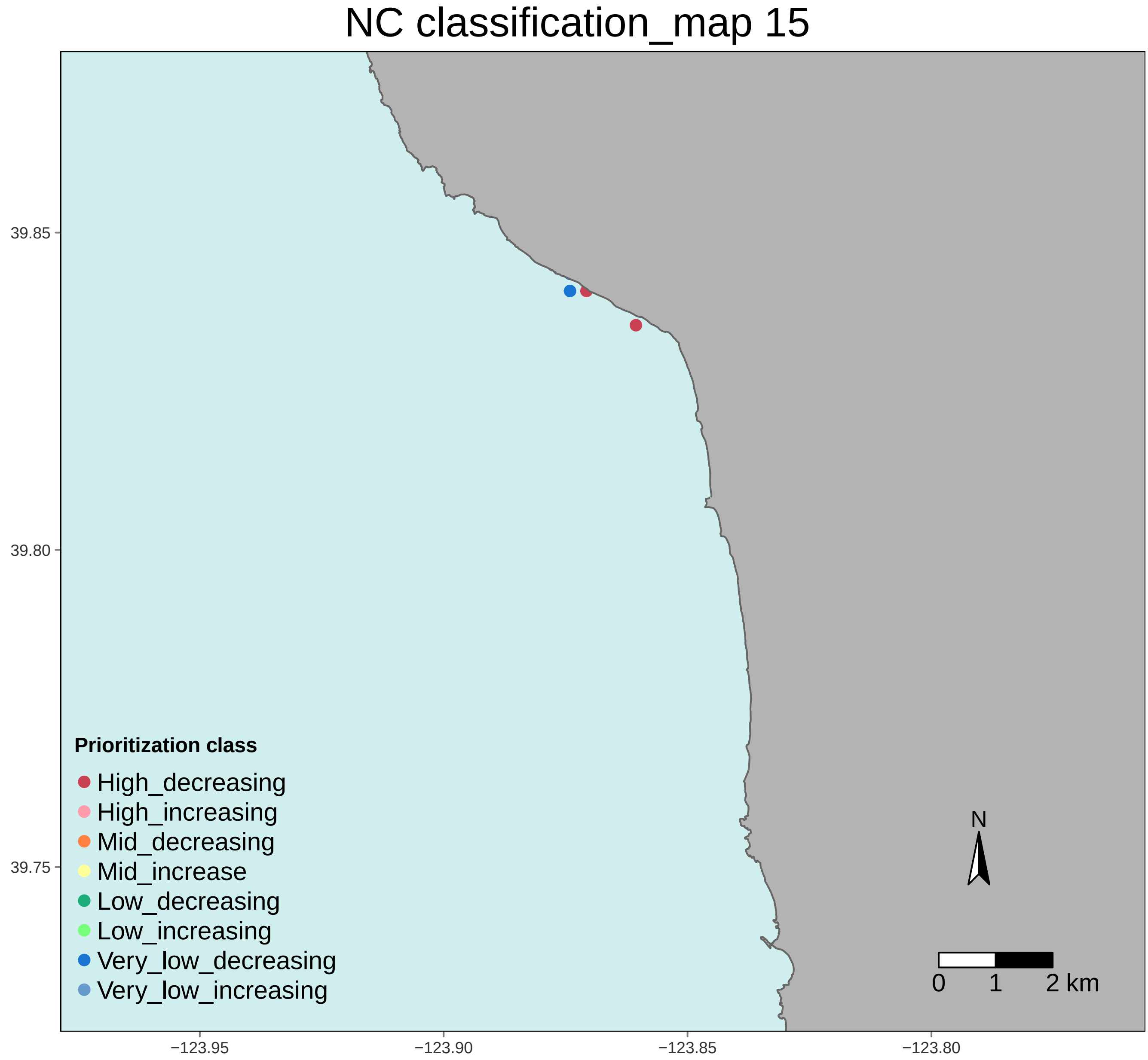
NC classification_map 15
39.85
39.80
39.75
−123.95
−123.90
−123.85
−123.80
Prioritization class
High_decreasing
High_increasing
Mid_decreasing
Mid_increase
Low_decreasing
Low_increasing
Very_low_decreasing
Very_low_increasing
N
0
1
2 km

### NC classification_map 16

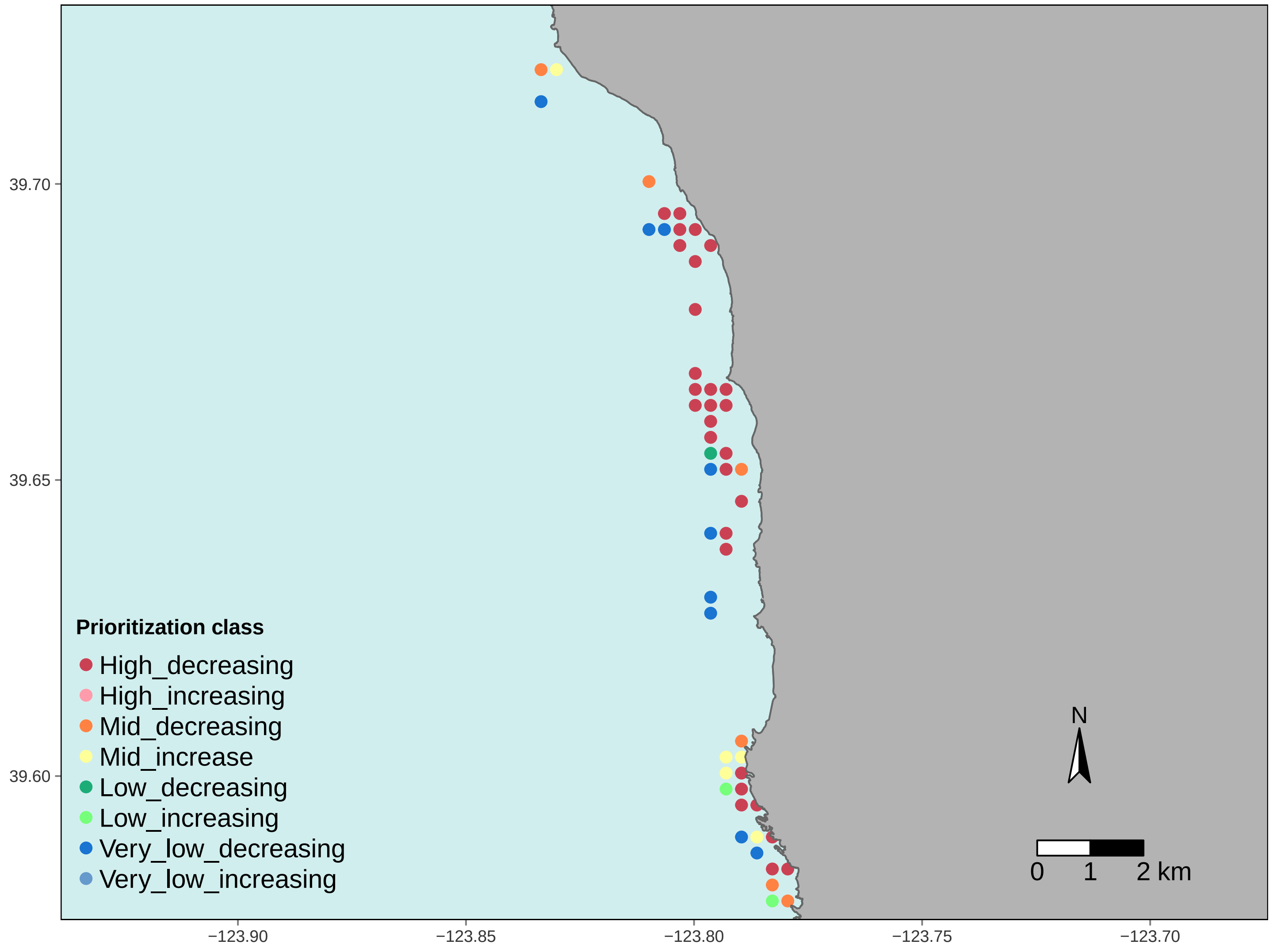

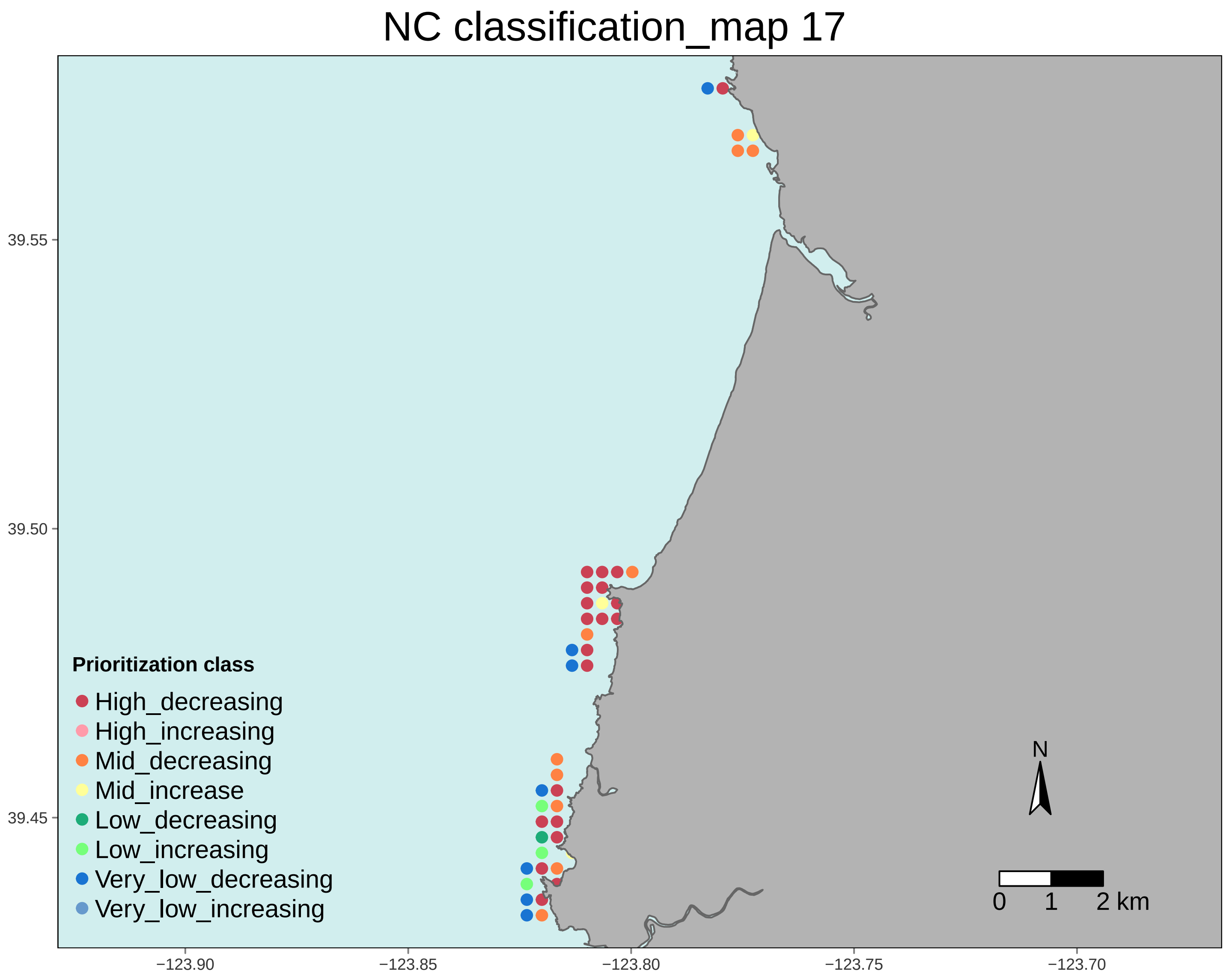
NC classification_map 17
39.55
39.50
39.45
−123.90
−123.85
−123.80
−123.75
−123.70
Prioritization class
High_decreasing
High_increasing
Mid_decreasing
Mid_increase
Low_decreasing
Low_increasing
Very_low_decreasing
Very_low_increasing
N
0
1
2 km

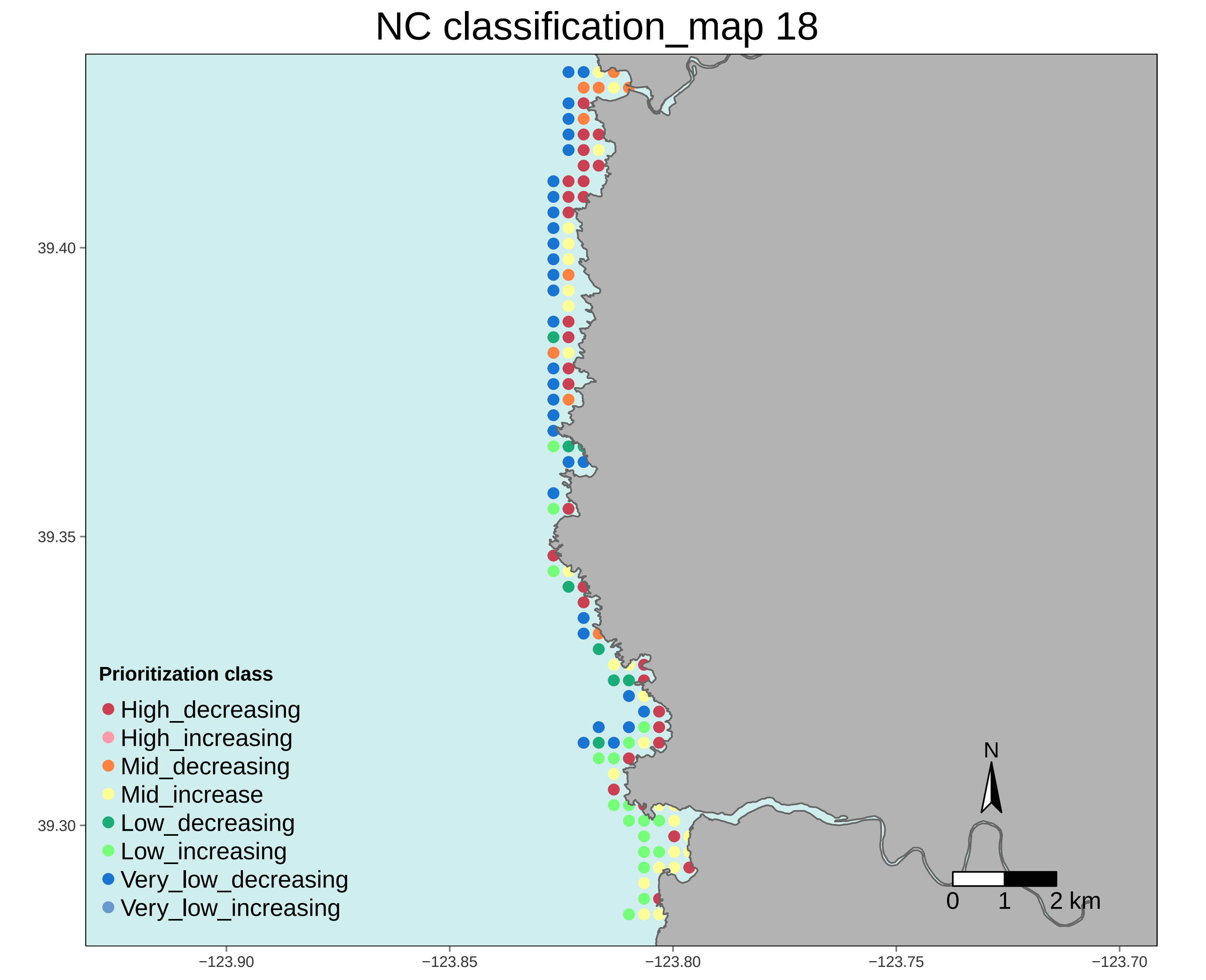
NC classification_map 18
39.40
39.35
39.30
−123.90
−123.85
−123.80
−123.75
−123.70
Prioritization class
High_decreasing
High_increasing
Mid_decreasing
Mid_increase
Low_decreasing
Low_increasing
Very_low_decreasing
Very_low_increasing
N
0
1
2 km

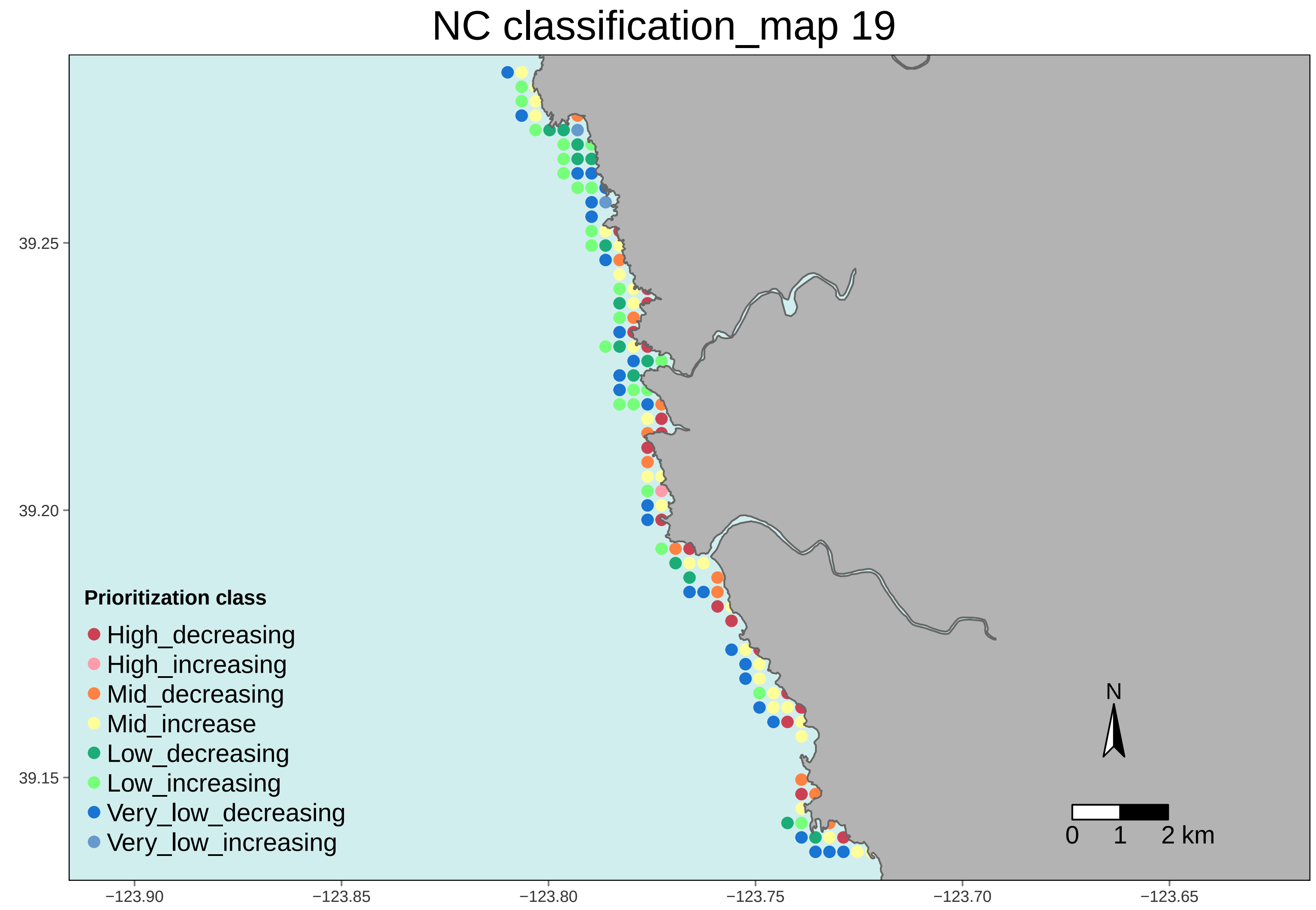
NC classification_map 19
39.25
39.20
39.15
−123.90
−123.85
−123.80
−123.75
−123.70
−123.65
Prioritization class
High_decreasing
High_increasing
Mid_decreasing
Mid_increase
Low_decreasing
Low_increasing
Very_low_decreasing
Very_low_increasing
N
0
1
2 km

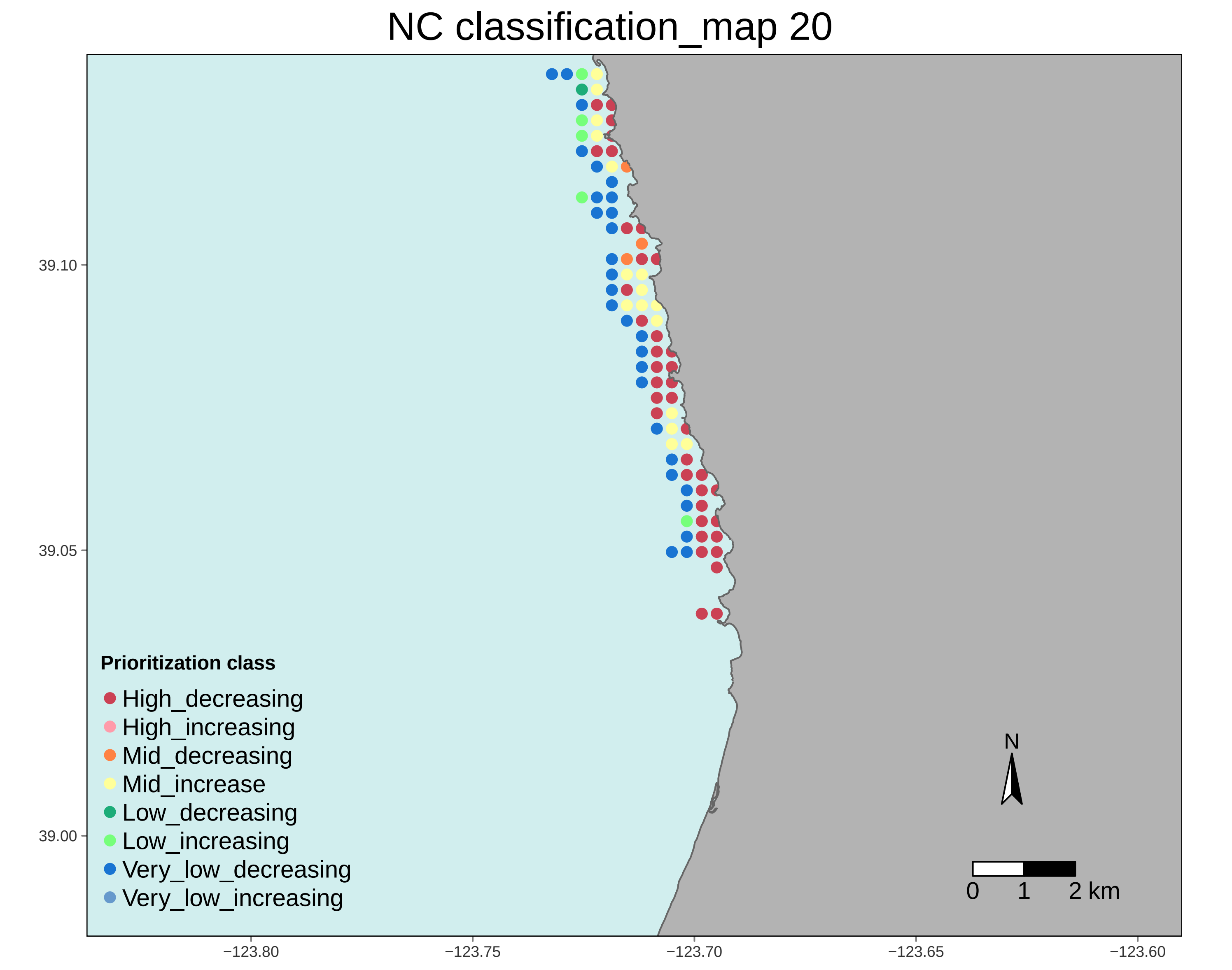
NC classification_map 20
39.10
39.05
39.00
−123.80
−123.75
−123.70
−123.65
−123.60
Prioritization class
High_decreasing
High_increasing
Mid_decreasing
Mid_increase
Low_decreasing
Low_increasing
Very_low_decreasing
Very_low_increasing
N
0
1
2 km

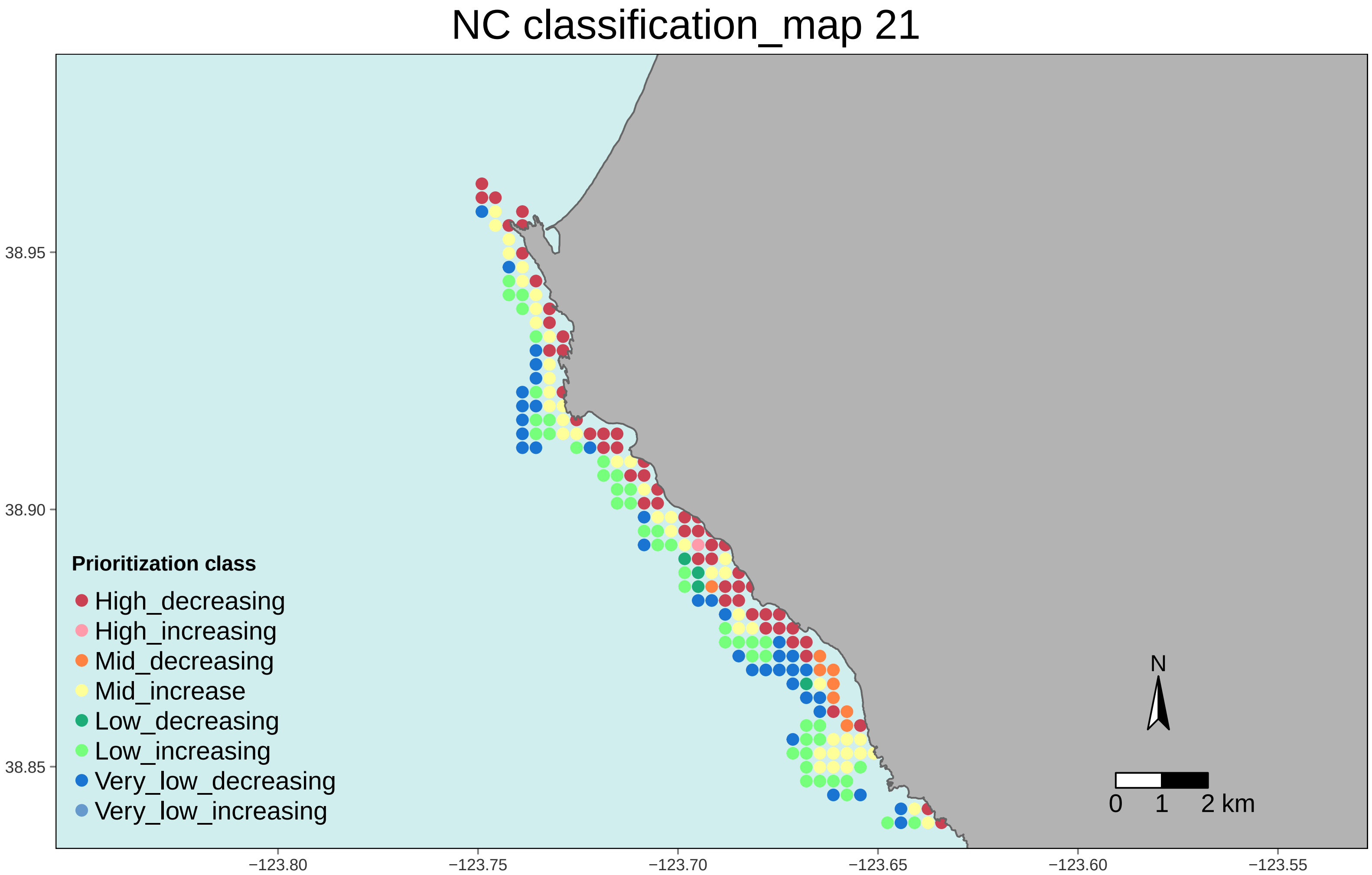
NC classification_map 21
38.95
38.90
38.85
−123.80
−123.75
−123.70
−123.65
−123.60
−123.55
Prioritization class
High_decreasing
High_increasing
Mid_decreasing
Mid_increase
Low_decreasing
Low_increasing
Very_low_decreasing
Very_low_increasing
N
0 1 2 km

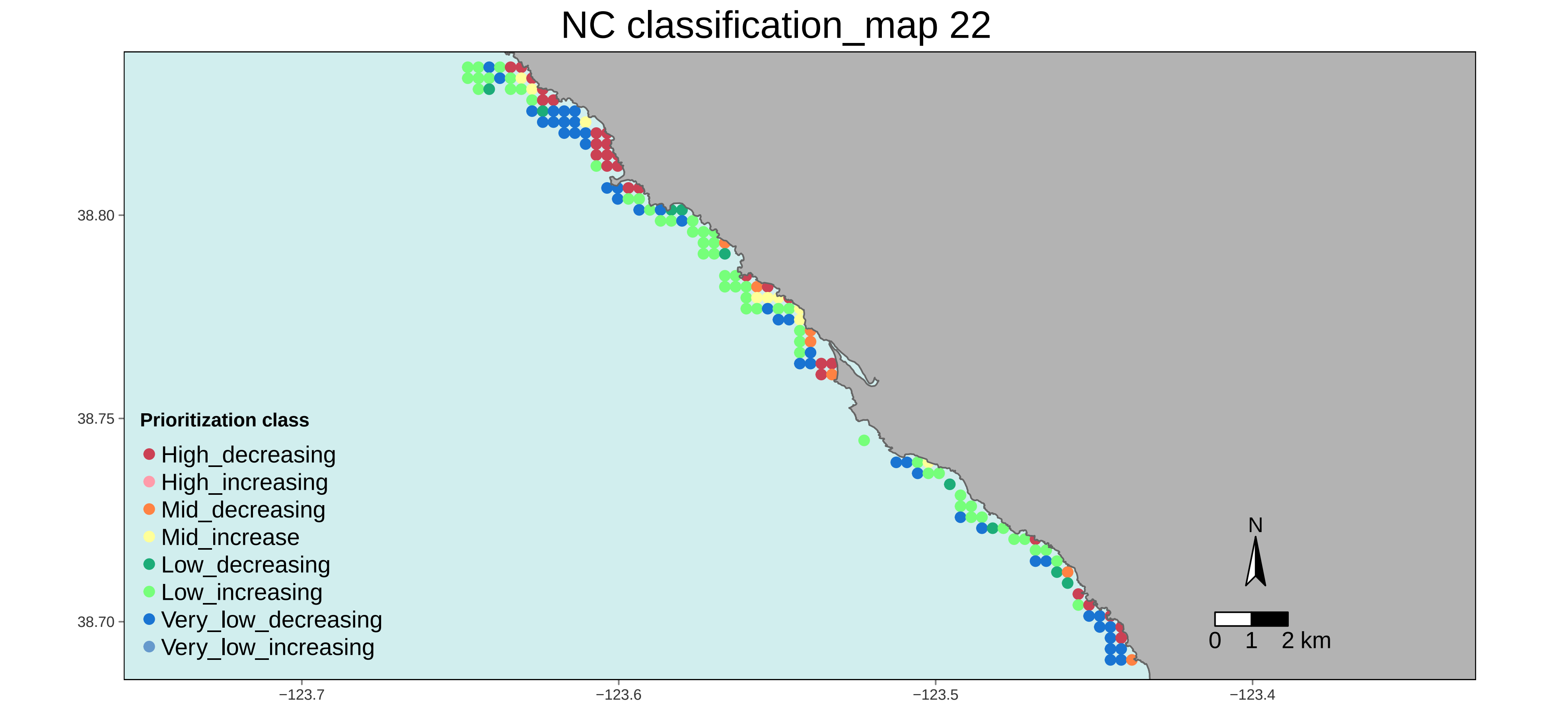
NC classification_map 22
38.80
38.75
38.70
−123.7
−123.6
−123.5
−123.4
Prioritization class
High_decreasing
High_increasing
Mid_decreasing
Mid_increase
Low_decreasing
Low_increasing
Very_low_decreasing
Very_low_increasing
N
0
1
2 km

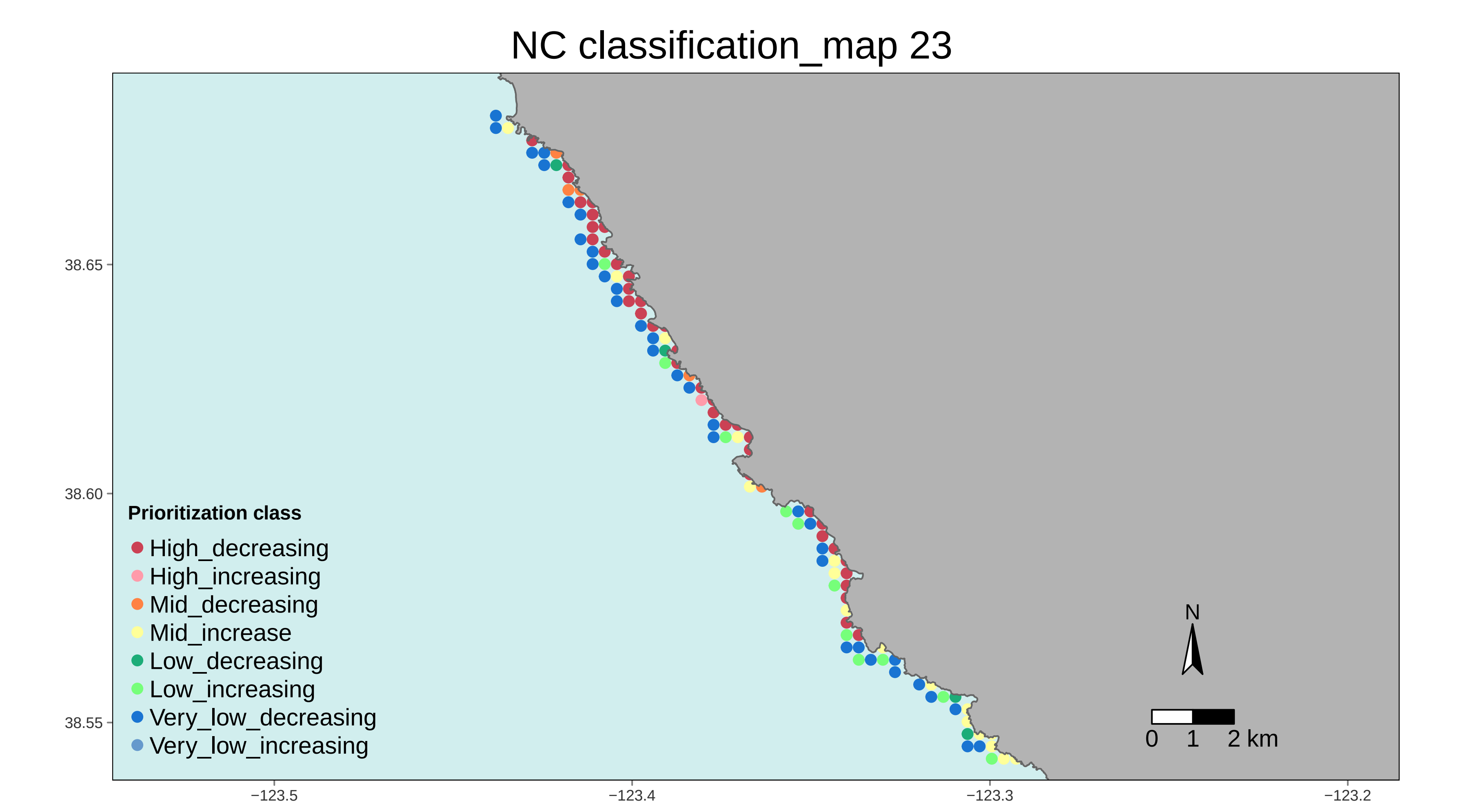
NC classification_map 23
38.65
38.60
38.55
−123.5
−123.4
−123.3
−123.2
Prioritization class
High_decreasing
High_increasing
Mid_decreasing
Mid_increase
Low_decreasing
Low_increasing
Very_low_decreasing
Very_low_increasing
N
0
1
2 km

### NC classification_map 24

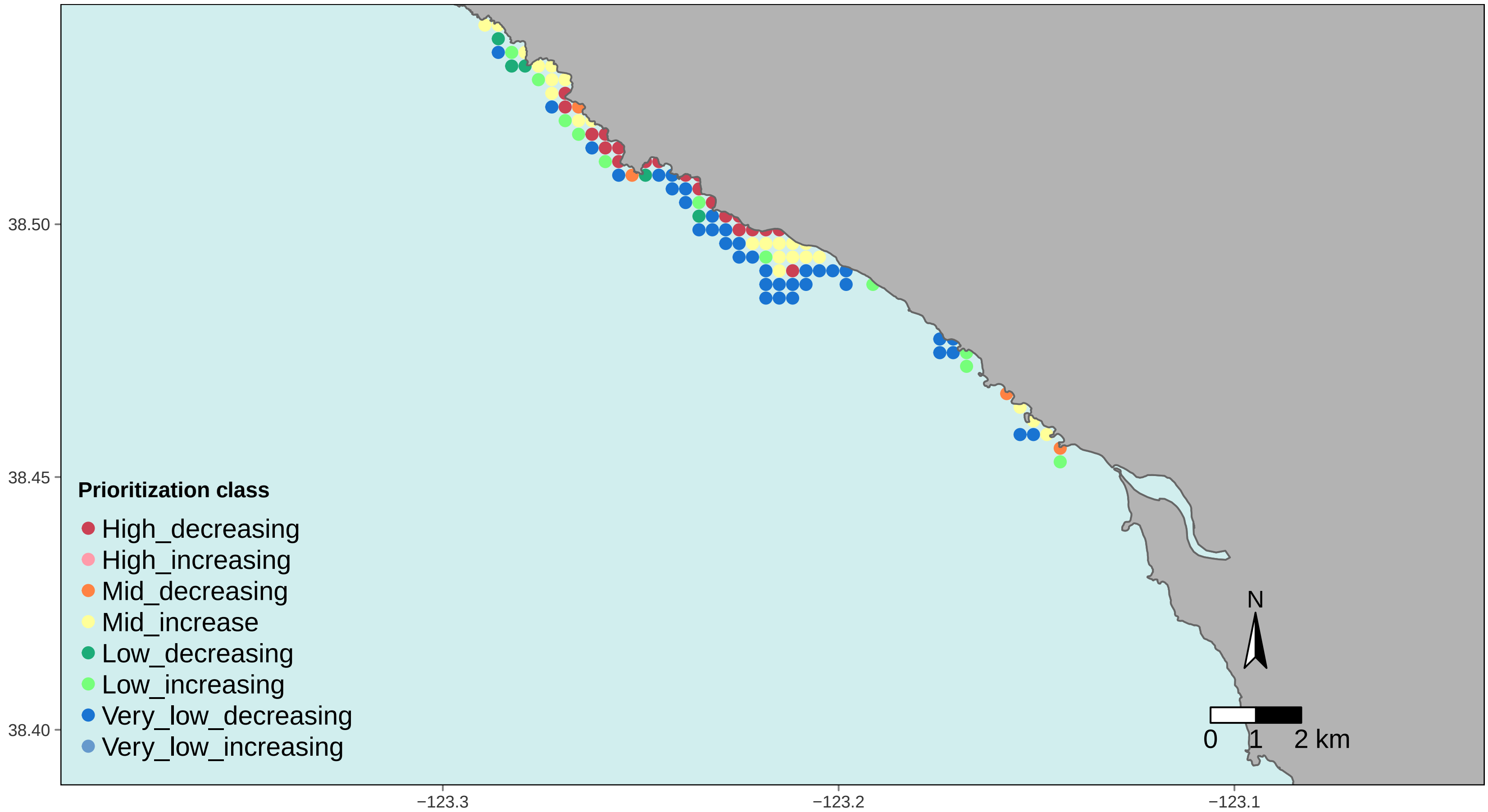

### NC classification_map 25

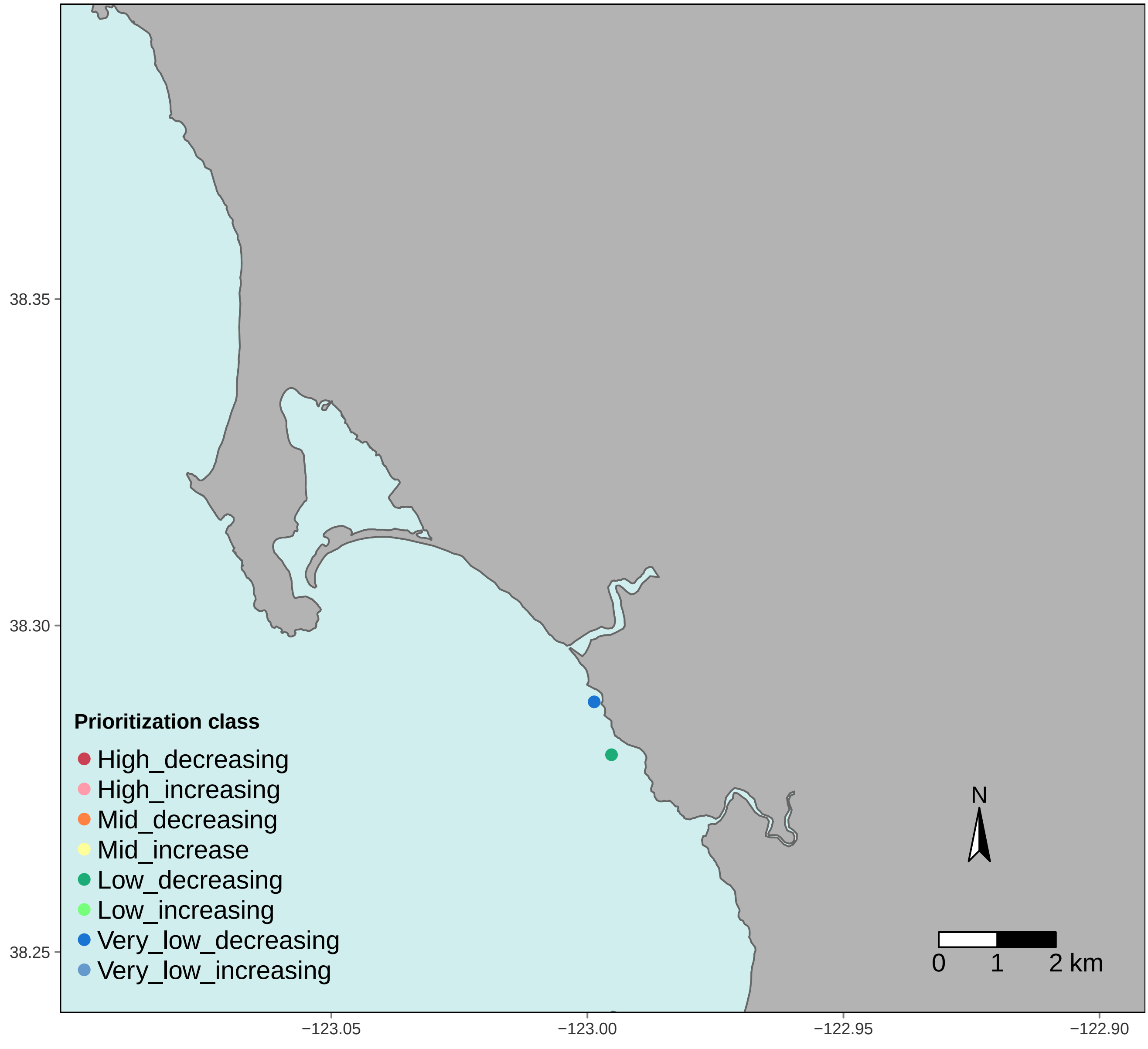

### NC classification_map 26

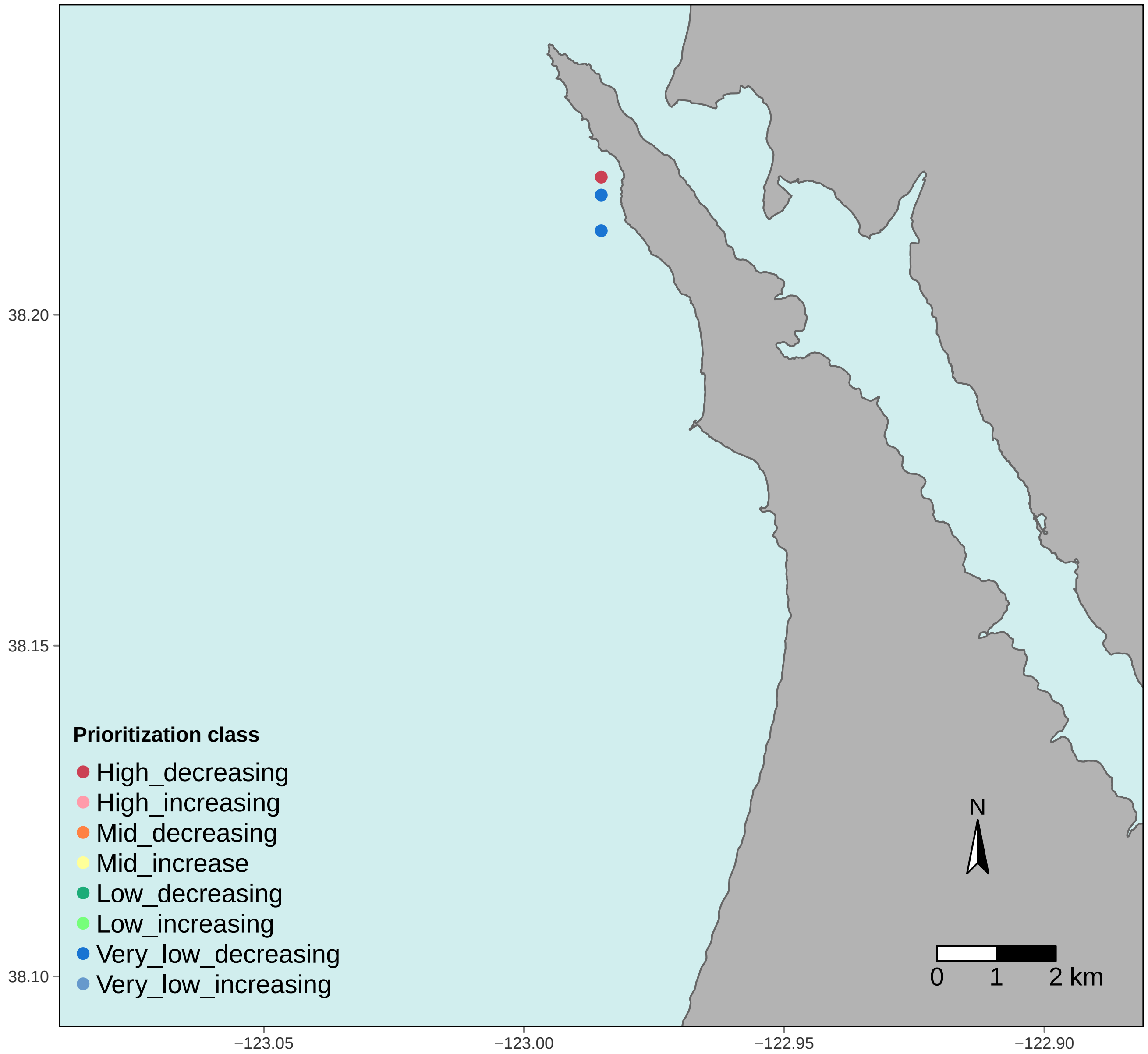

#### NC classification_map 27

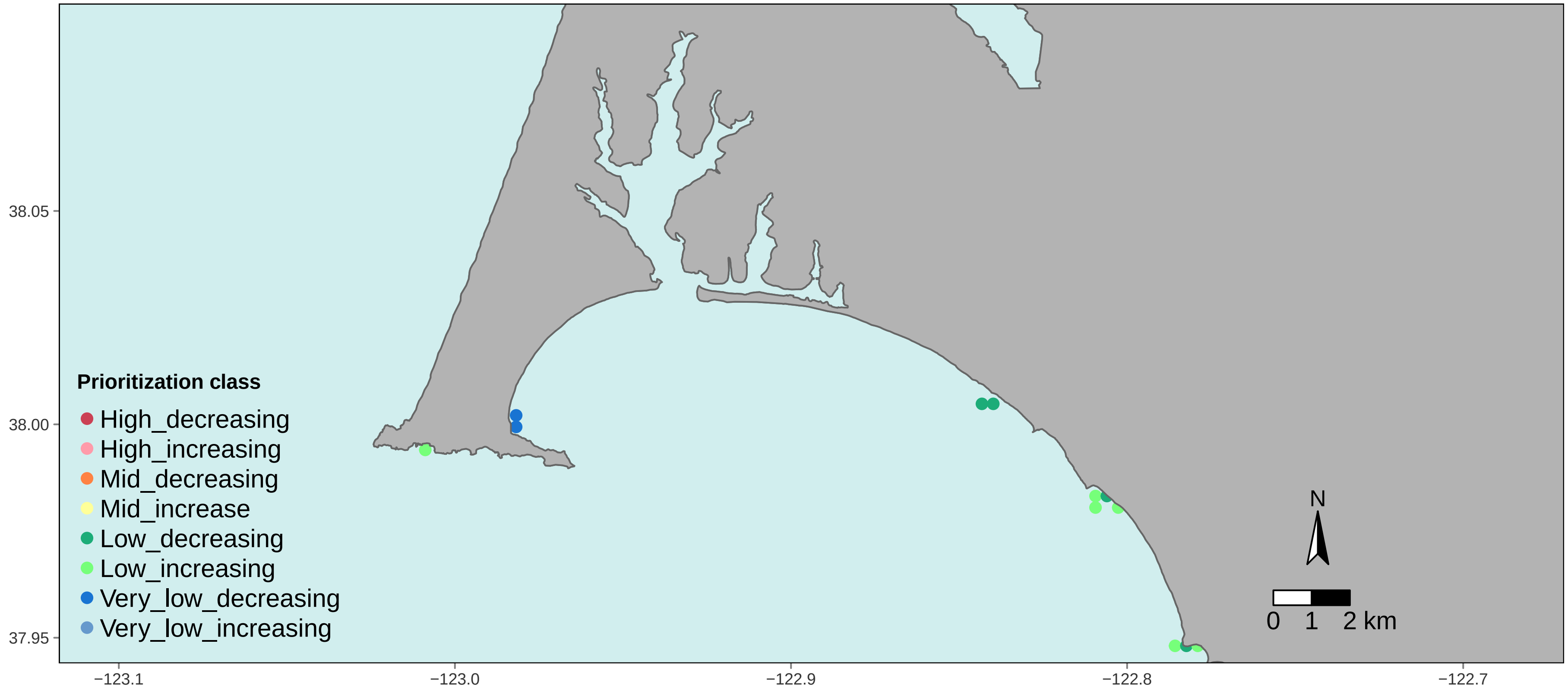

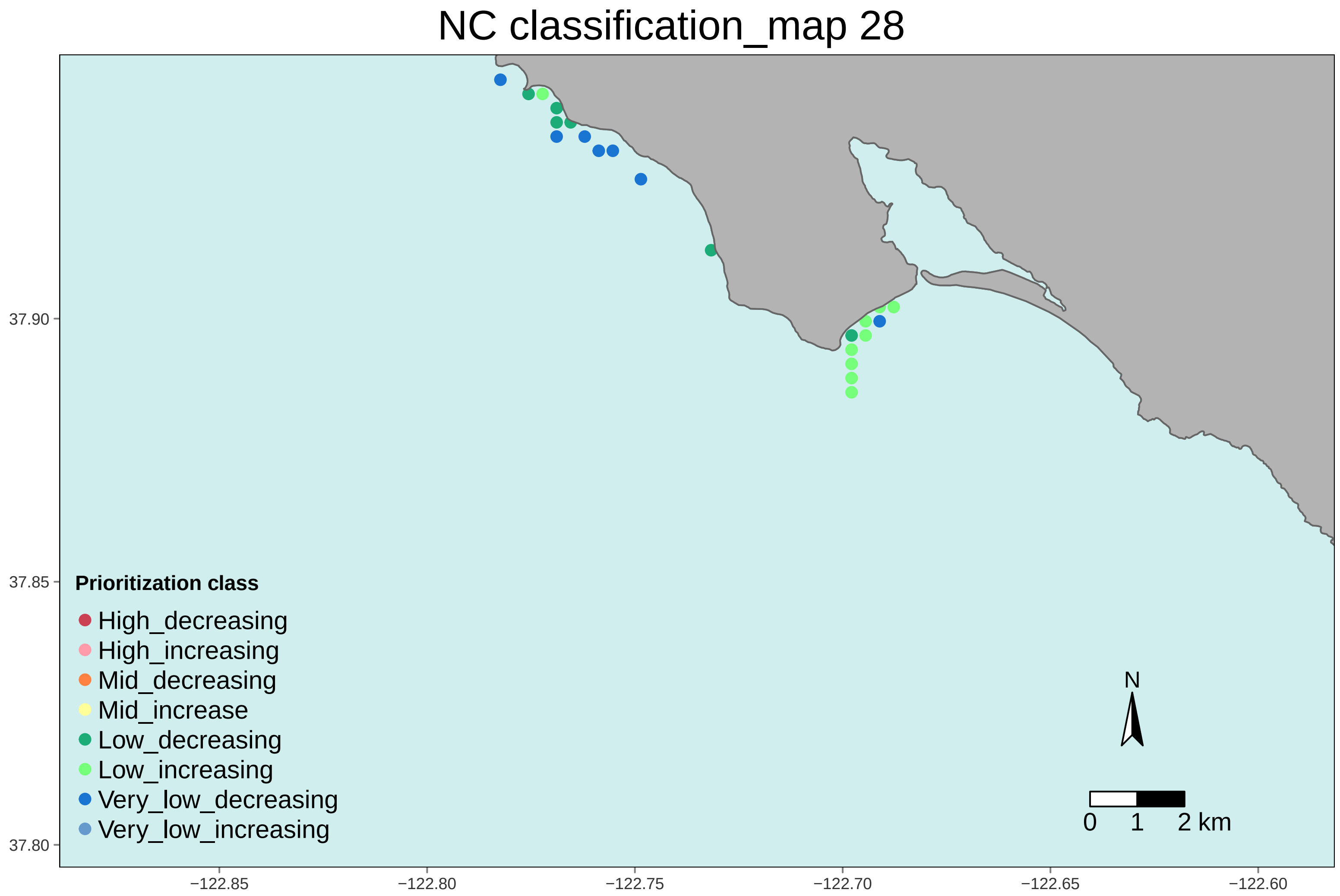
NC classification_map 28
37.90
37.85
37.80
−122.85
−122.80
−122.75
−122.70
−122.65
−122.60
Prioritization class
High_decreasing
High_increasing
Mid_decreasing
Mid_increase
Low_decreasing
Low_increasing
Very_low_decreasing
Very_low_increasing
N
0
1
2 km

NC classification_map 29

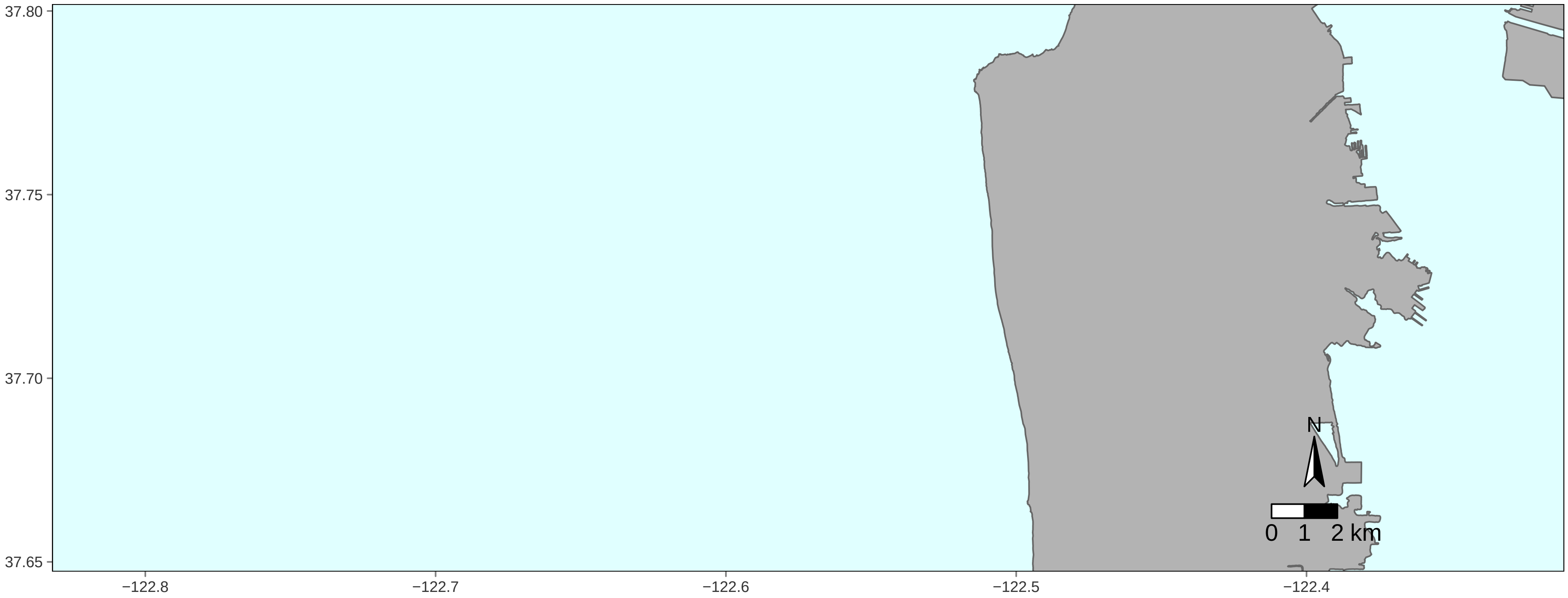

### NC classification_map 30

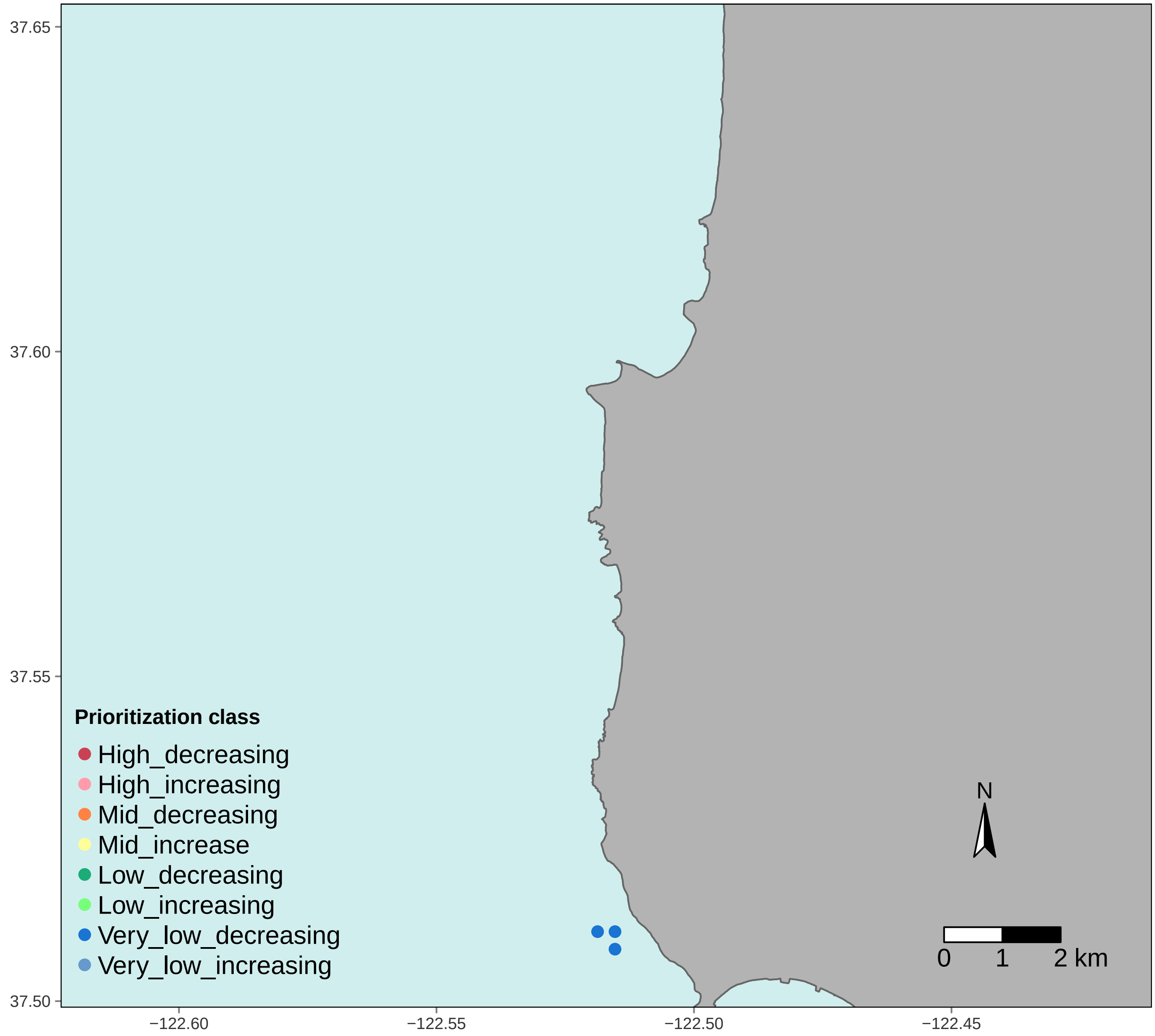

### NC classification_map 31

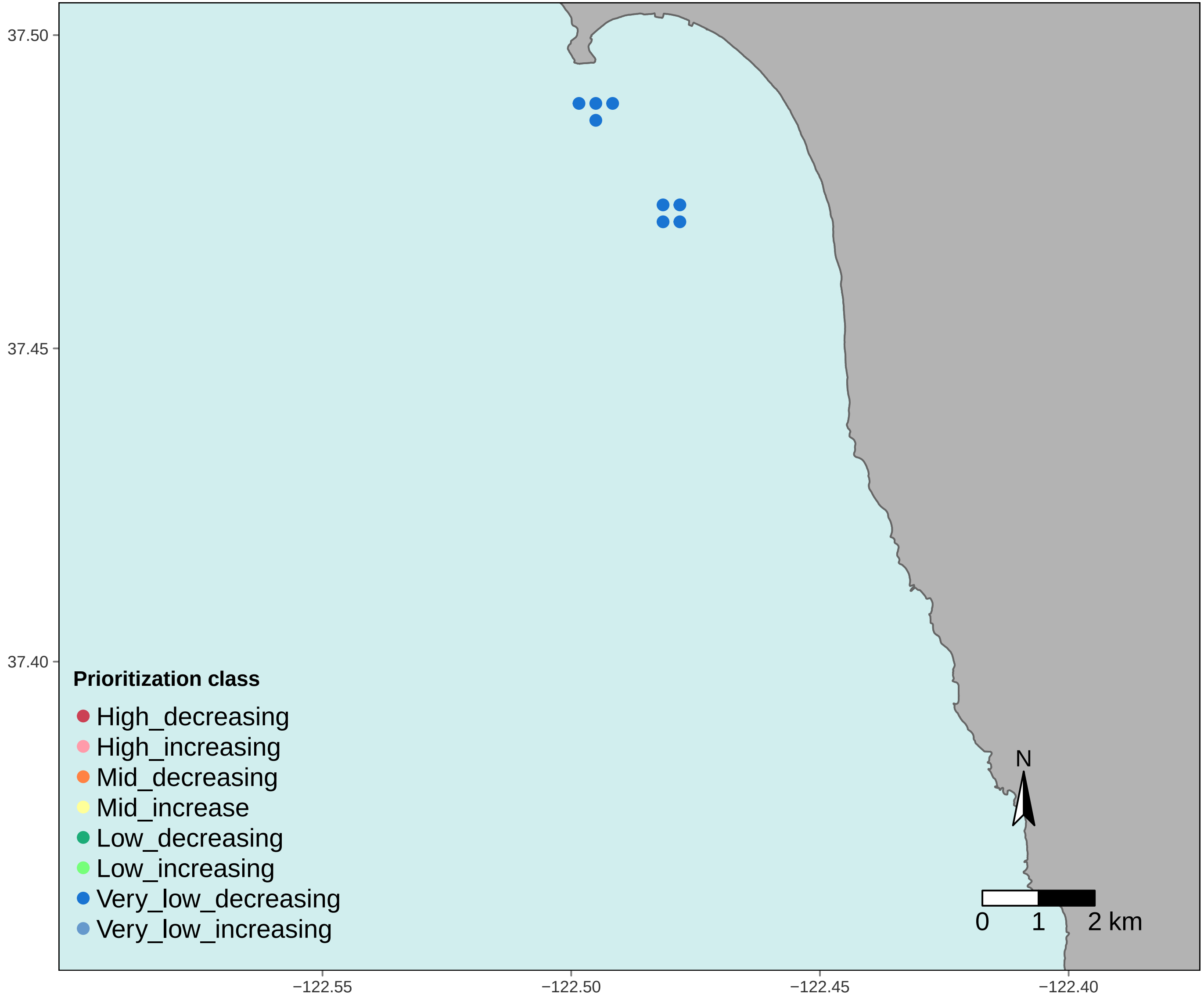

### NC classification_map 32

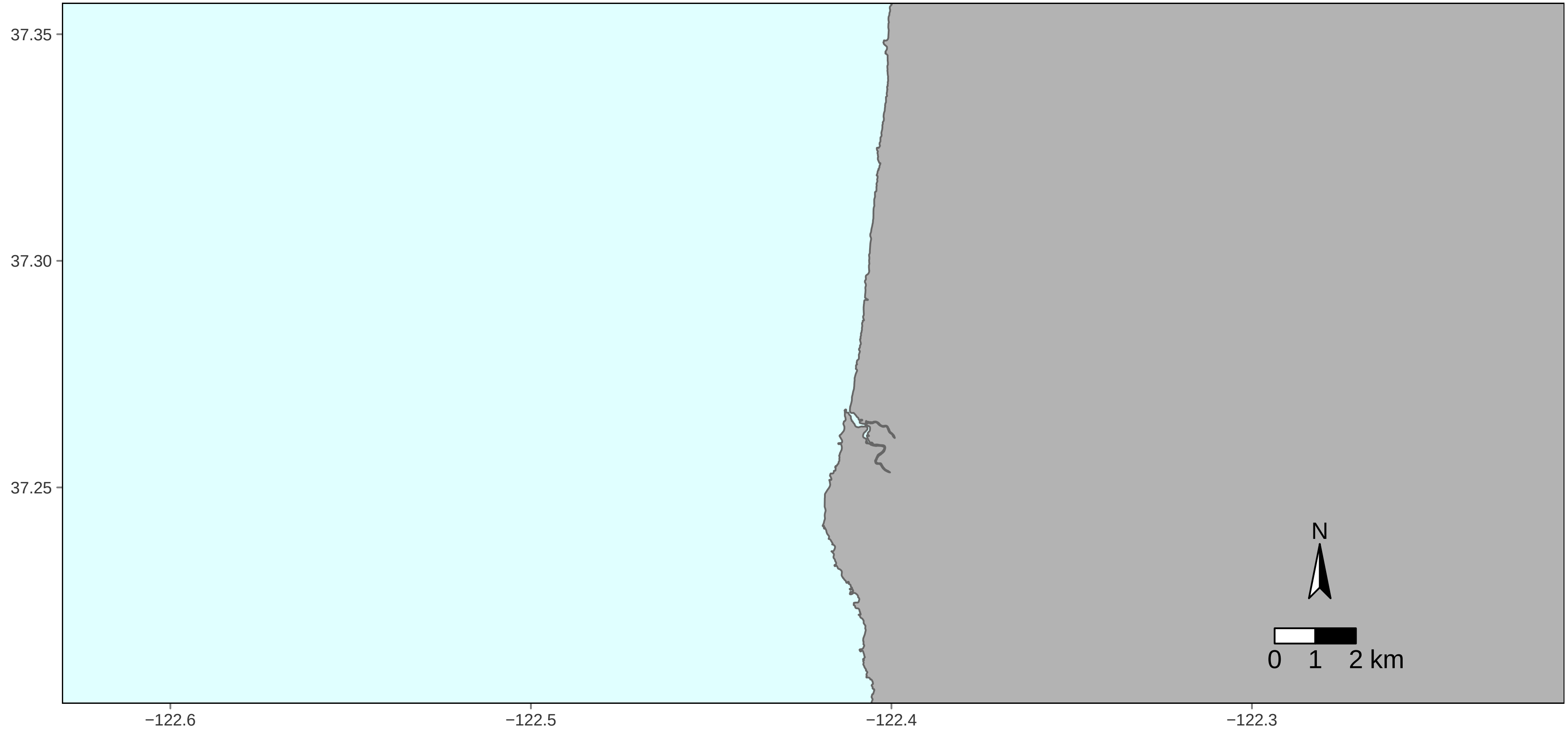

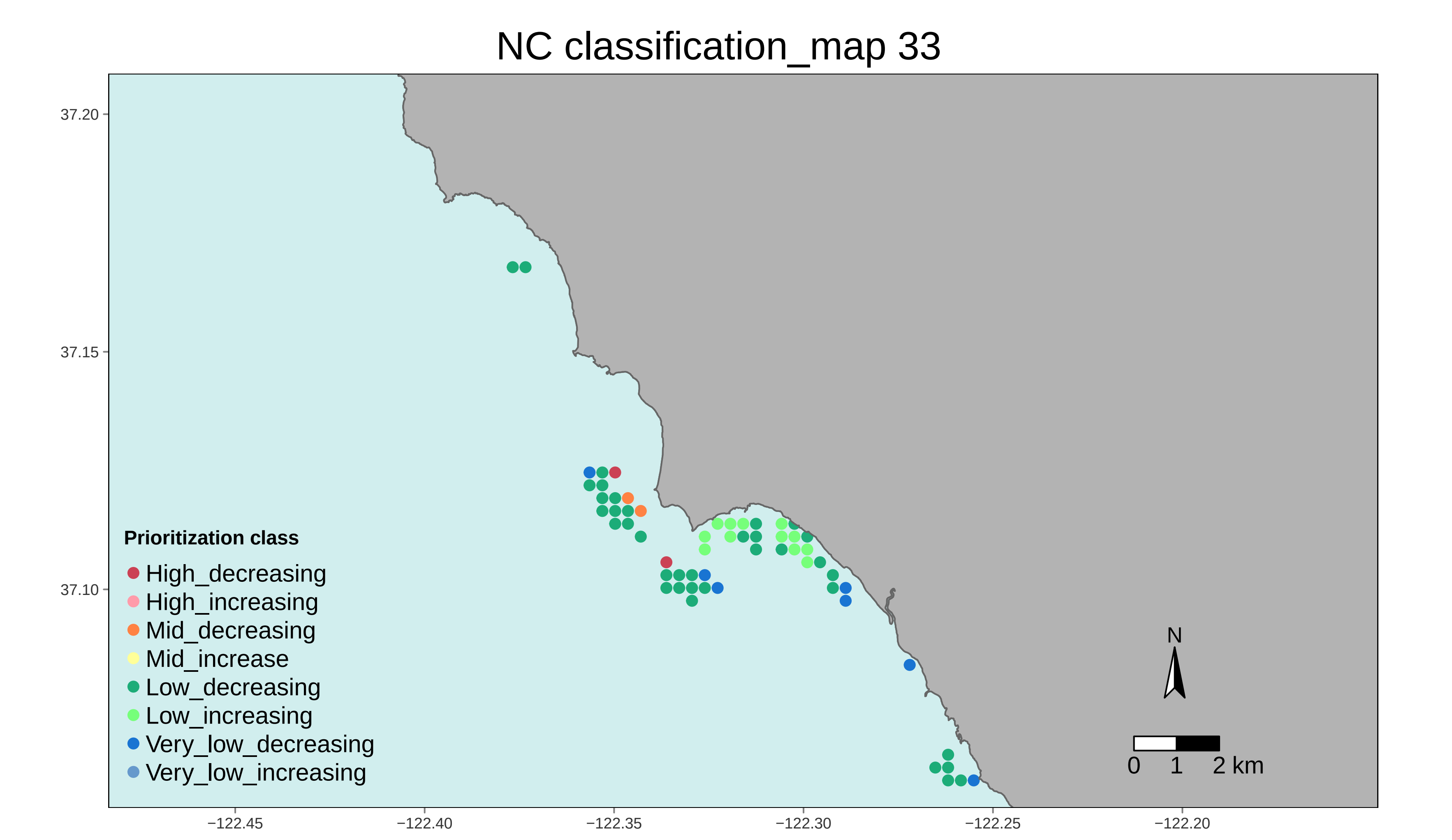
NC classification_map 33
37.20
37.15
37.10
−122.45
−122.40
−122.35
−122.30
−122.25
−122.20
Prioritization class
High_decreasing
High_increasing
Mid_decreasing
Mid_increase
Low_decreasing
Low_increasing
Very_low_decreasing
Very_low_increasing
N
0 1 2 km

### CC classification_map 1

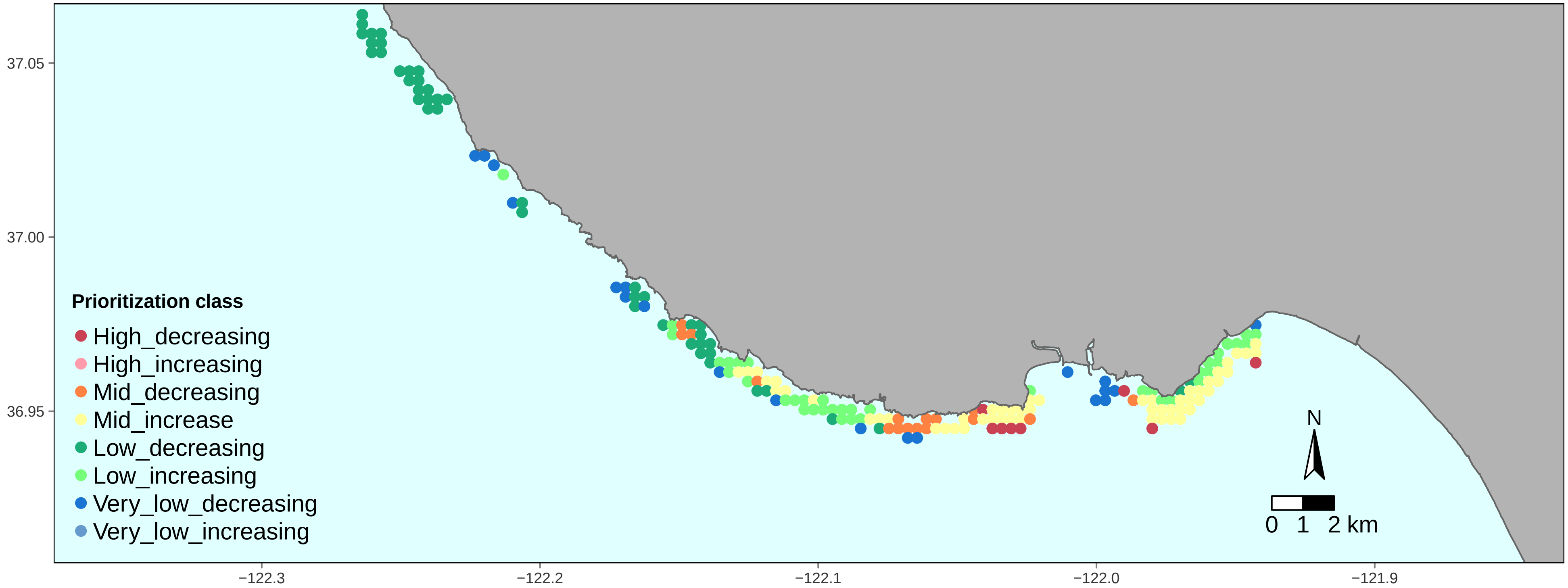

### CC classification_map 2

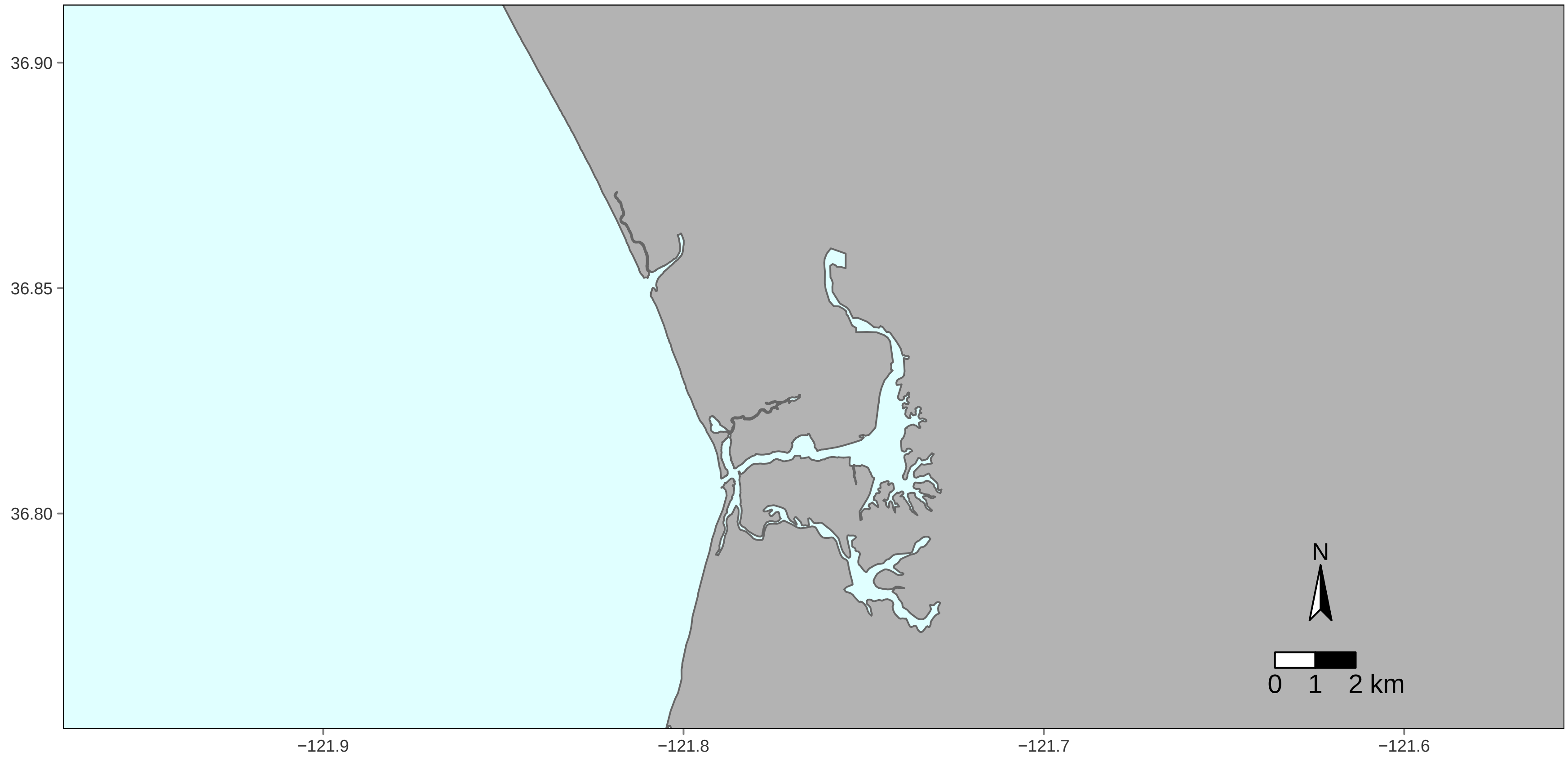

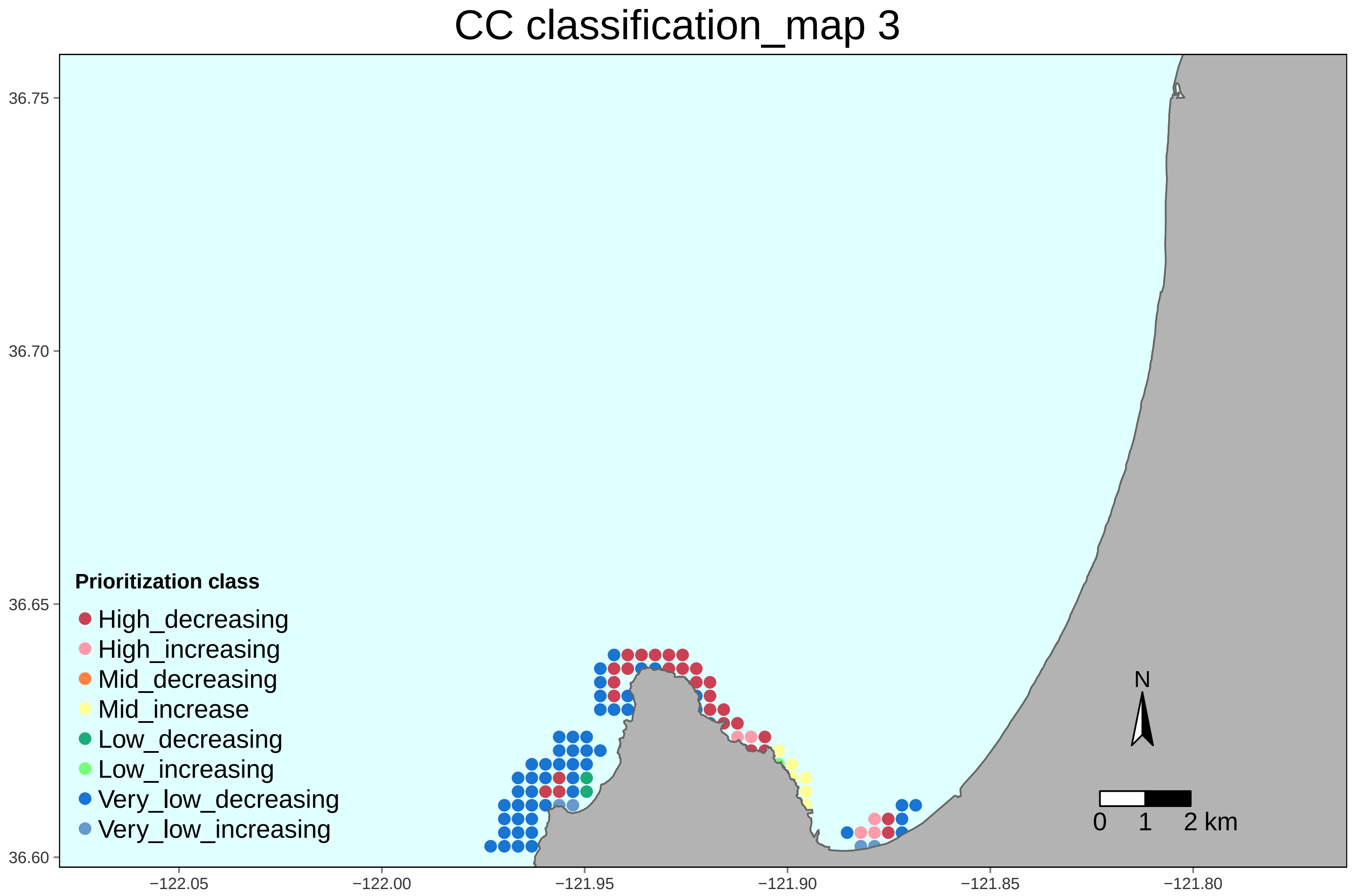
CC classification_map 3
36.75
36.70
36.65
36.60
−122.05
−122.00
−121.95
−121.90
−121.85
−121.80
Prioritization class
High_decreasing
High_increasing
Mid_decreasing
Mid_increase
Low_decreasing
Low_increasing
Very_low_decreasing
Very_low_increasing
N
0
1
2 km

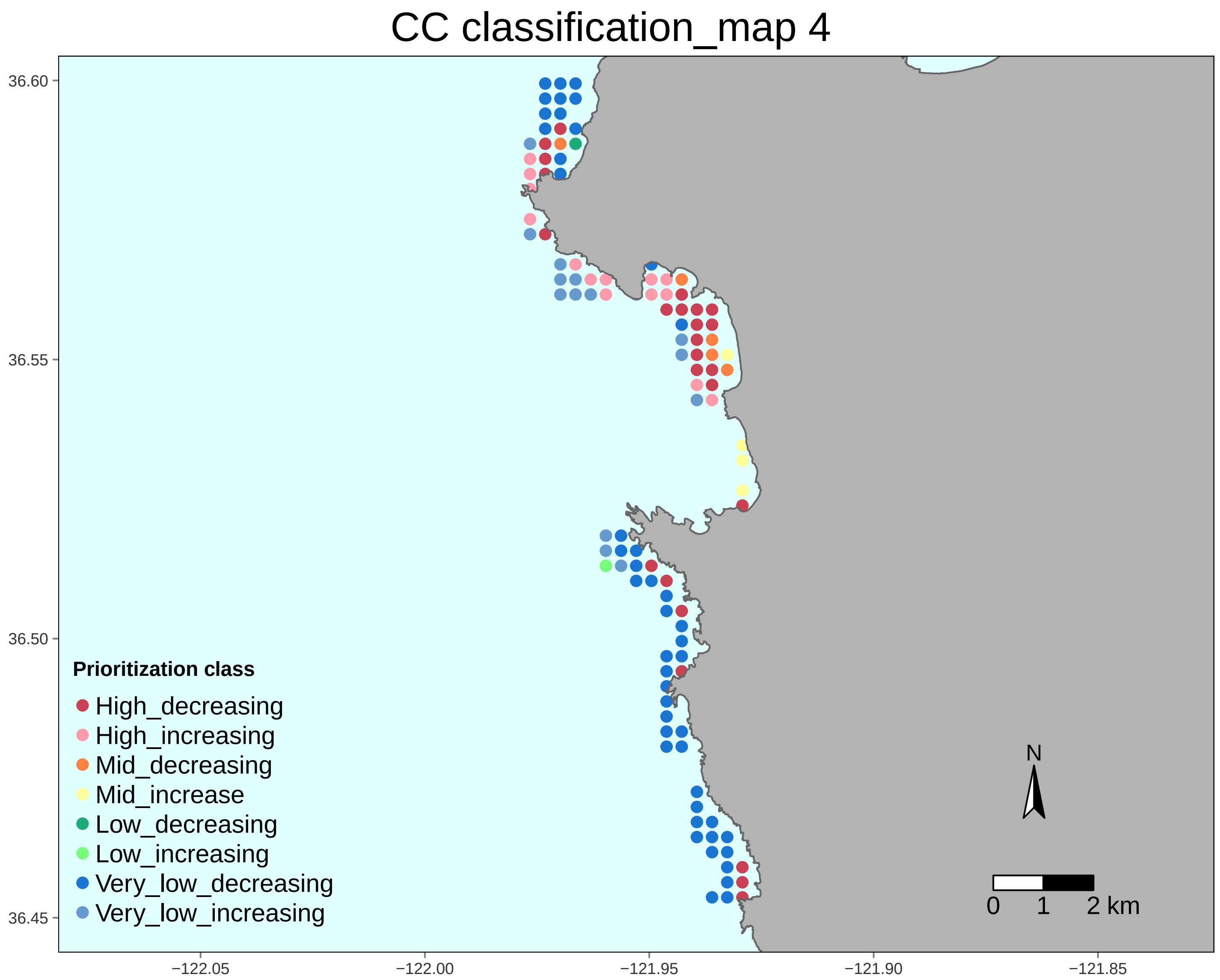
CC classification_map 4
36.60
36.55
36.50
36.45
−122.05
−122.00
−121.95
−121.90
−121.85
Prioritization class
High_decreasing
High_increasing
Mid_decreasing
Mid_increase
Low_decreasing
Low_increasing
Very_low_decreasing
Very_low_increasing
N
0
1
2 km

CC classification_map 5

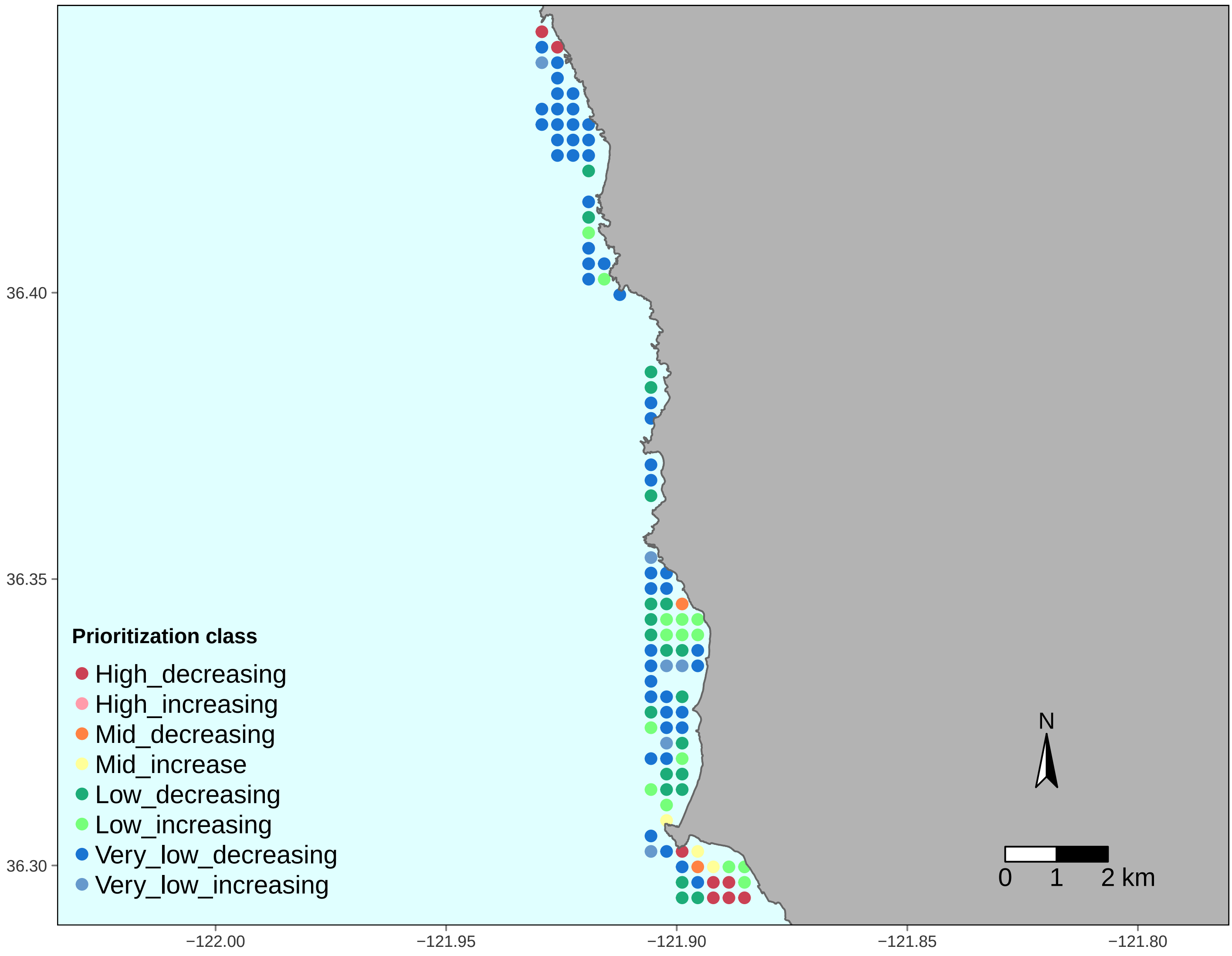

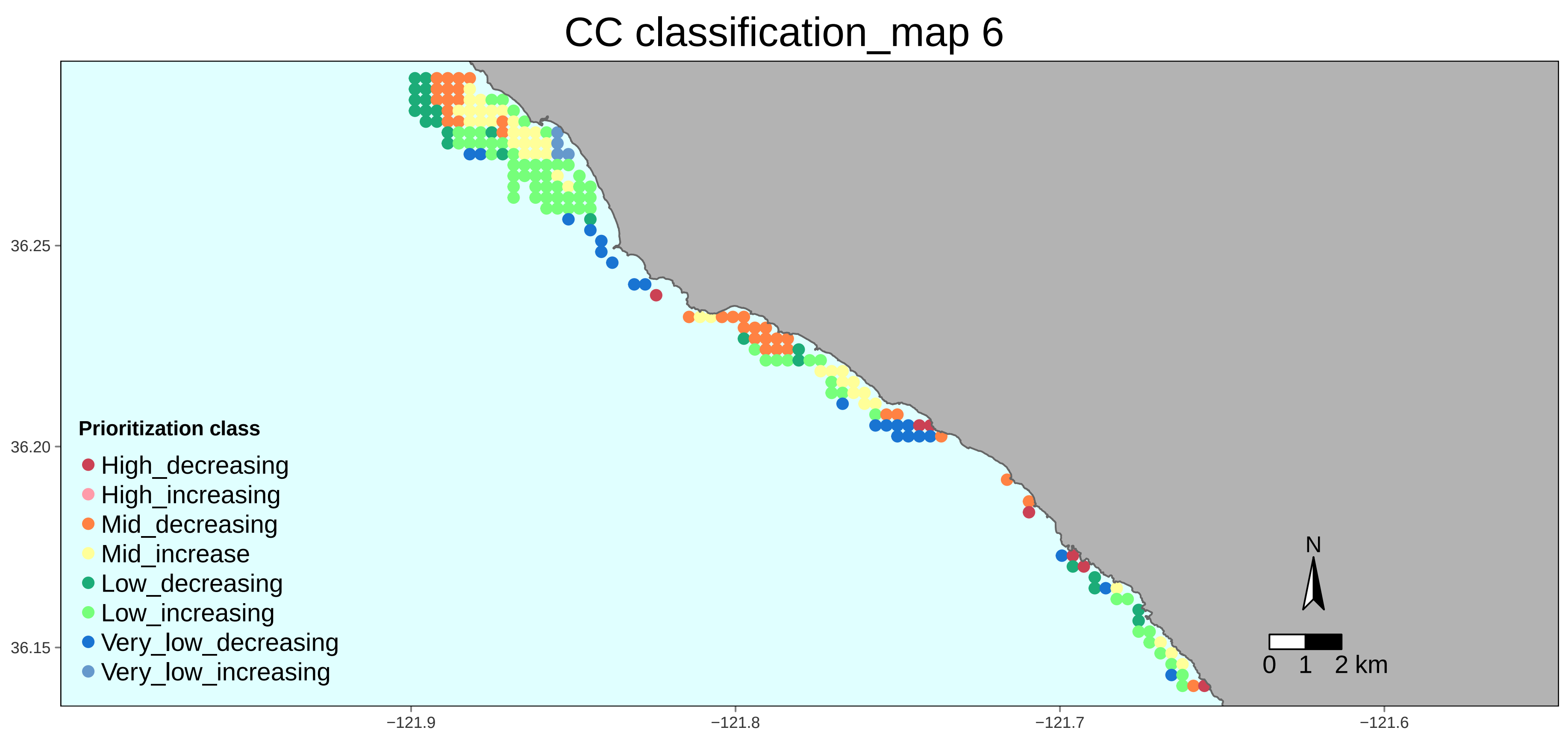
CC classification_map 6
36.25
36.20
36.15
−121.9
−121.8
−121.7
−121.6
Prioritization class
High_decreasing
High_increasing
Mid_decreasing
Mid_increase
Low_decreasing
Low_increasing
Very_low_decreasing
Very_low_increasing
N
0 1 2 km

### CC classification_map 7

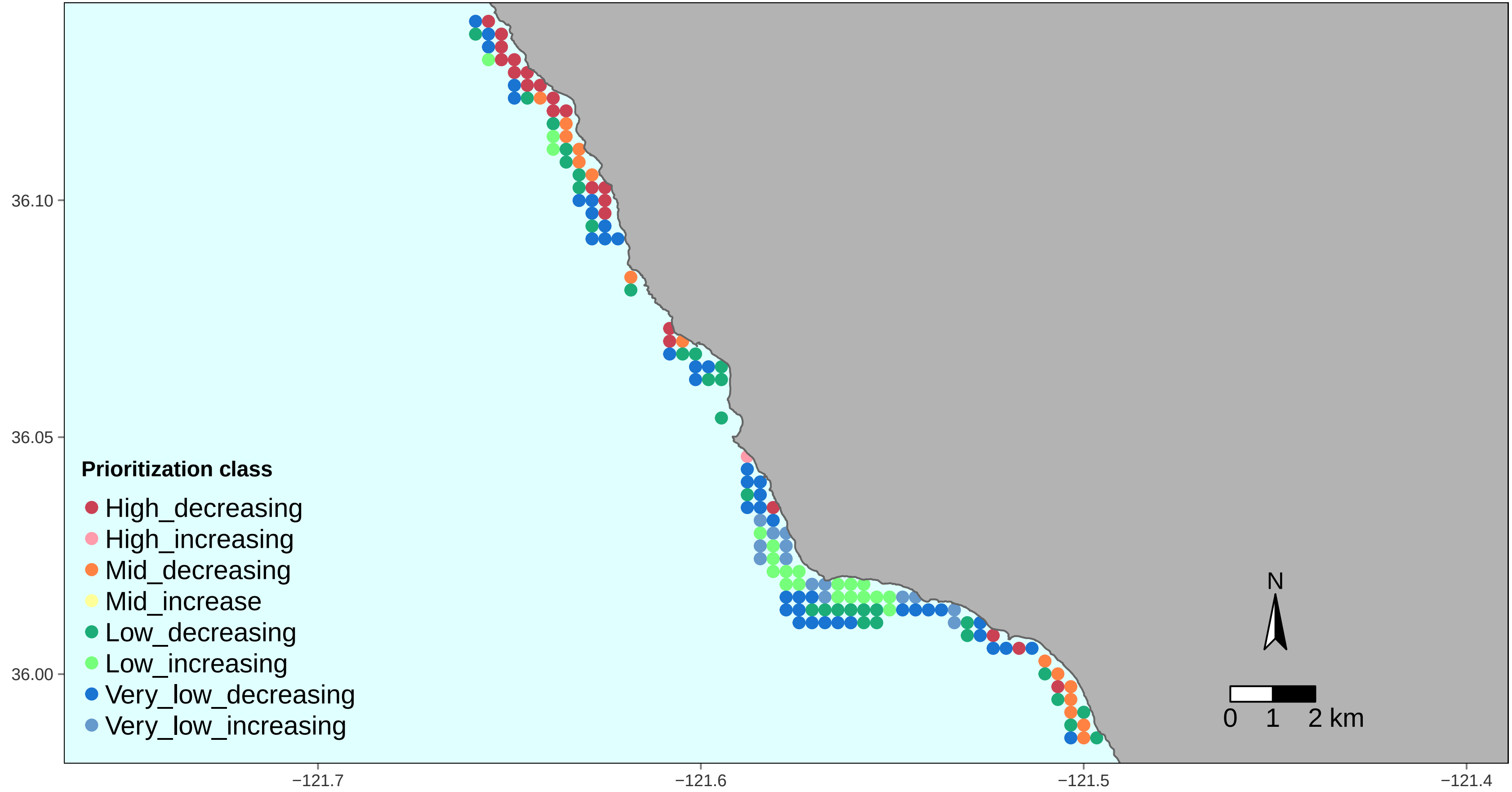

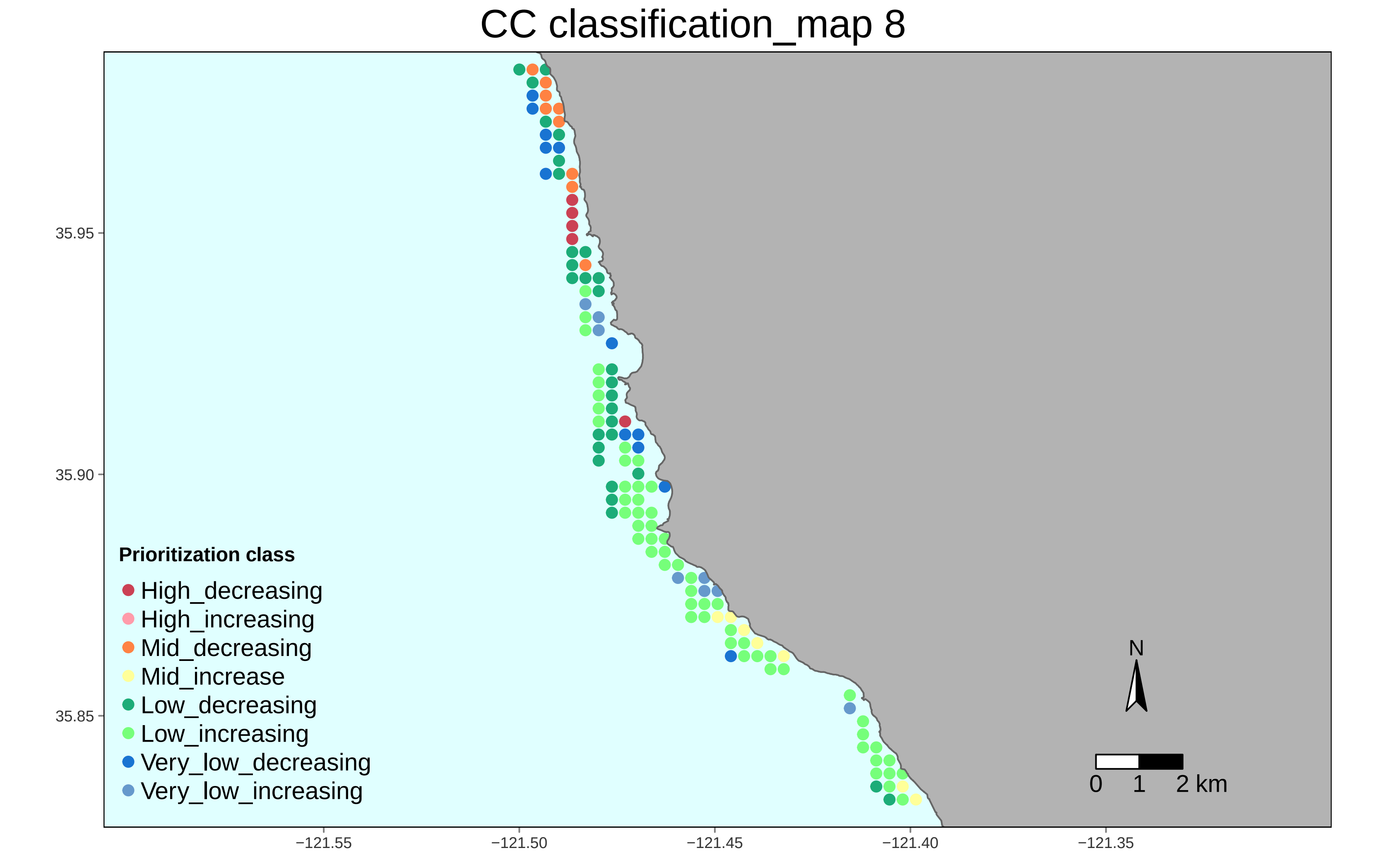

CC classification_map 8
35.95
35.90
35.85
−121.55
−121.50
−121.45
−121.40
−121.35
Prioritization class
High_decreasing
High_increasing
Mid_decreasing
Mid_increase
Low_decreasing
Low_increasing
Very_low_decreasing
Very_low_increasing
N
0 1 2 km

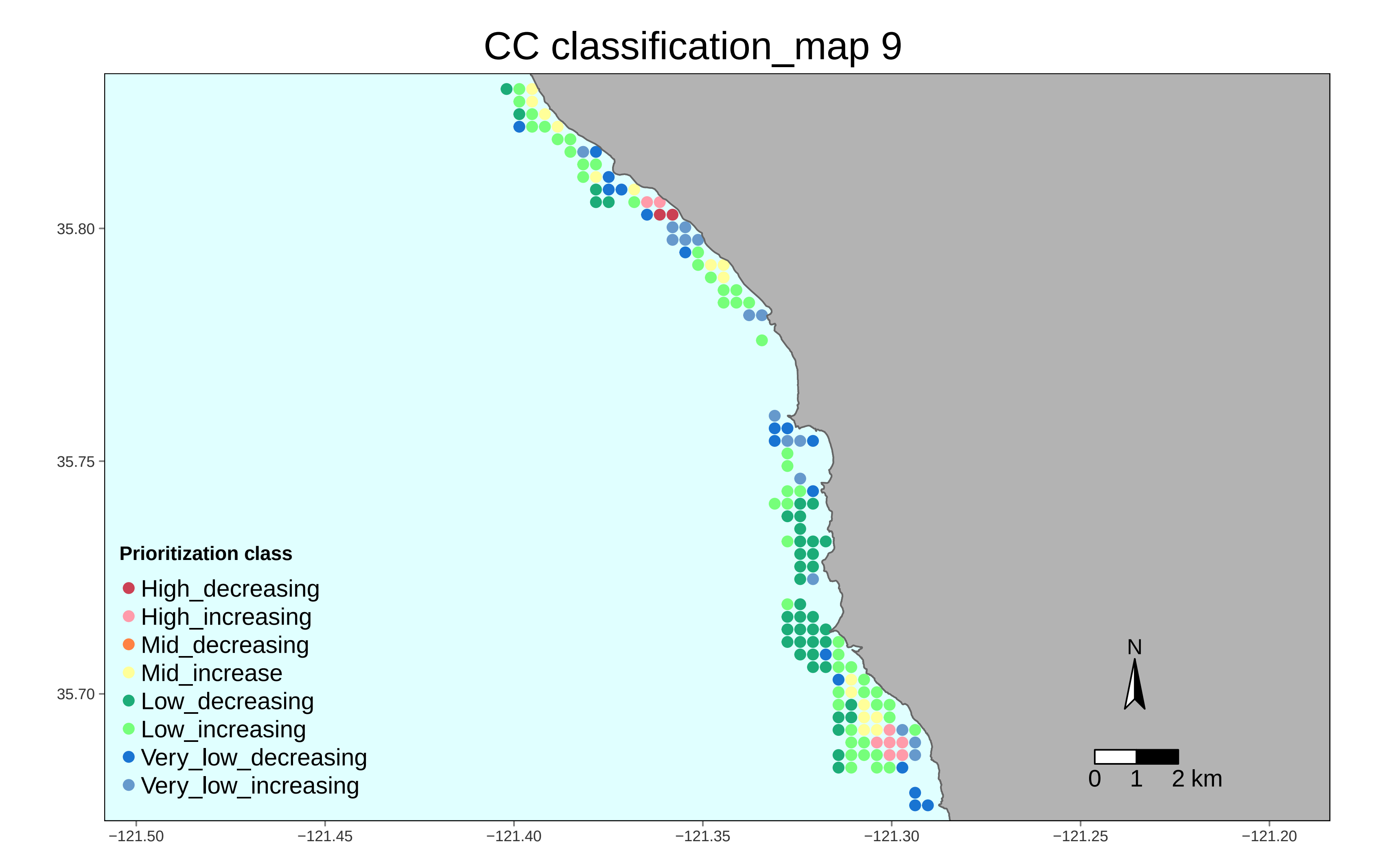
CC classification_map 9
Prioritization class
High_decreasing
High_increasing
Mid_decreasing
Mid_increase
Low_decreasing
Low_increasing
Very_low_decreasing
Very_low_increasing
N
0
1
2 km
35.80
35.75
35.70
−121.50
−121.45
−121.40
−121.35
−121.30
−121.25
−121.20

### CC classification_map 10

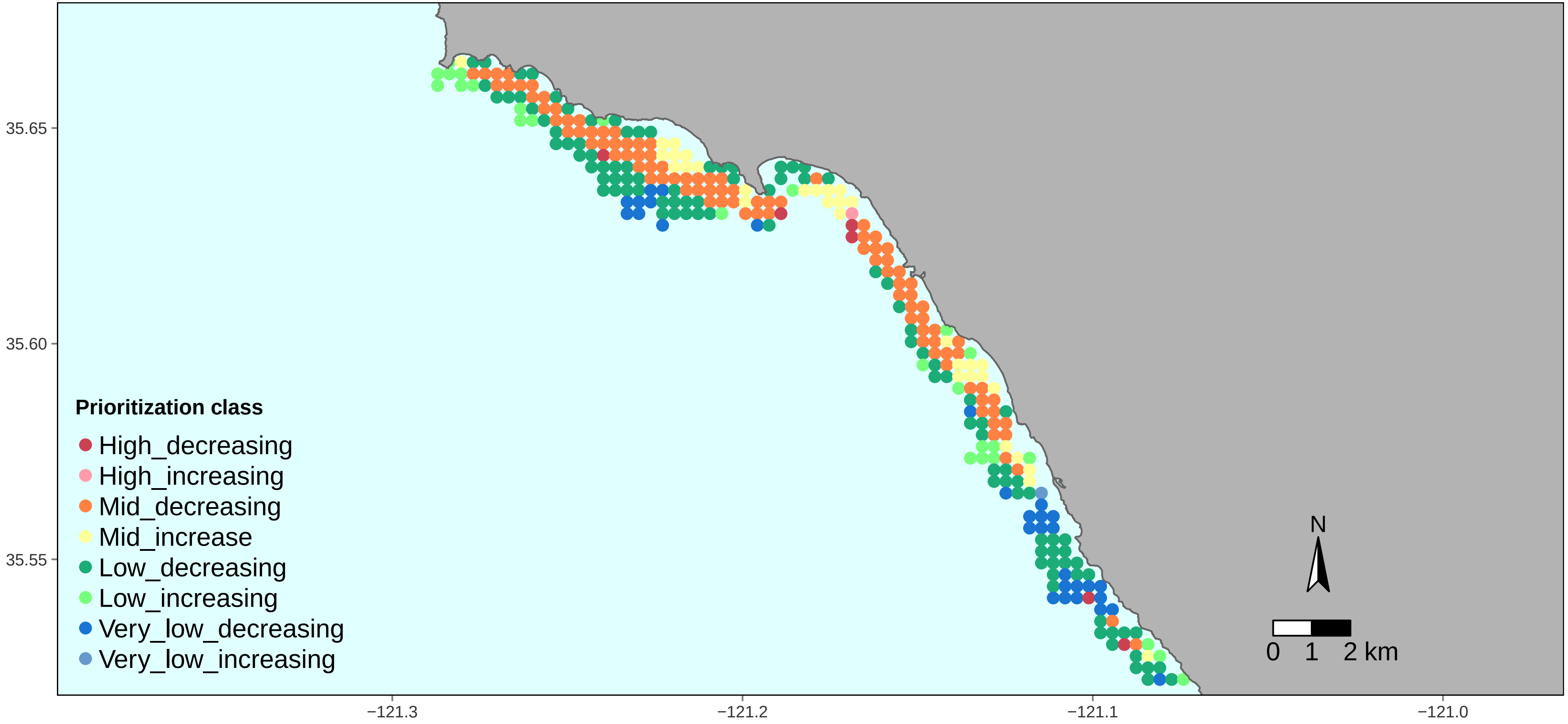

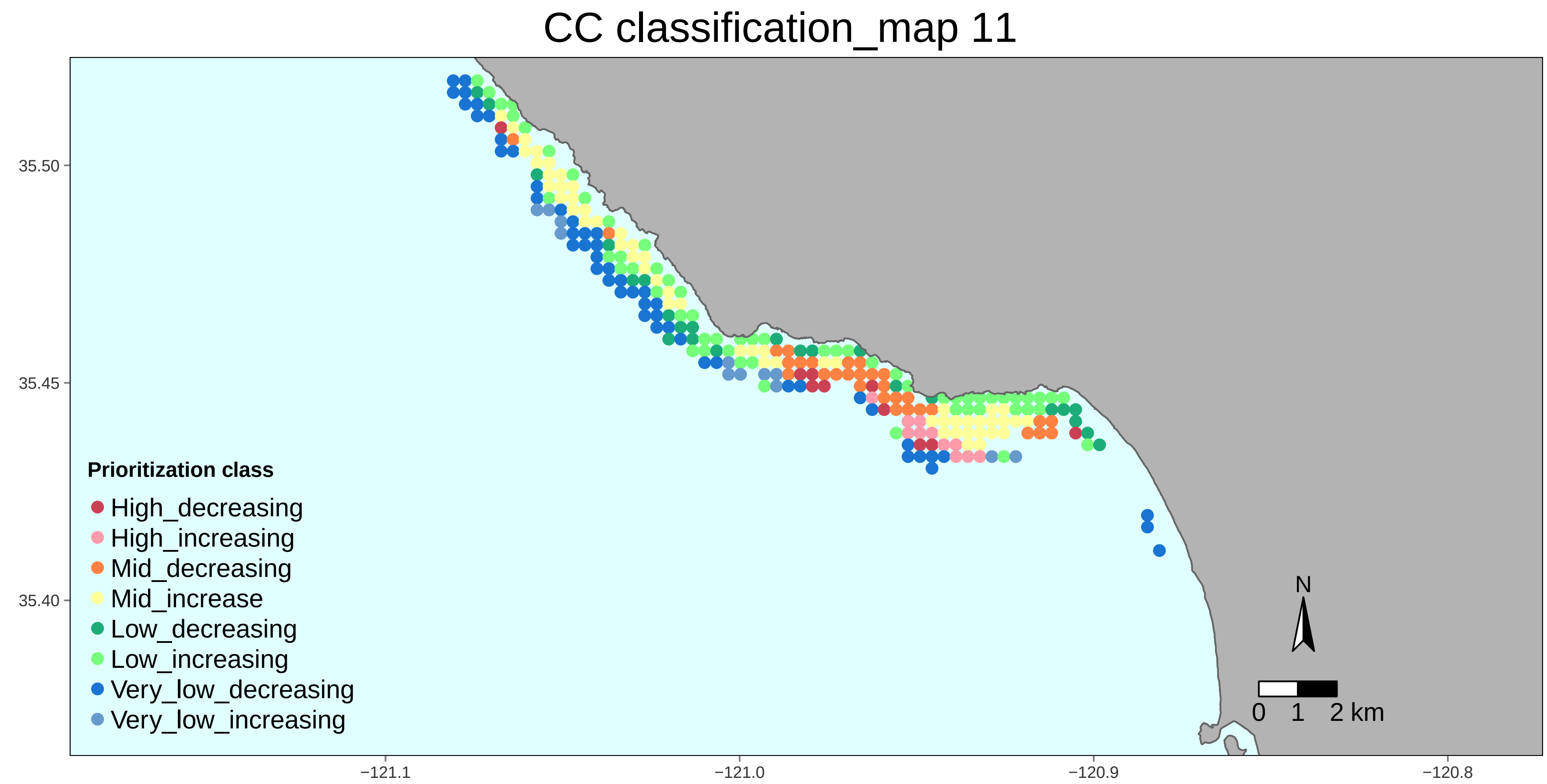
CC classification_map 11
35.50
35.45
35.40
−121.1
−121.0
−120.9
−120.8
Prioritization class
High_decreasing
High_increasing
Mid_decreasing
Mid_increase
Low_decreasing
Low_increasing
Very_low_decreasing
Very_low_increasing
N
0
1
2 km

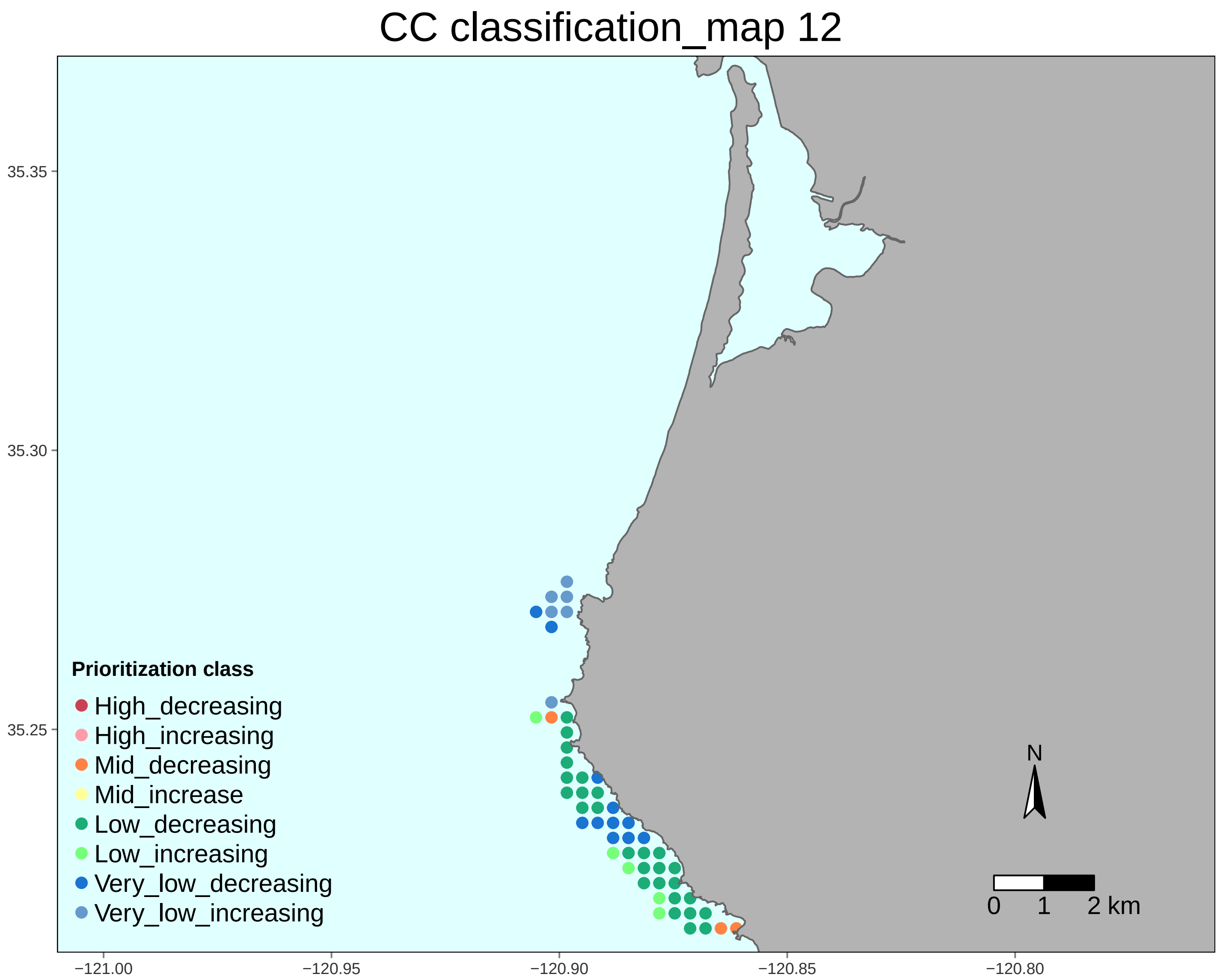
CC classification_map 12
35.35
35.30
35.25
−121.00
−120.95
−120.90
−120.85
−120.80
Prioritization class
High_decreasing
High_increasing
Mid_decreasing
Mid_increase
Low_decreasing
Low_increasing
Very_low_decreasing
Very_low_increasing
N
0 1 2 km

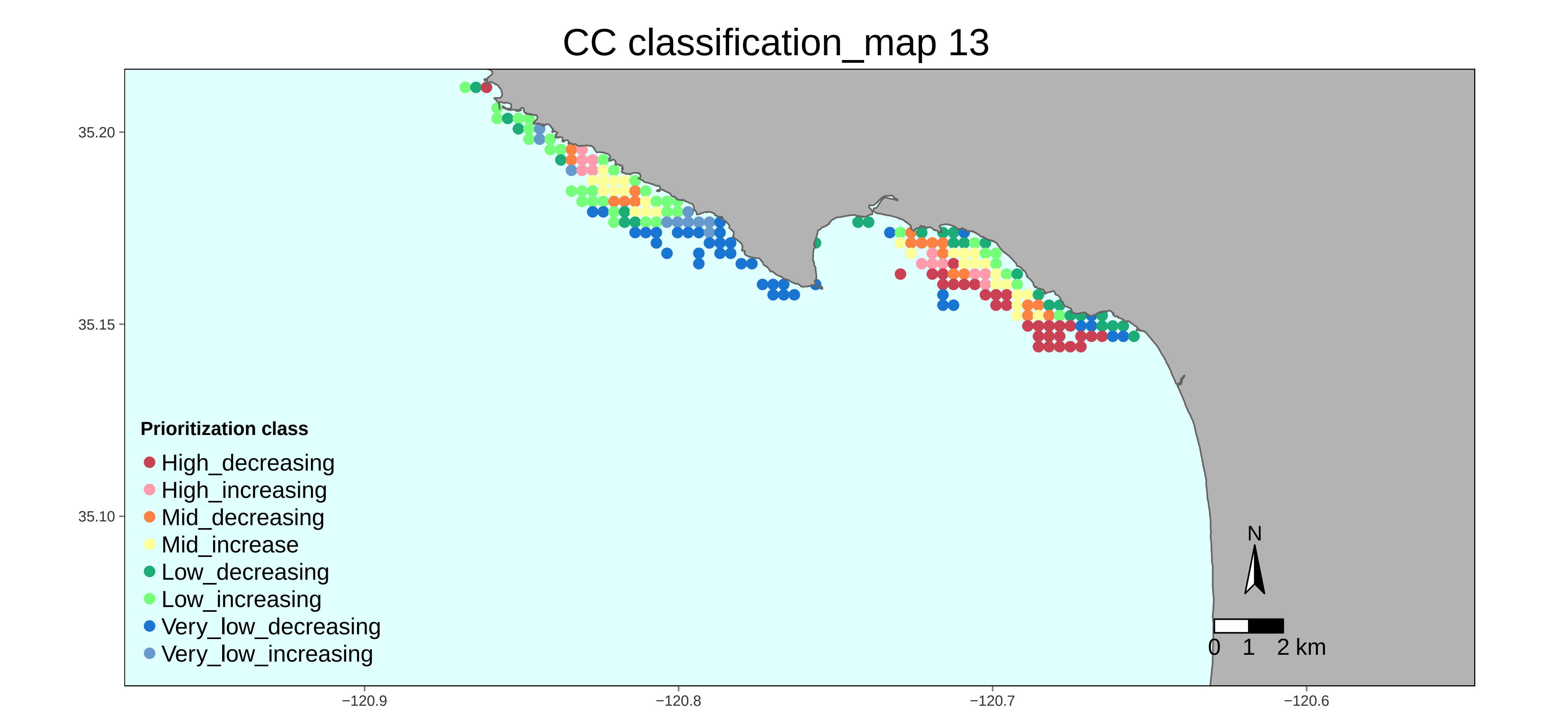
CC classification_map 13
35.20
35.15
35.10
−120.9
−120.8
−120.7
−120.6
Prioritization class
High_decreasing
High_increasing
Mid_decreasing
Mid_increase
Low_decreasing
Low_increasing
Very_low_decreasing
Very_low_increasing
N
0 1 2 km

CC classification_map 14

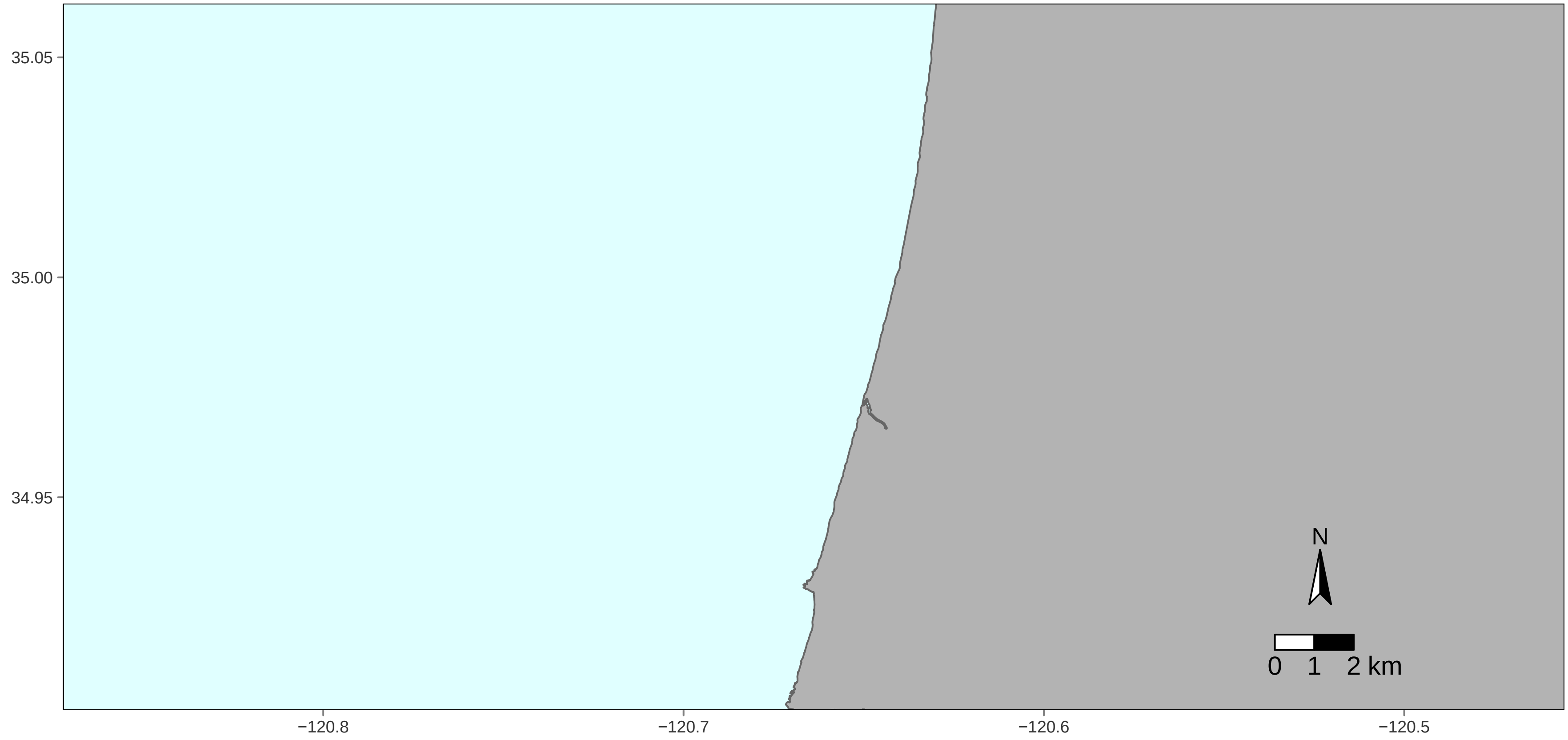

### CC classification_map 15

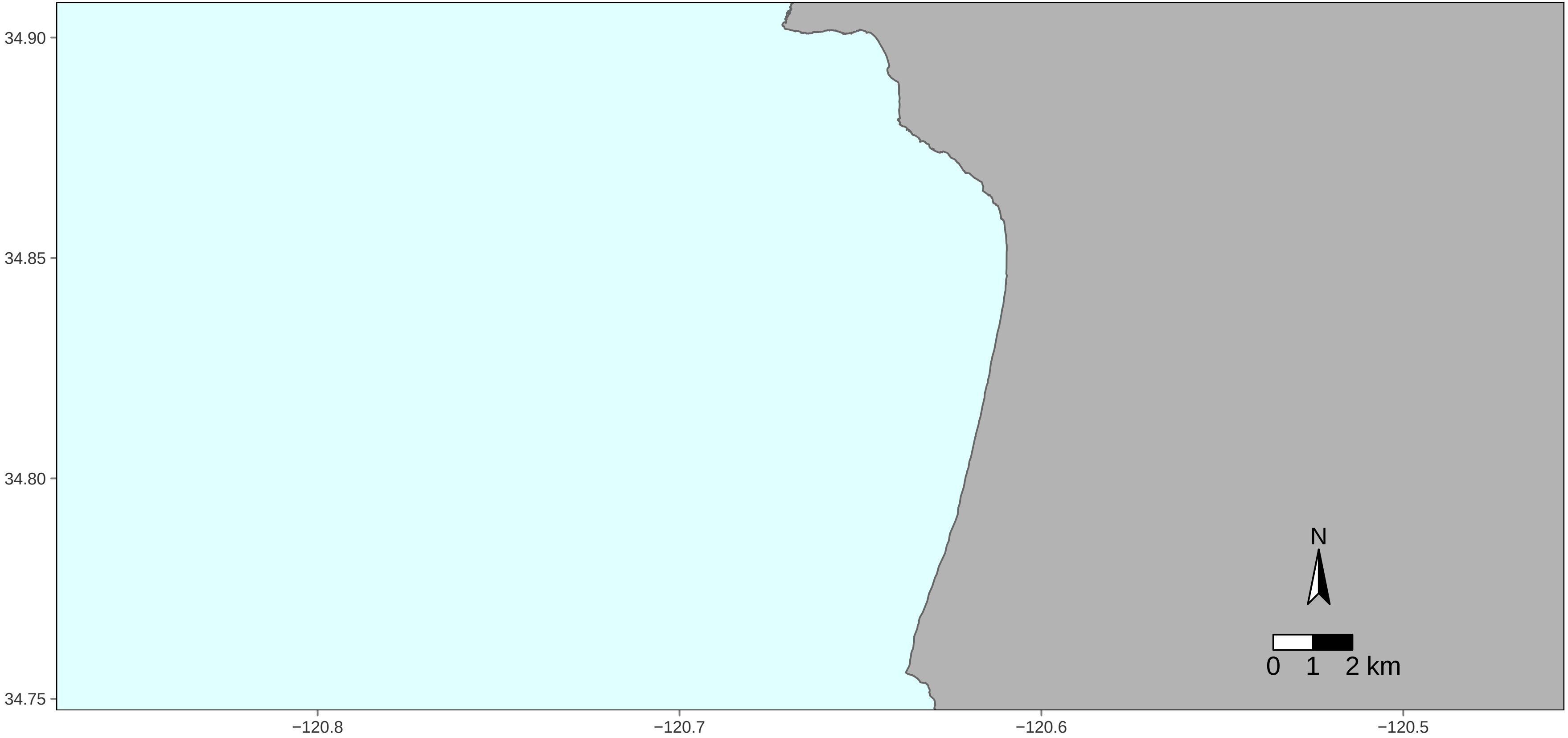

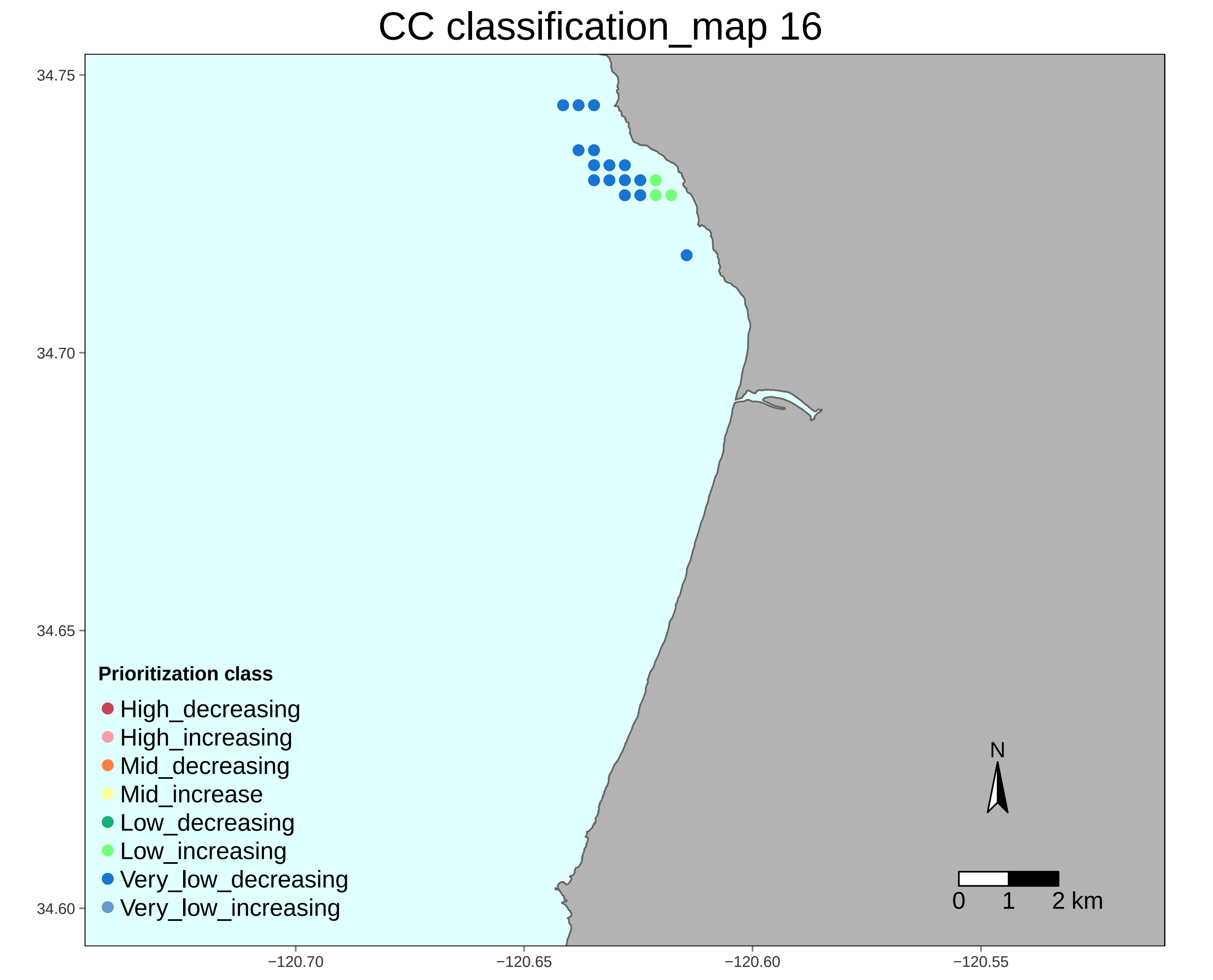

CC classification_map 16
34.75
34.70
34.65
34.60
−120.70
−120.65
−120.60
−120.55
Prioritization class
High_decreasing
High_increasing
Mid_decreasing
Mid_increase
Low_decreasing
Low_increasing
Very_low_decreasing
Very_low_increasing
N
0 1 2 km

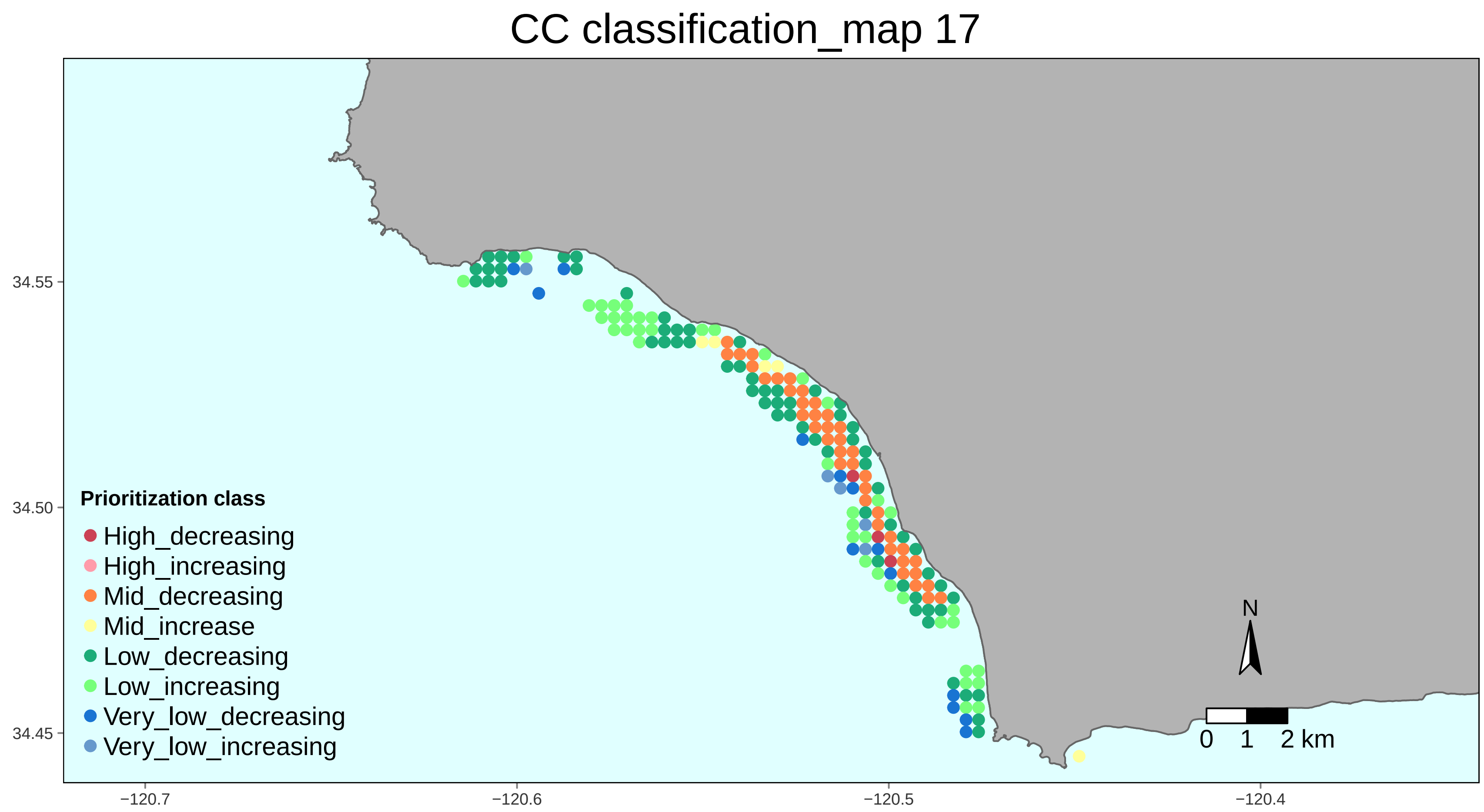

CC classification_map 17
34.55
34.50
34.45
−120.7
−120.6
−120.5
−120.4
Prioritization class
High_decreasing
High_increasing
Mid_decreasing
Mid_increase
Low_decreasing
Low_increasing
Very_low_decreasing
Very_low_increasing
N
0
1
2 km

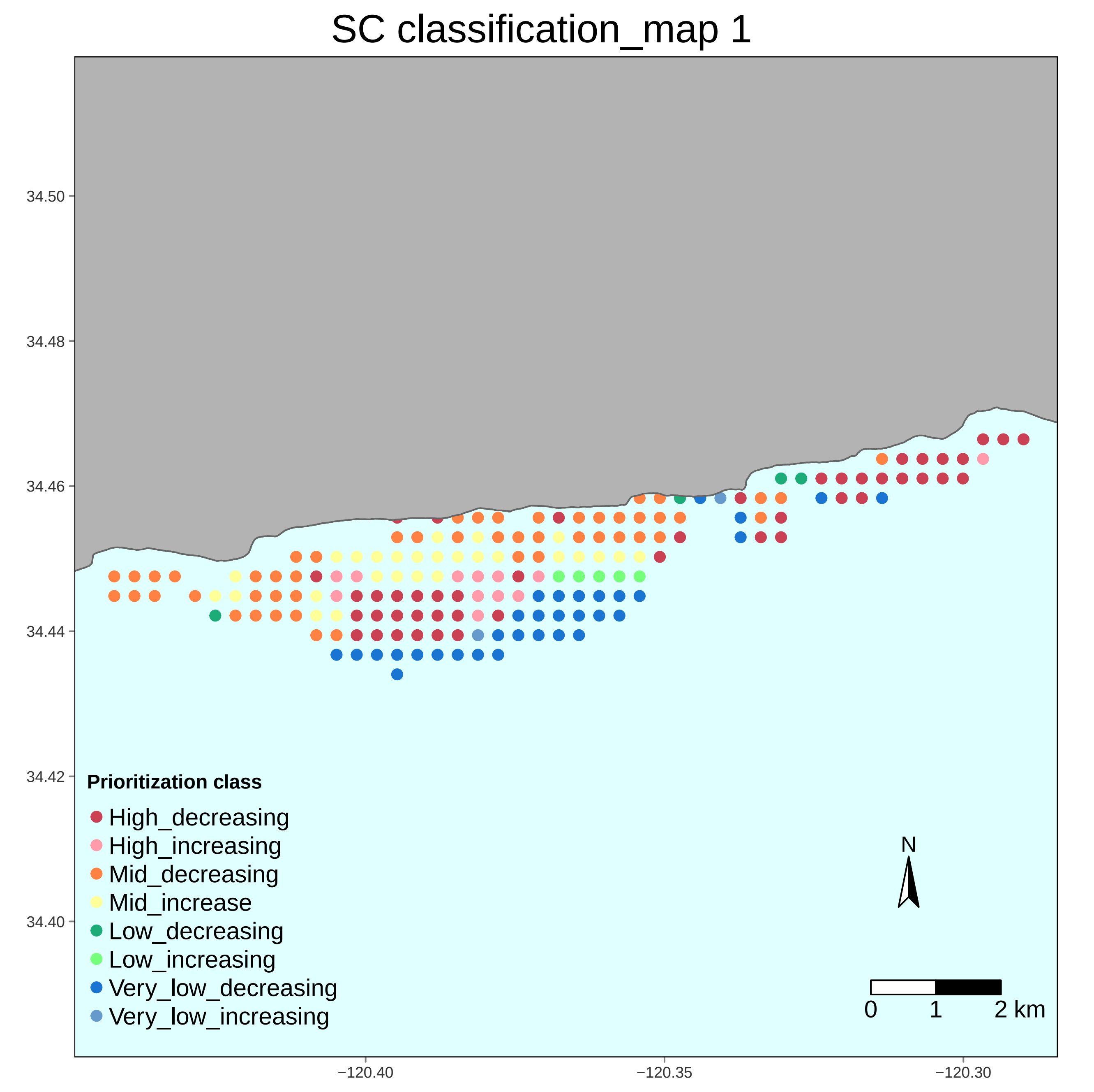

SC classification_map 1
34.50
34.48
34.46
34.44
34.42
34.40
−120.40
−120.35
−120.30
Prioritization class
High_decreasing
High_increasing
Mid_decreasing
Mid_increase
Low_decreasing
Low_increasing
Very_low_decreasing
Very_low_increasing
N
0
1
2 km

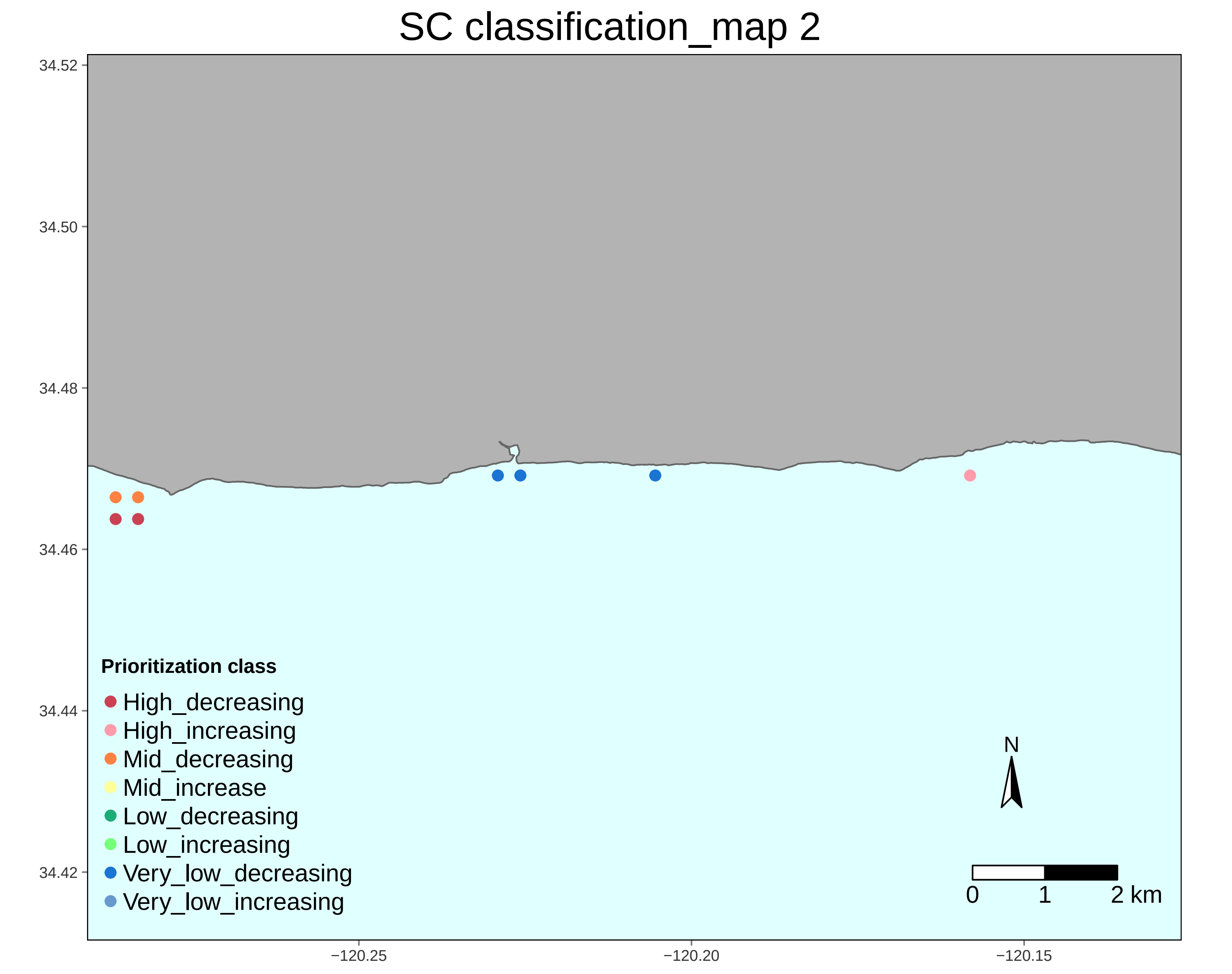
SC classification_map 2
34.52
34.50
34.48
34.46
34.44
34.42
−120.25
−120.20
−120.15
Prioritization class
High_decreasing
High_increasing
Mid_decreasing
Mid_increase
Low_decreasing
Low_increasing
Very_low_decreasing
Very_low_increasing
N
0
1
2 km

### SC classification_map 3

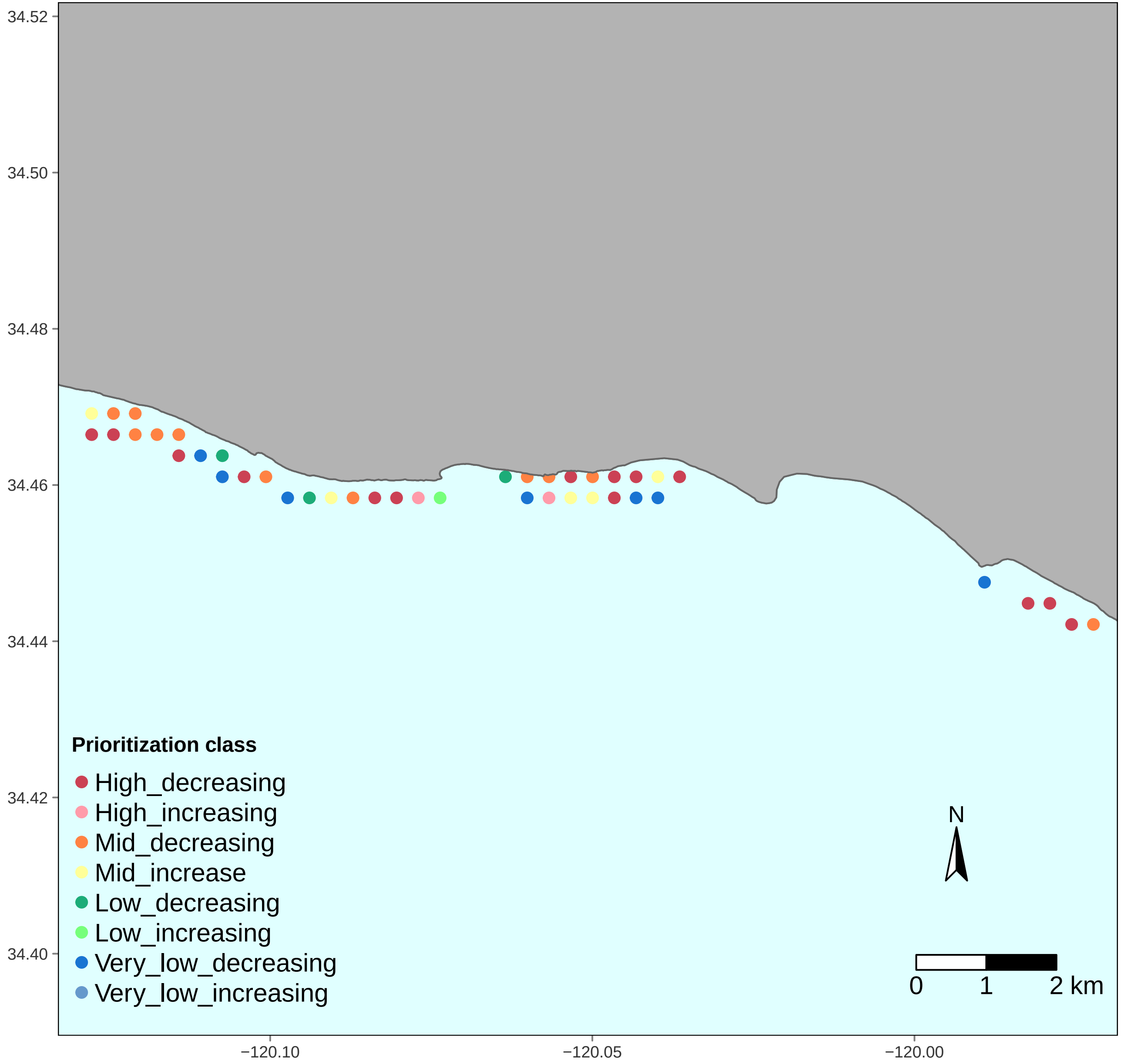

### SC classification_map 4

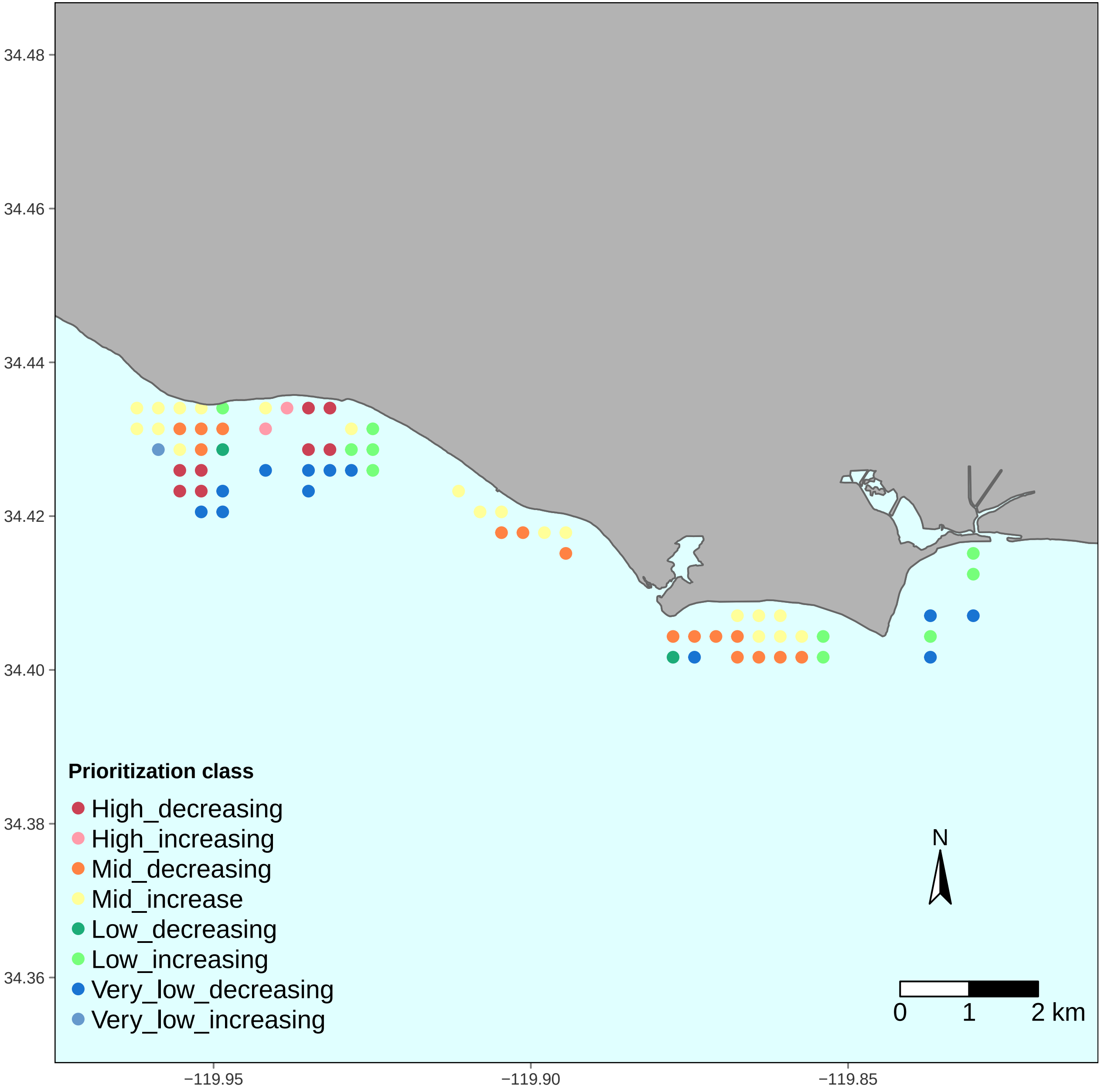

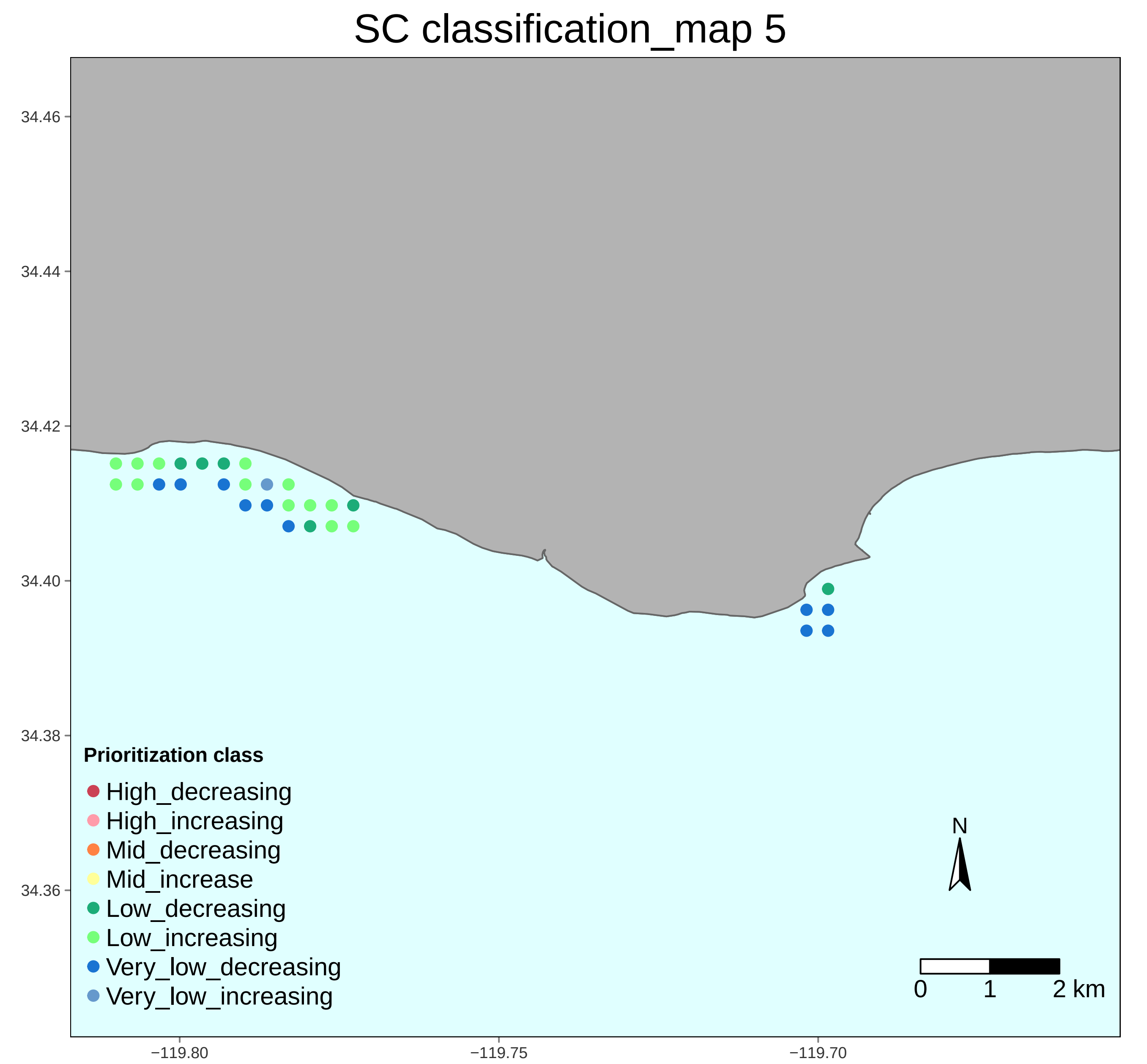

SC classification_map 5
34.46
34.44
34.42
34.40
34.38
34.36
−119.80
−119.75
−119.70
Prioritization class
High_decreasing
High_increasing
Mid_decreasing
Mid_increase
Low_decreasing
Low_increasing
Very_low_decreasing
Very_low_increasing
N
0
1
2 km

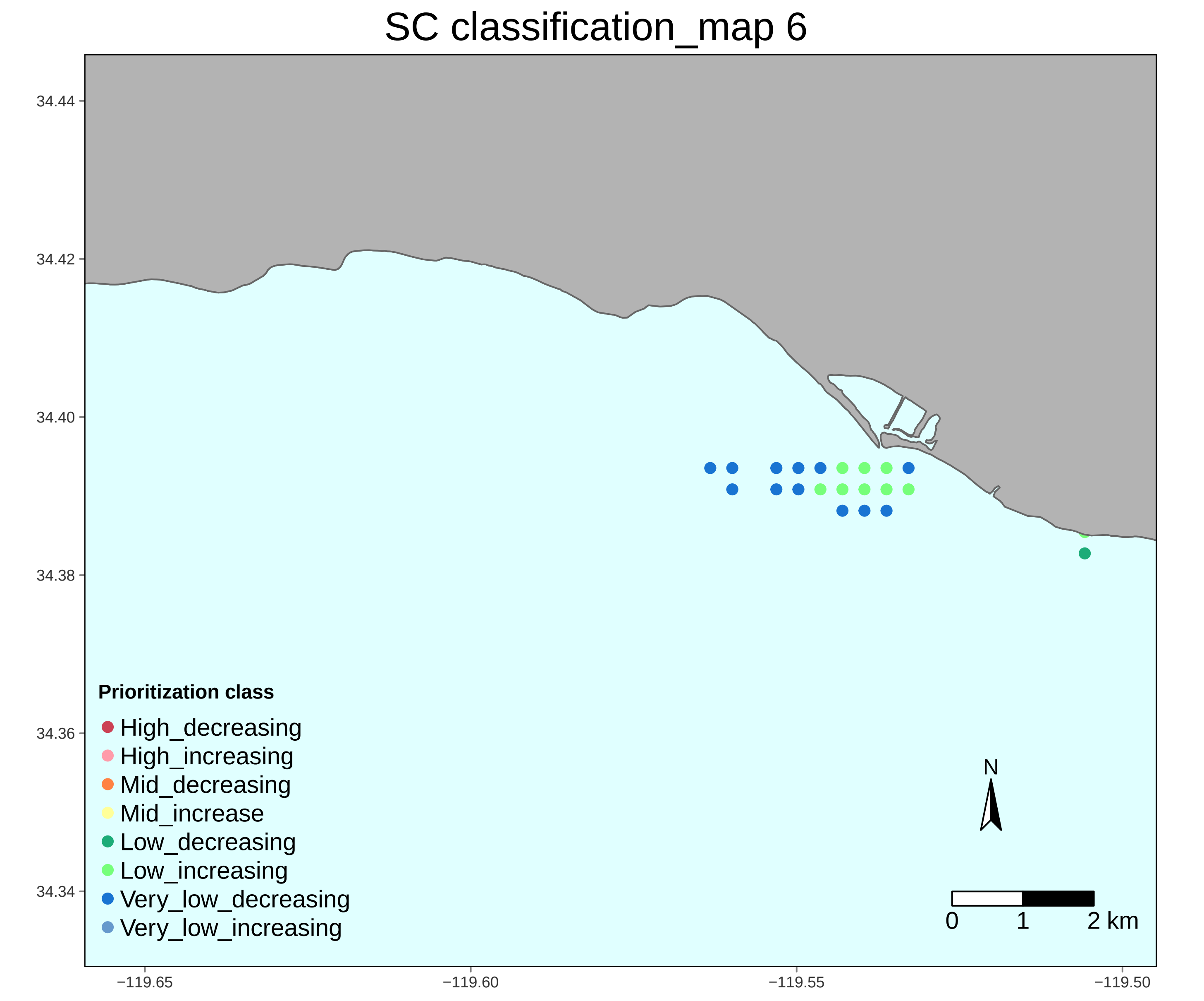
SC classification_map 6
34.44
34.42
34.40
34.38
34.36
34.34
−119.65
−119.60
−119.55
−119.50
Prioritization class
High_decreasing
High_increasing
Mid_decreasing
Mid_increase
Low_decreasing
Low_increasing
Very_low_decreasing
Very_low_increasing
N
0
1
2 km

### SC classification_map 7

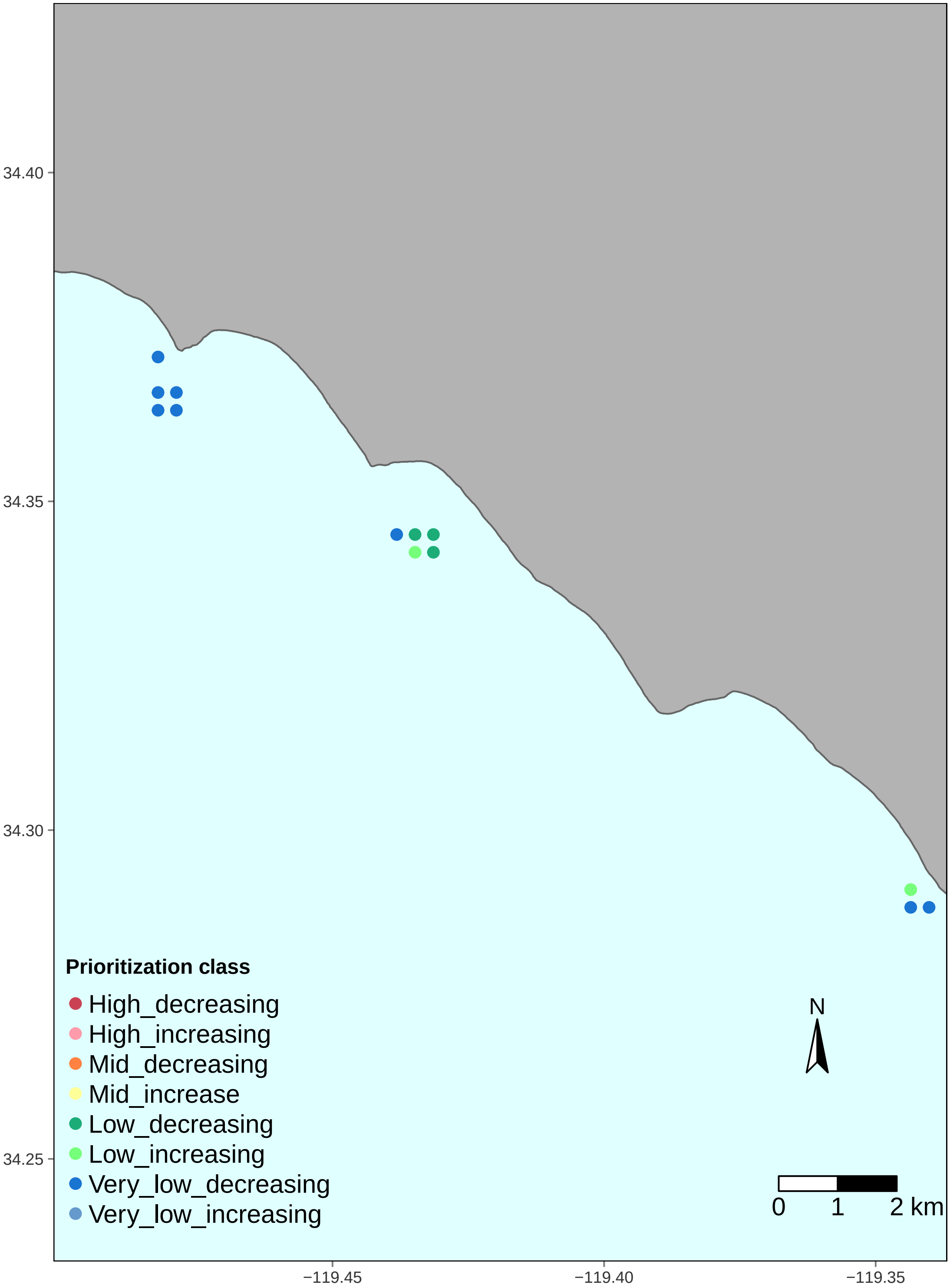

### SC classification_map 8

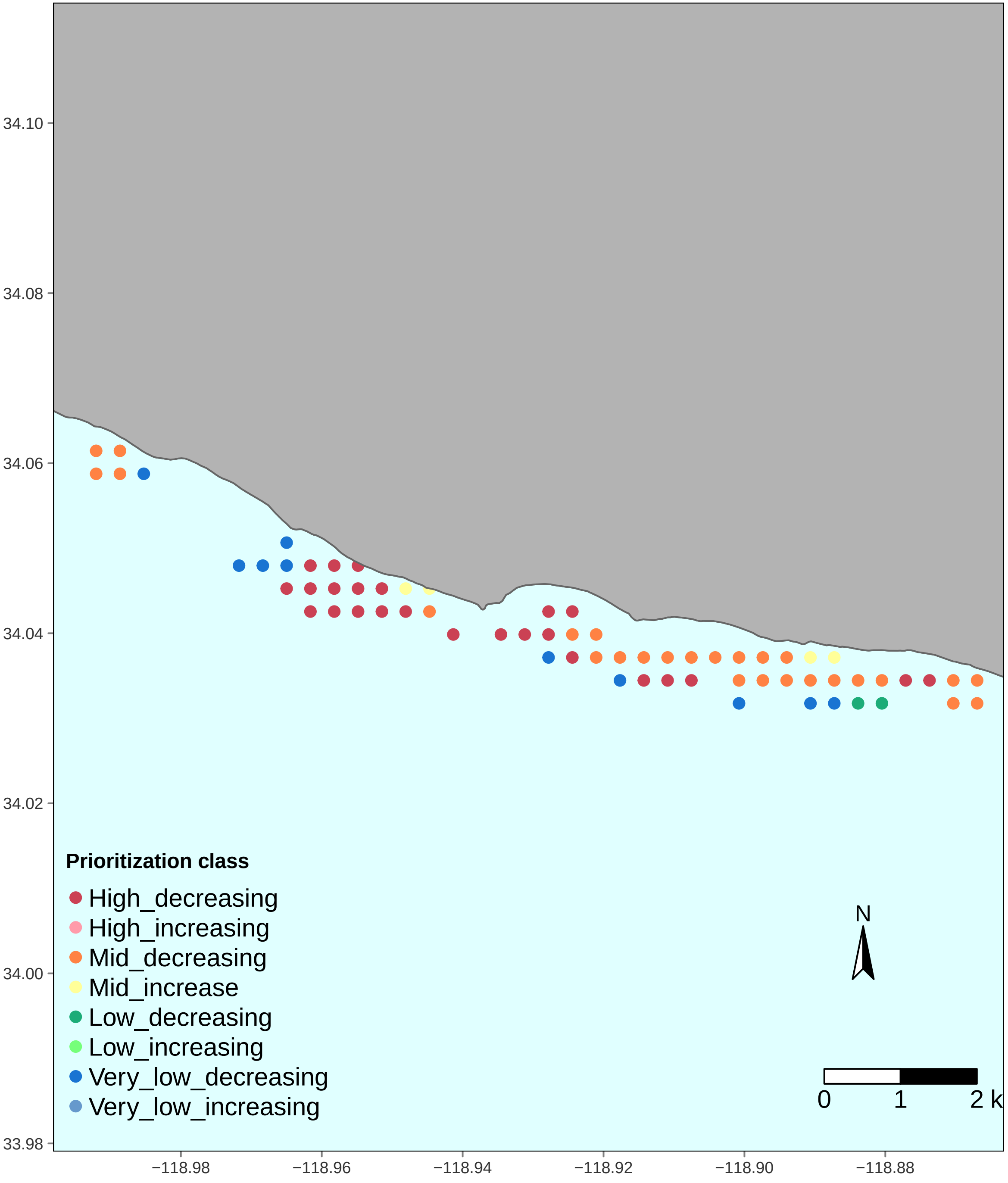

### SC classification_map 9

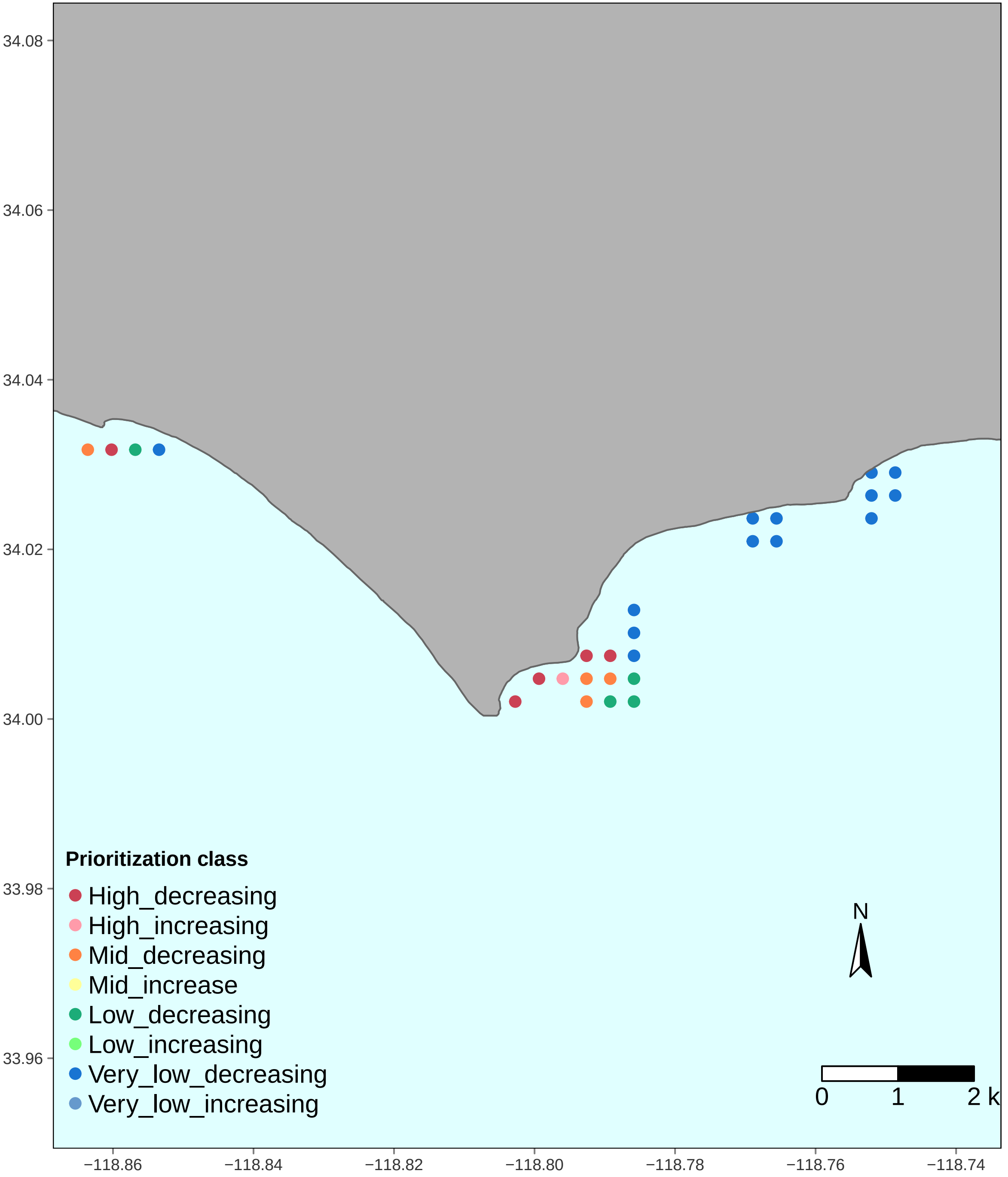

### SC classification_map 10

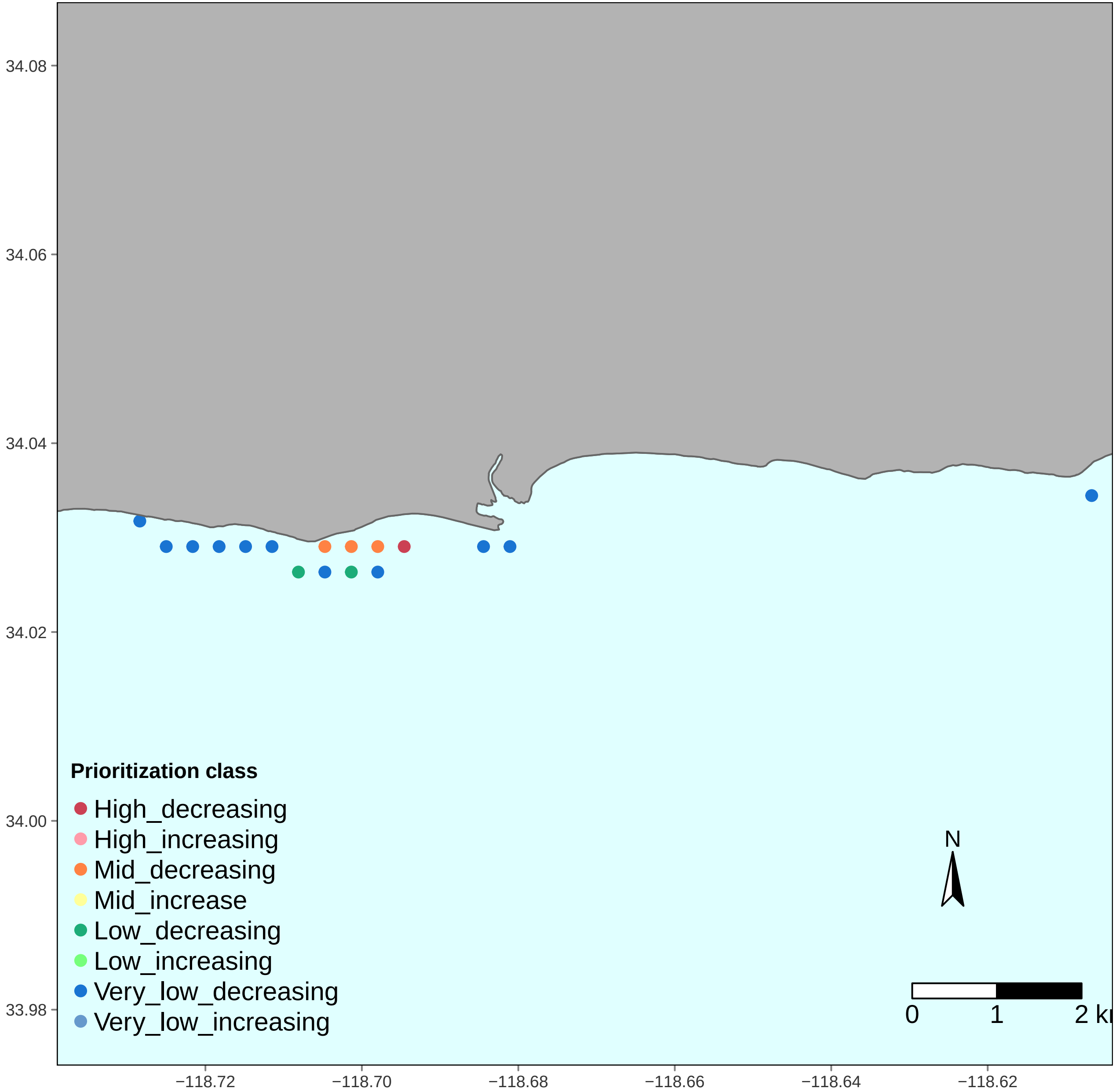

### SC classification_map 11

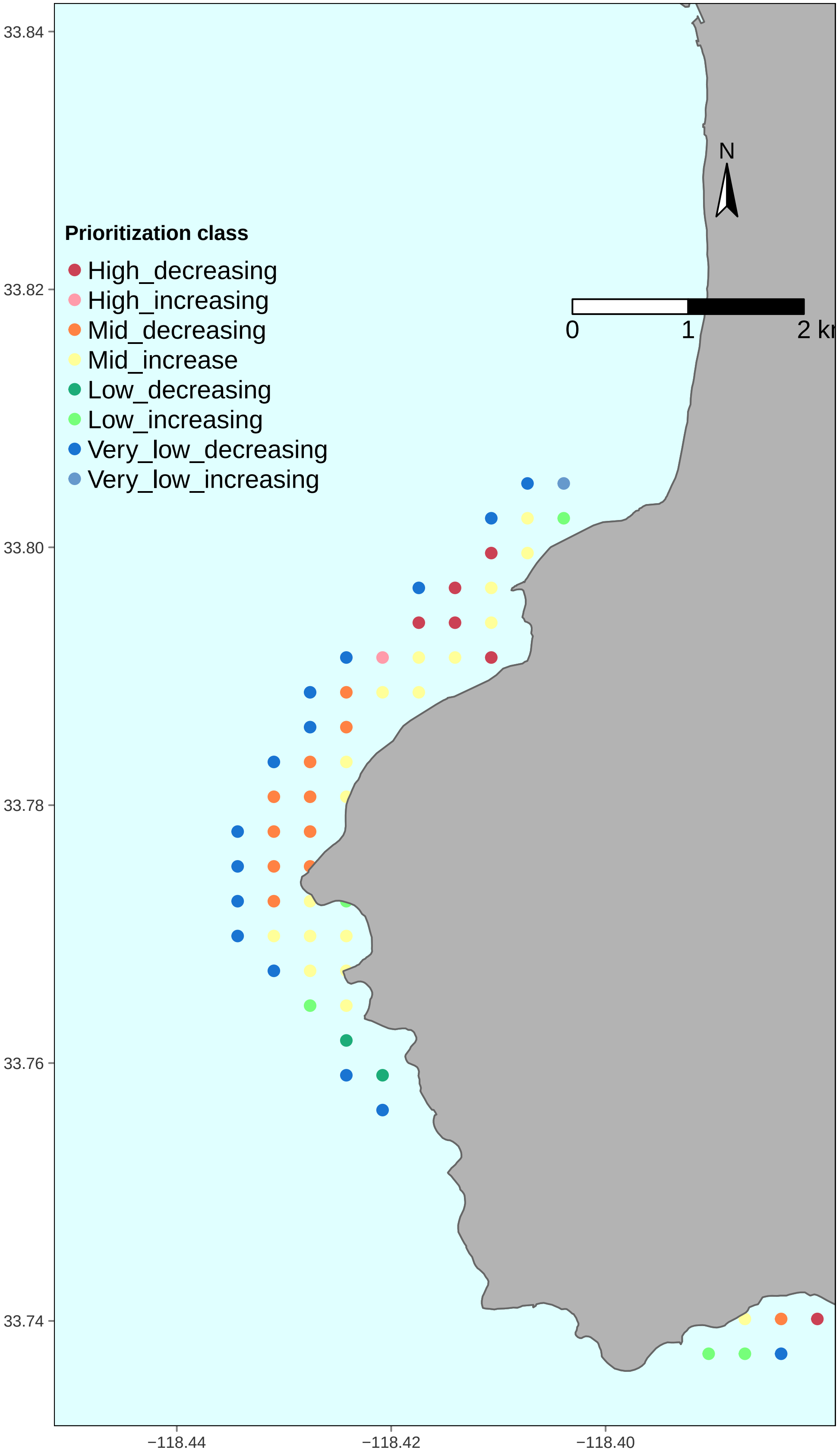

SC classification_map 12

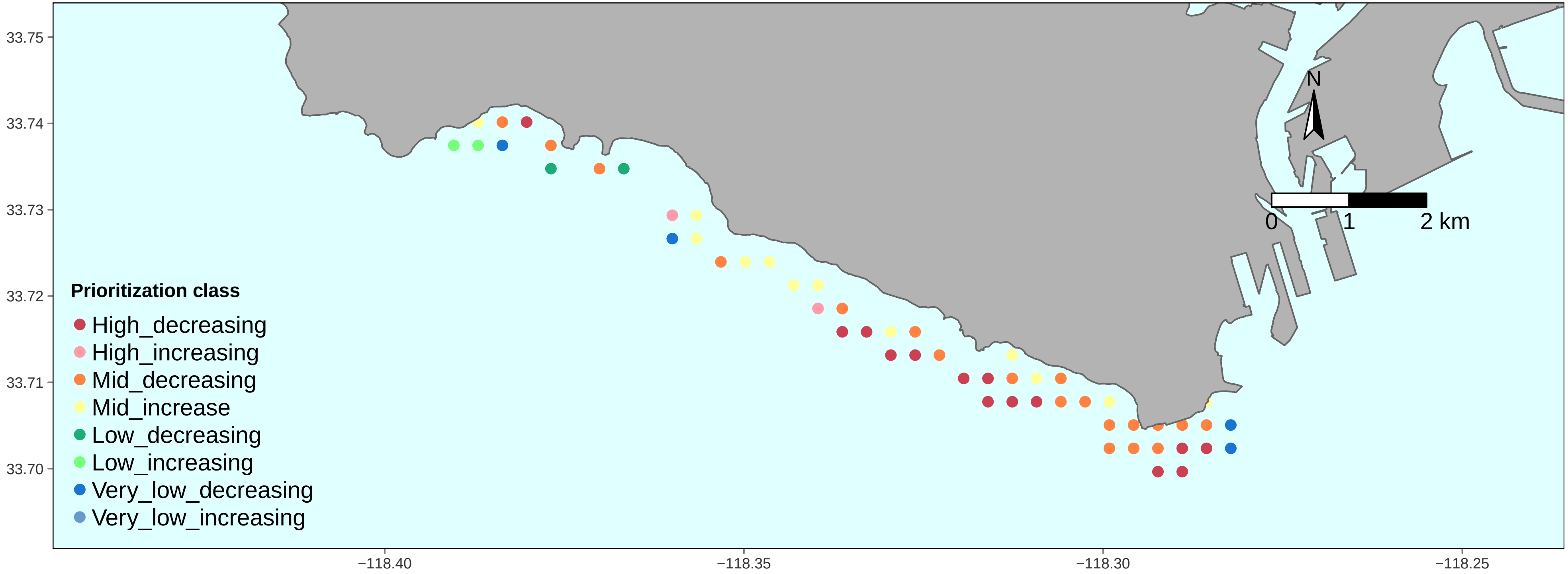

### SC classification_map 13

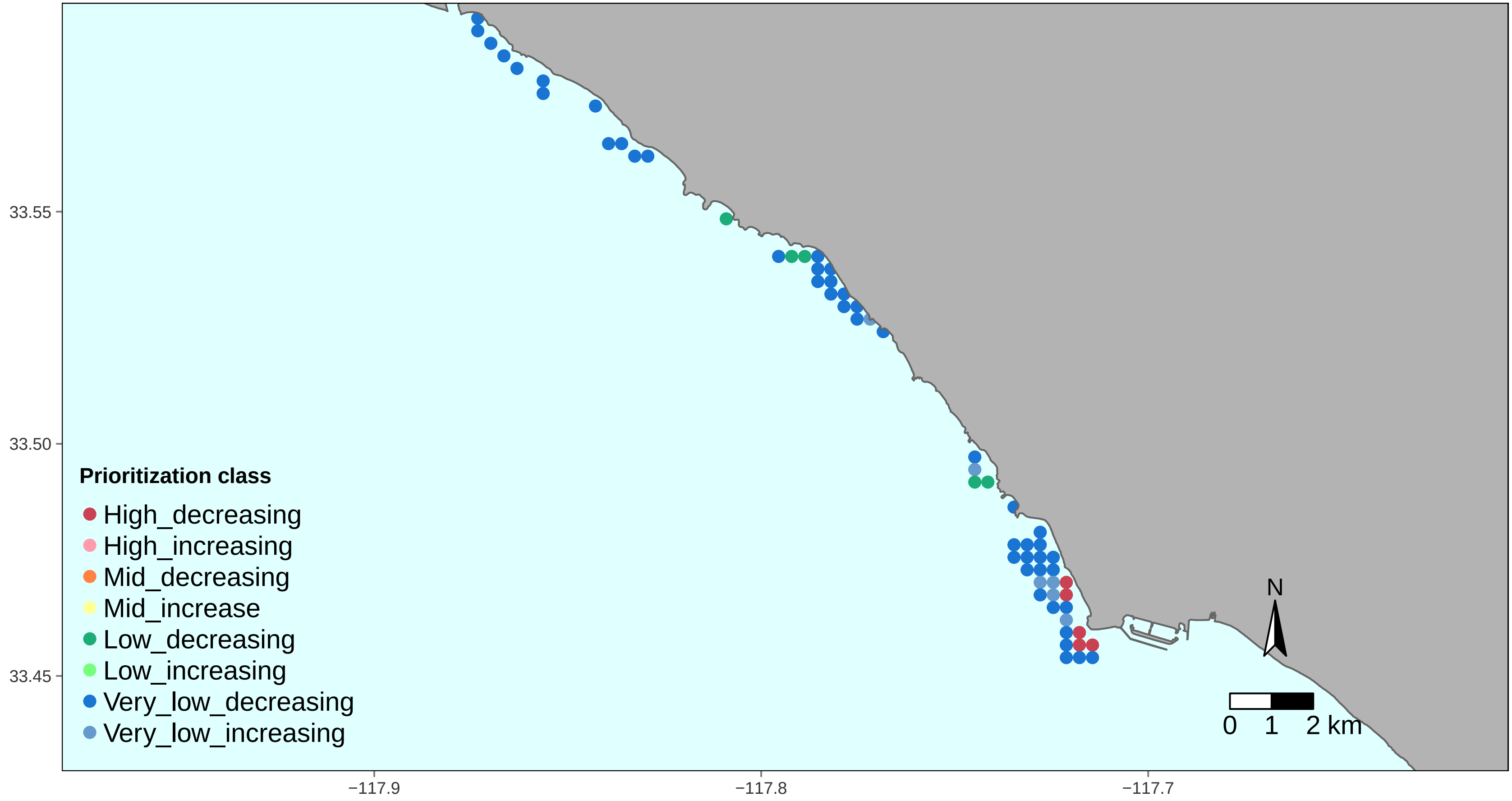

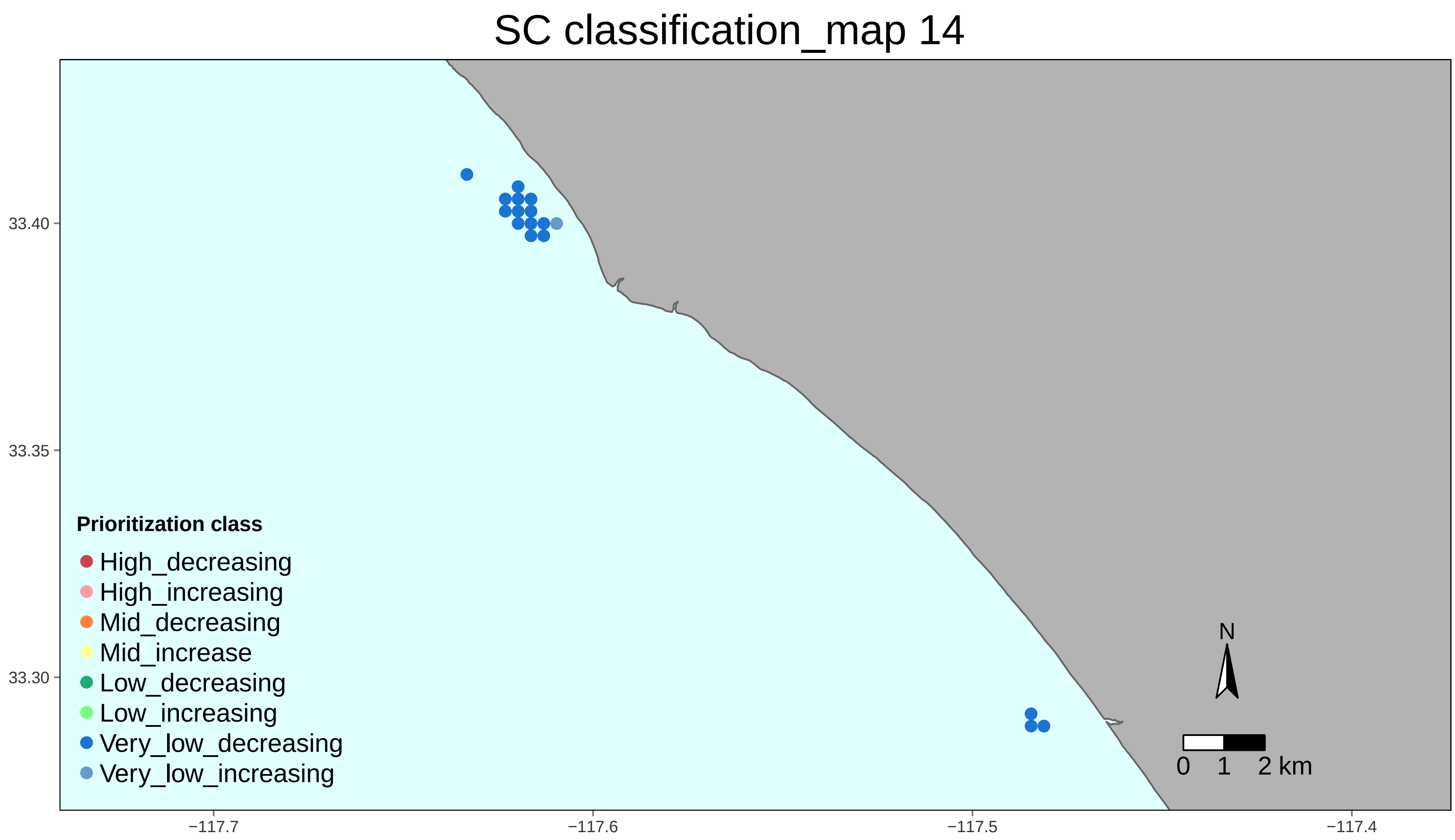

SC classification_map 14
33.40
33.35
33.30
−117.7
−117.6
−117.5
−117.4
Prioritization class
High_decreasing
High_increasing
Mid_decreasing
Mid_increase
Low_decreasing
Low_increasing
Very_low_decreasing
Very_low_increasing
N
0 1 2 km

### SC classification_map 15

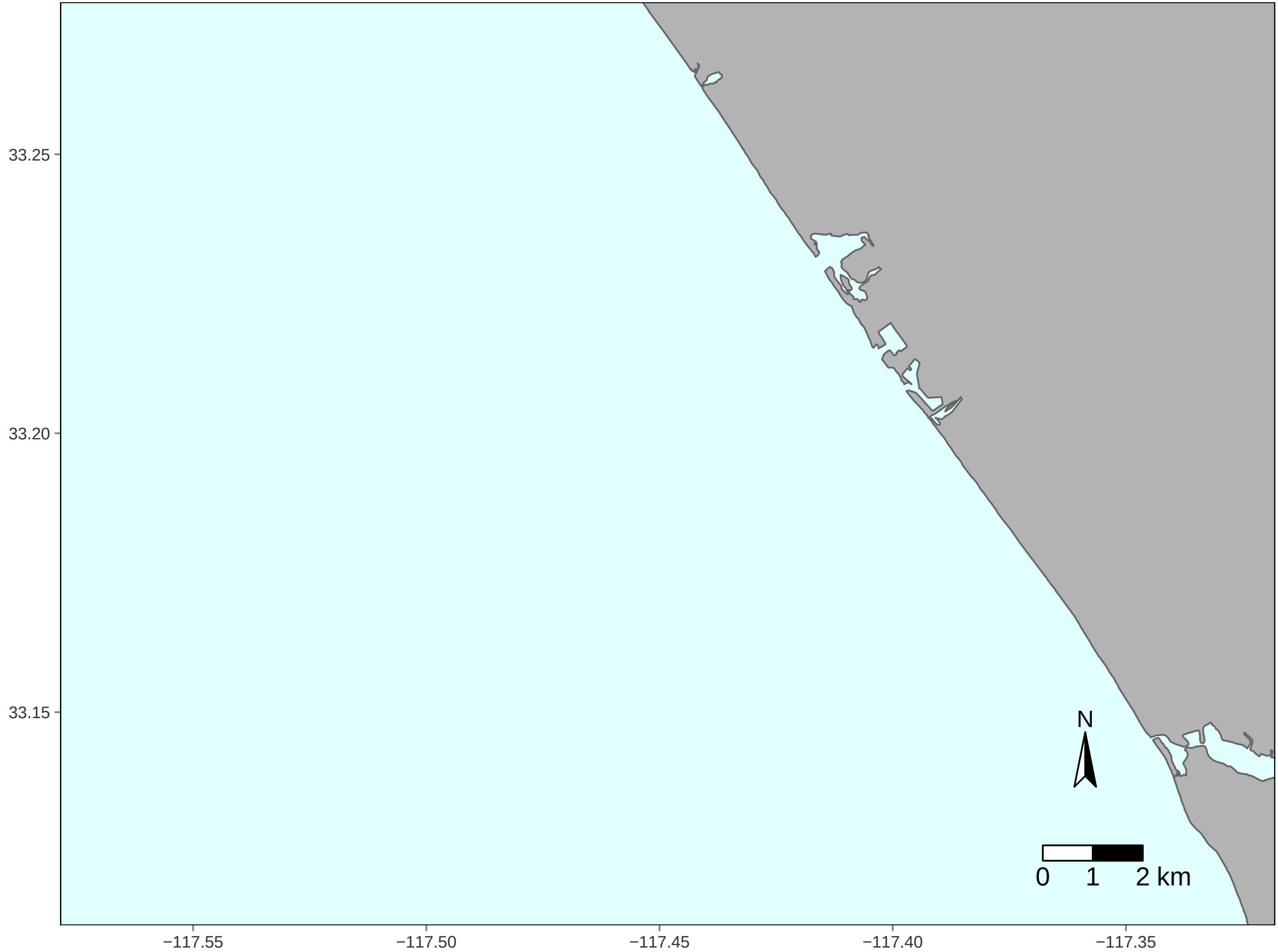

### SC classification_map 16

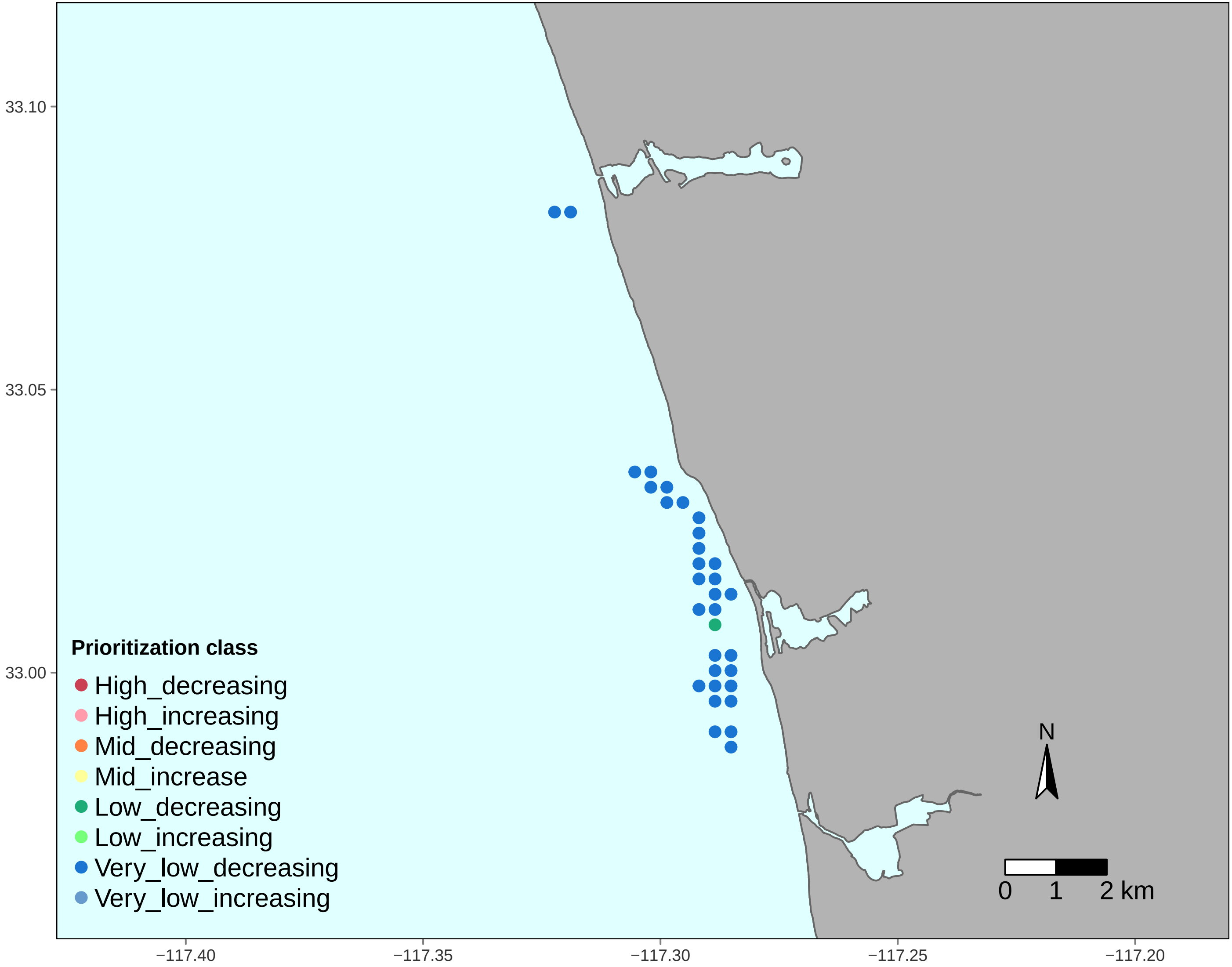

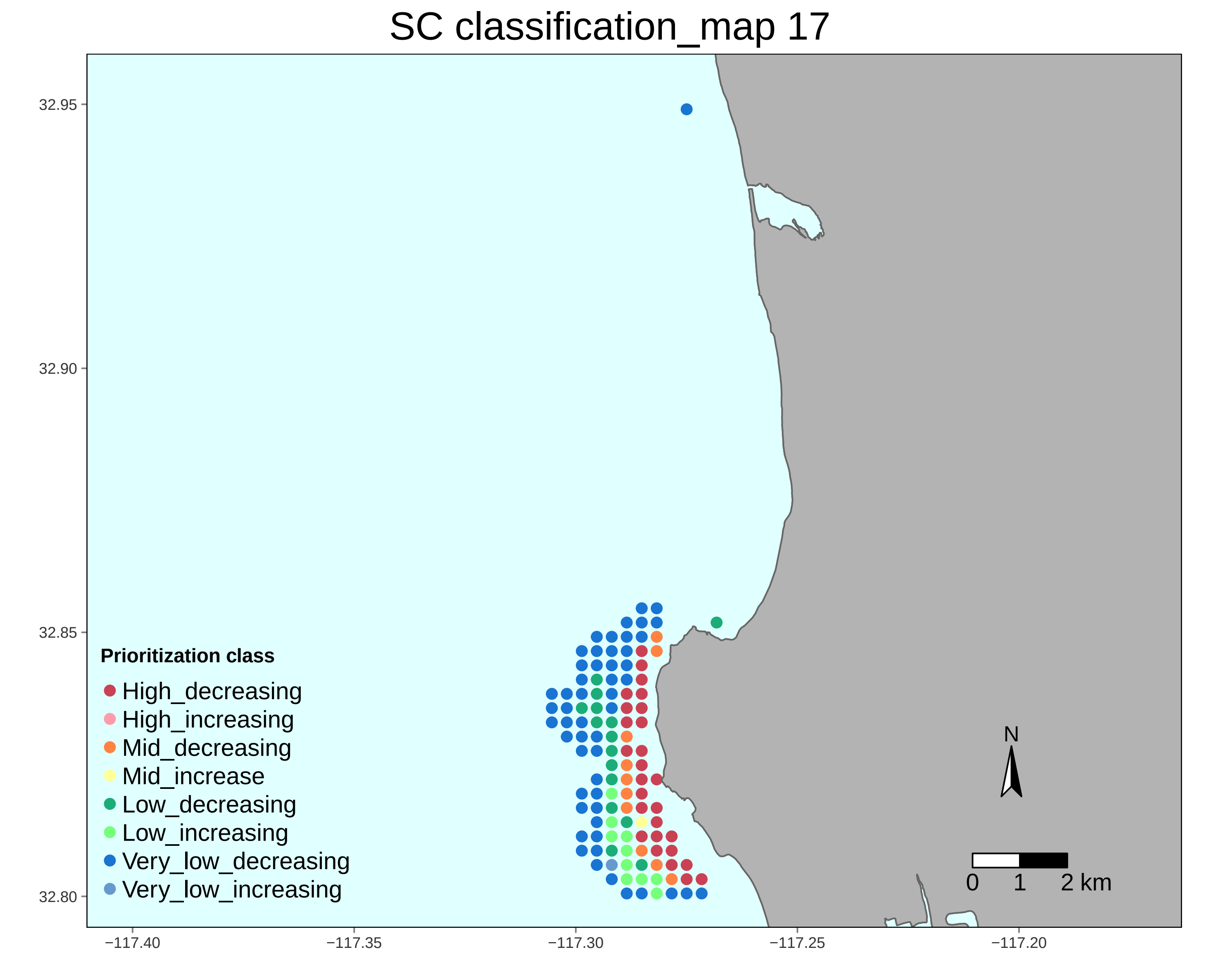
SC classification_map 17
32.95
32.90
32.85
32.80
−117.40
−117.35
−117.30
−117.25
−117.20
Prioritization class
High_decreasing
High_increasing
Mid_decreasing
Mid_increase
Low_decreasing
Low_increasing
Very_low_decreasing
Very_low_increasing
N
0
1
2 km

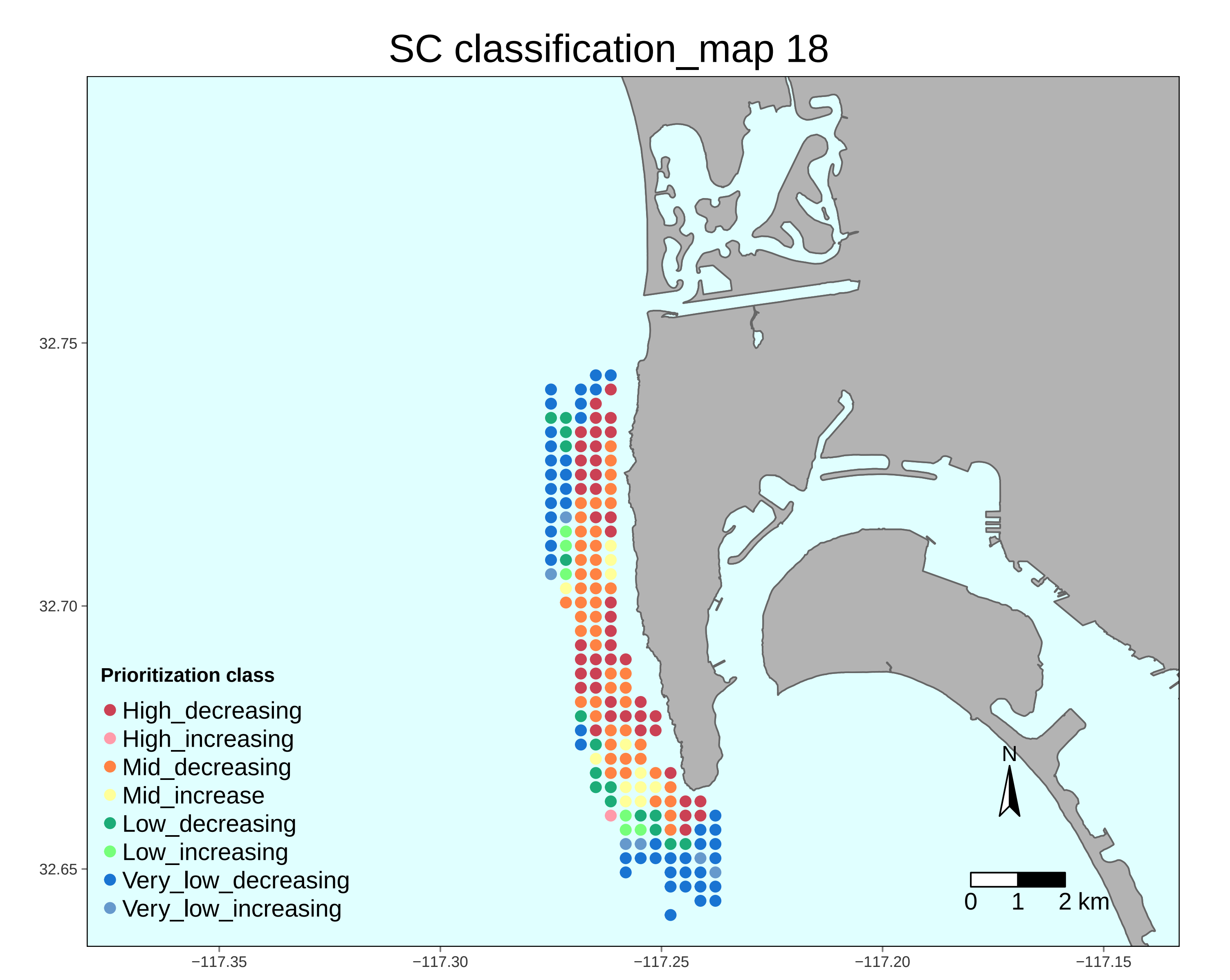
SC classification_map 18
32.75
32.70
32.65
−117.35
−117.30
−117.25
−117.20
−117.15
Prioritization class
High_decreasing
High_increasing
Mid_decreasing
Mid_increase
Low_decreasing
Low_increasing
Very_low_decreasing
Very_low_increasing
N
0
1
2 km

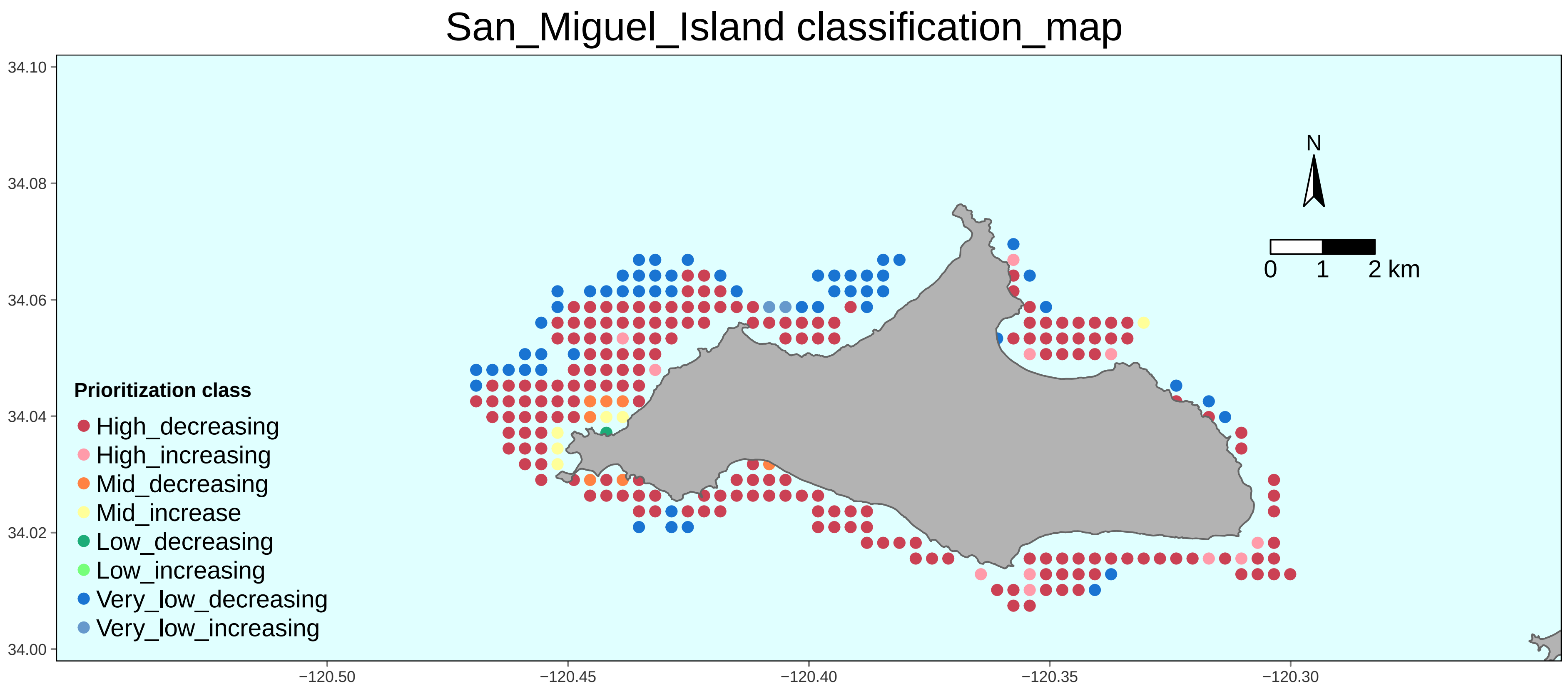

San_Miguel_Island classification_map
34.10
34.08
34.06
34.04
34.02
34.00
−120.50
−120.45
−120.40
−120.35
−120.30
N
0
1
2 km
Prioritization class
High_decreasing
High_increasing
Mid_decreasing
Mid_increase
Low_decreasing
Low_increasing
Very_low_decreasing
Very_low_increasing

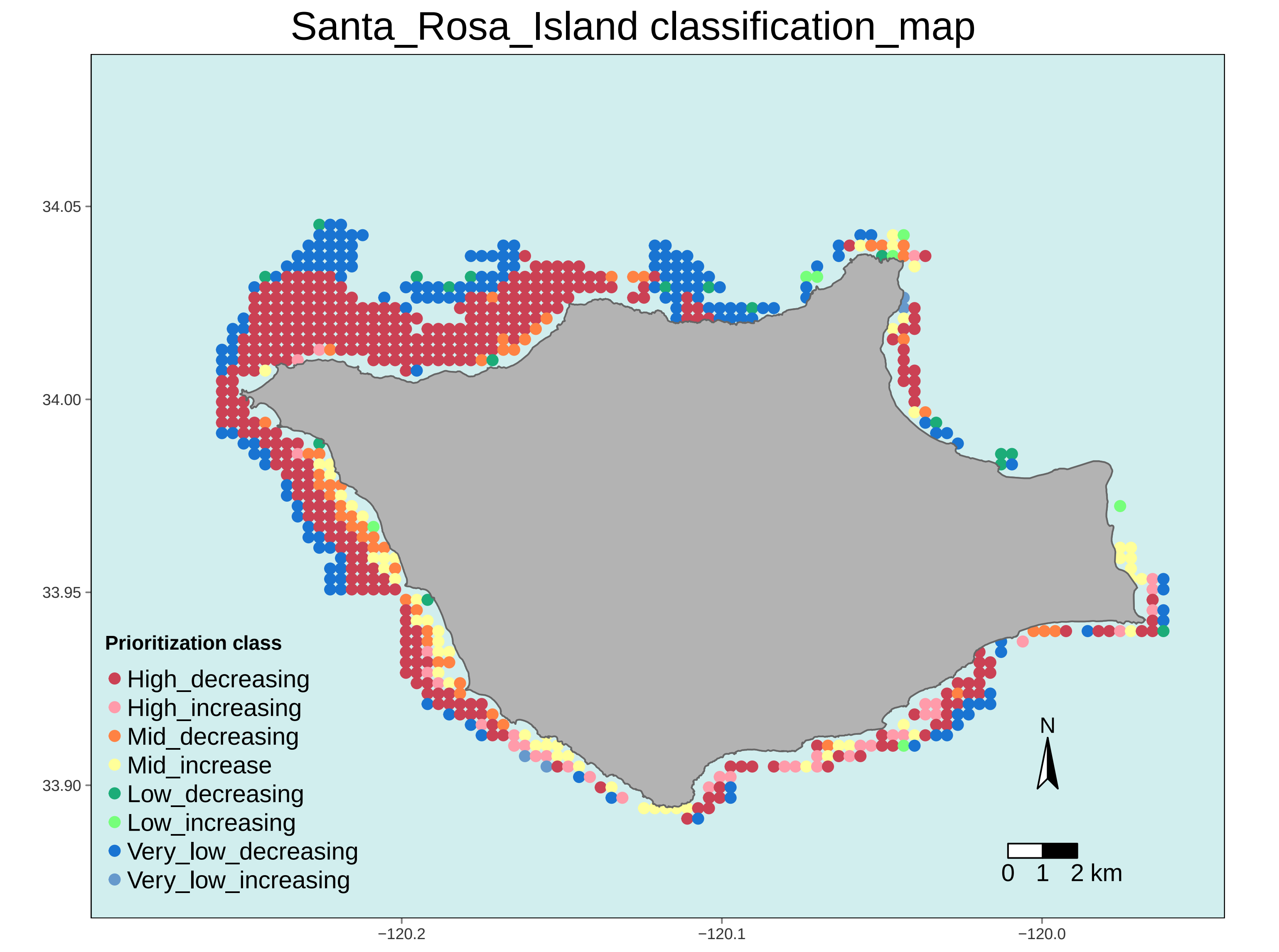
Santa_Rosa_Island classification_map
34.05
34.00
33.95
33.90
−120.2
−120.1
−120.0
Prioritization class
High_decreasing
High_increasing
Mid_decreasing
Mid_increase
Low_decreasing
Low_increasing
Very_low_decreasing
Very_low_increasing
N
0 1 2 km

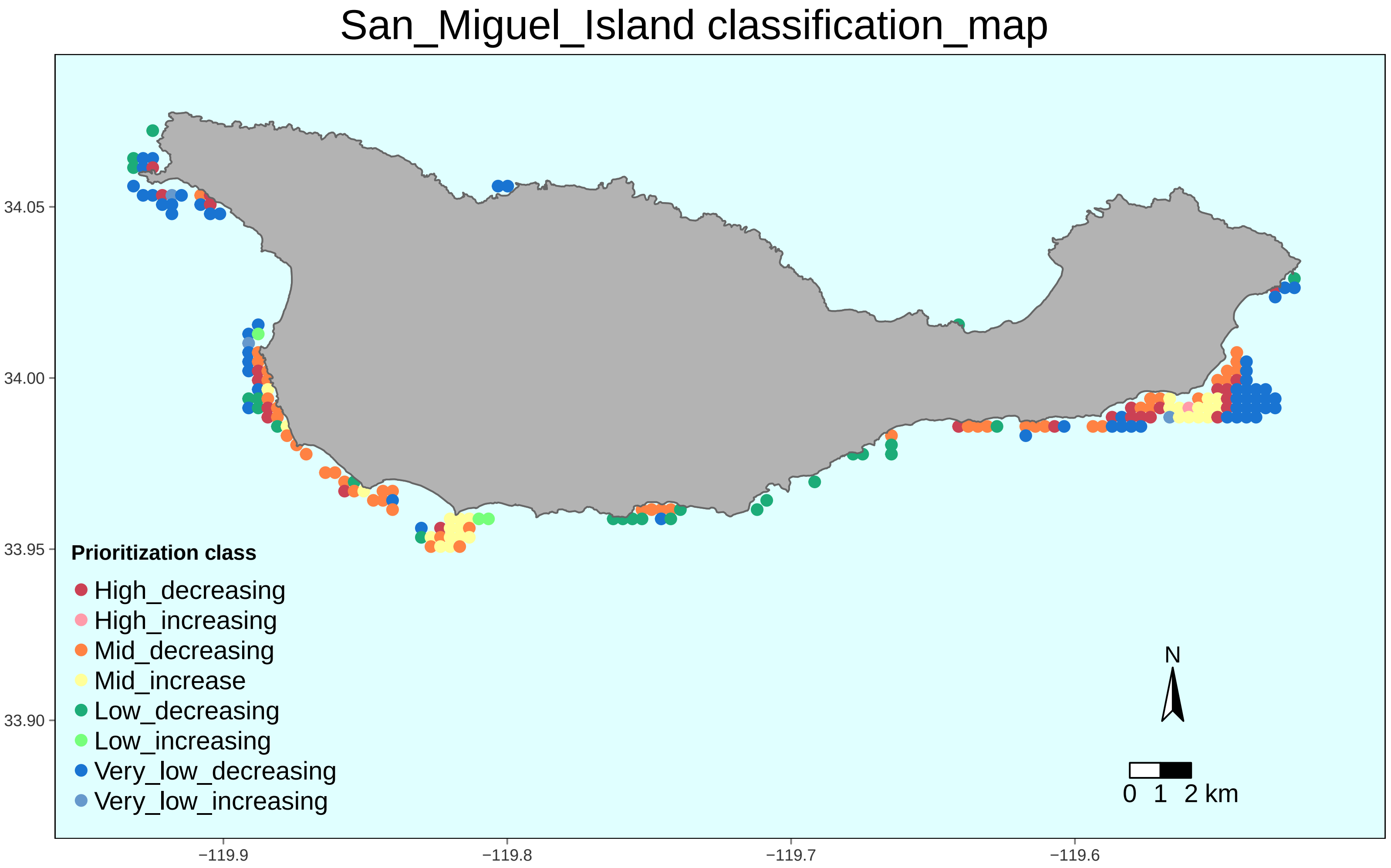
San_Miguel_Island classification_map
34.05
34.00
33.95
33.90
−119.9
−119.8
−119.7
−119.6
Prioritization class
High_decreasing
High_increasing
Mid_decreasing
Mid_increase
Low_decreasing
Low_increasing
Very_low_decreasing
Very_low_increasing
N
0 1 2 km

### Anacapa_Island classification_map

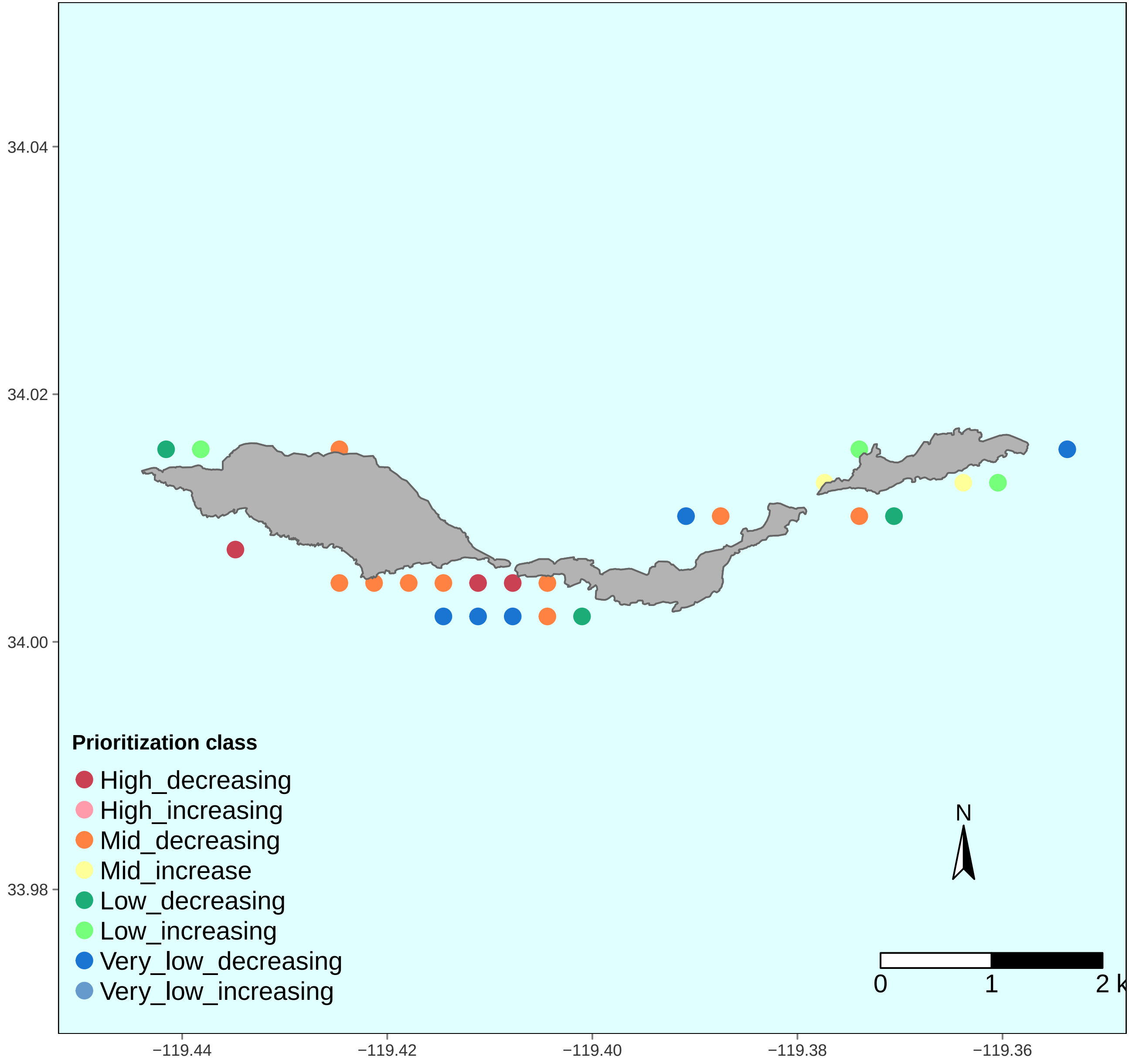

### Santa_Barbara_Island classification_map

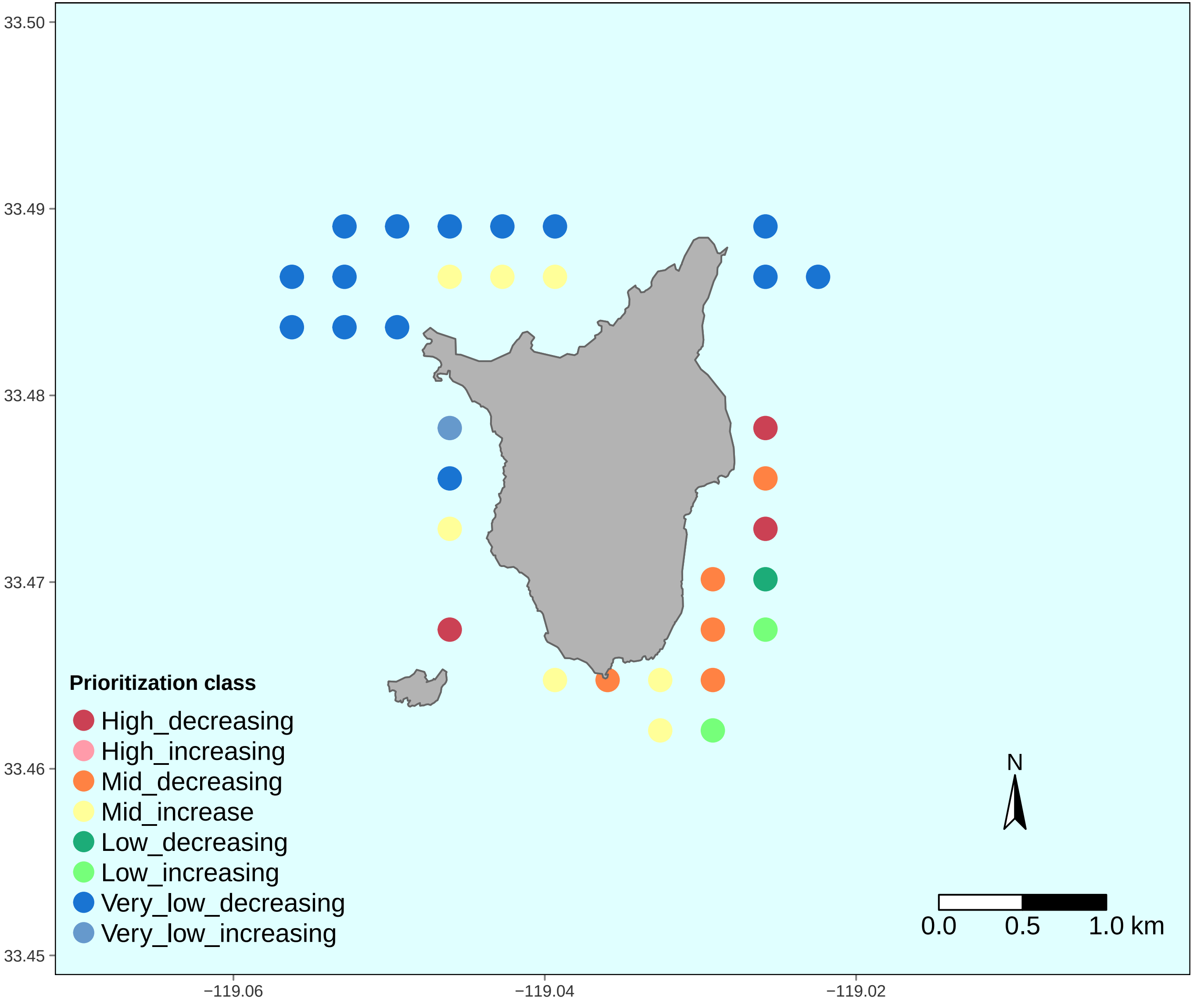

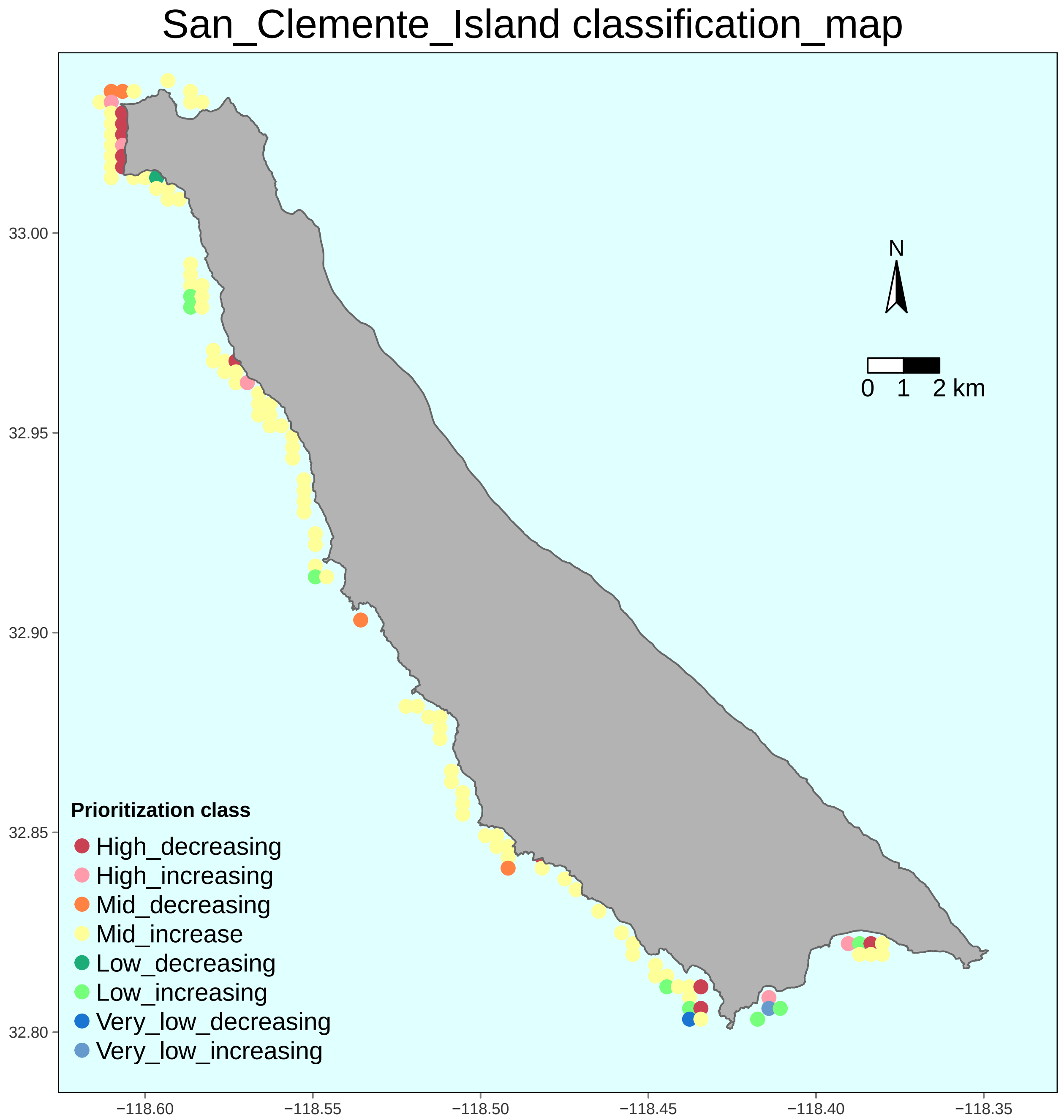
San_Clemente_Island classification_map
N
0 1 2 km
33.00
32.95
32.90
32.85
32.80
−118.60
−118.55
−118.50
−118.45
−118.40
−118.35
Prioritization class
High_decreasing
High_increasing
Mid_decreasing
Mid_increase
Low_decreasing
Low_increasing
Very_low_decreasing
Very_low_increasing

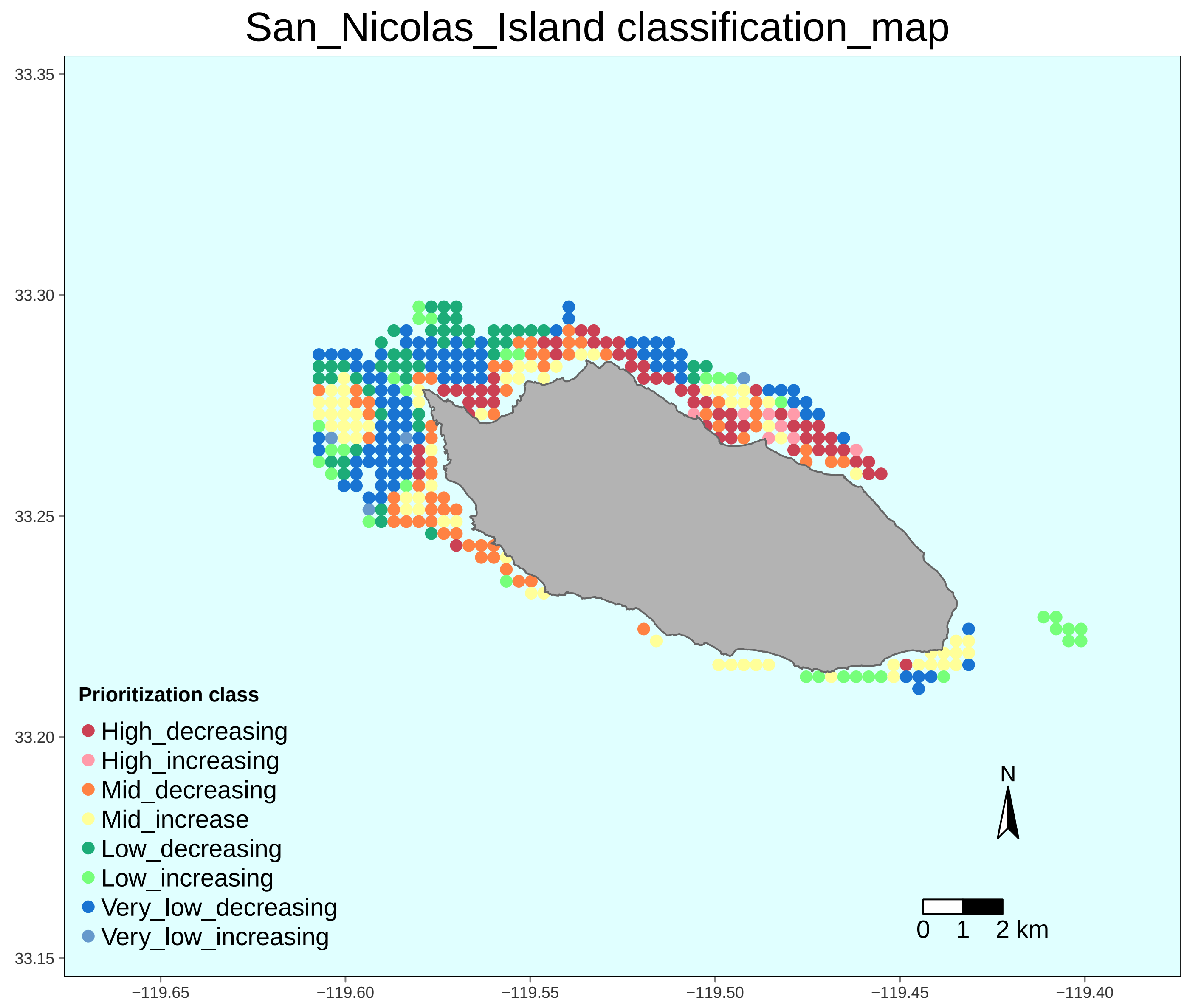
San_Nicolas_Island classification_map
33.35
33.30
33.25
33.20
33.15
−119.65
−119.60
−119.55
−119.50
−119.45
−119.40
Prioritization class
High_decreasing
High_increasing
Mid_decreasing
Mid_increase
Low_decreasing
Low_increasing
Very_low_decreasing
Very_low_increasing
N
0 1 2 km

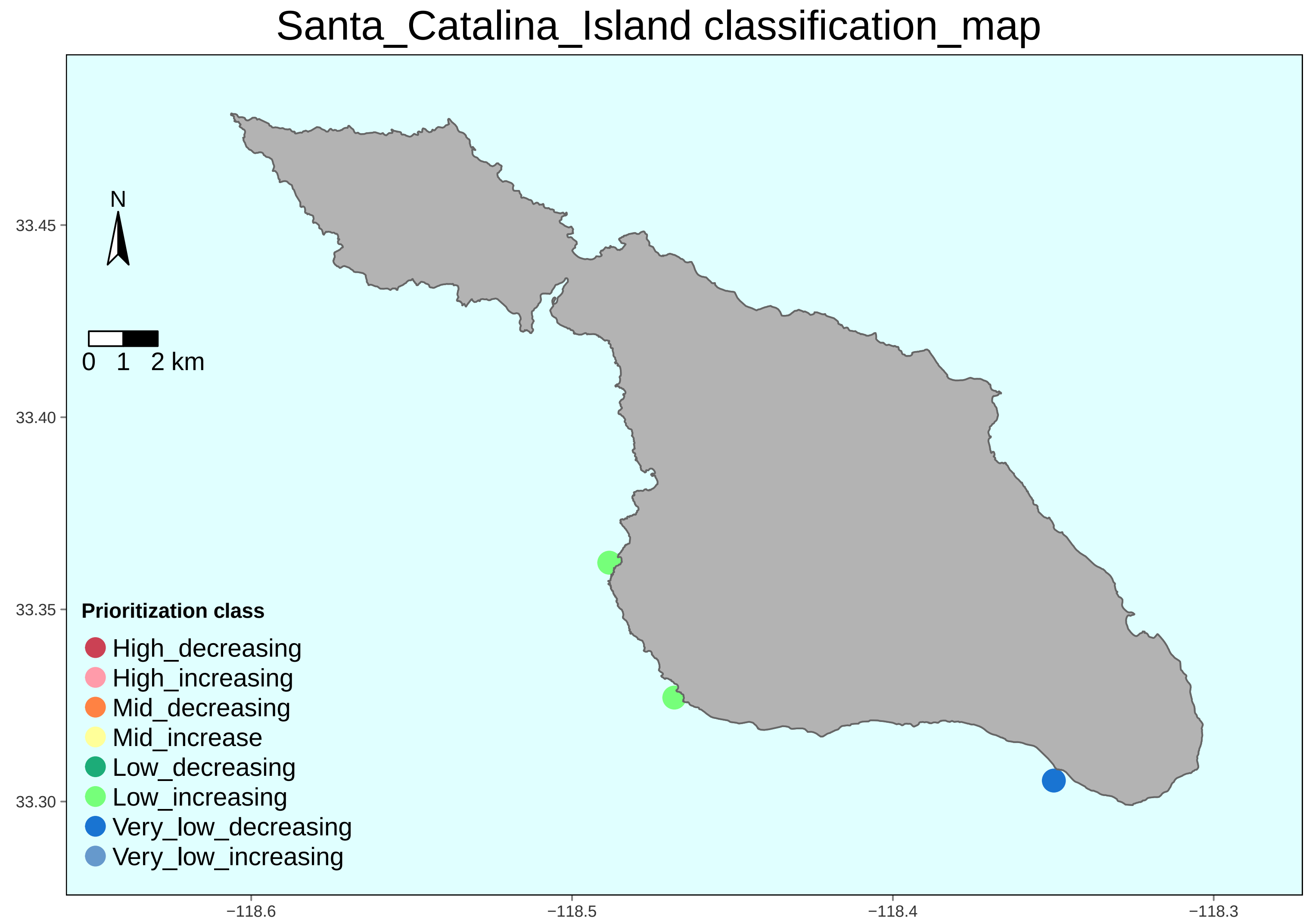

Santa_Catalina_Island classification_map
N
0 1 2 km
33.45
33.40
33.35
33.30
−118.6
−118.5
−118.4
−118.3
Prioritization class
High_decreasing
High_increasing
Mid_decreasing
Mid_increase
Low_decreasing
Low_increasing
Very_low_decreasing
Very_low_increasing
